# Novel artificial intelligence-based identification of drug-gene-disease interaction using protein-protein interaction

**DOI:** 10.1101/2024.06.01.596984

**Authors:** Y-h Taguchi, Turki Turki

## Abstract

The evaluation of drug-gene-disease interactions is key for the identification of drugs effective against disease. However, at present, drugs that are effective against genes that are critical for disease are difficult to identify. Following a disease-centric approach, there is a need to identify genes critical to disease function and find drugs that are effective against them. By contrast, following a drug-centric approach comprises identifying the genes targeted by drugs, and then the diseases in which the identified genes are critical. Both of these processes are complex. Using a gene-centric approach, whereby we identify genes that are effective against the disease and can be targeted by drugs, is much easier. However, how such sets of genes can be identified without specifying either the target diseases or drugs is not known. In this study, a novel artificial intelligence-based approach that employs unsupervised methods and identifies genes without specifying neither diseases nor drugs is presented. To evaluate its feasibility, we applied tensor decomposition (TD)-based unsupervised feature extraction (FE) to perform drug repositioning from protein-protein interactions (PPI) without any other information. Proteins selected by TD-based unsupervised FE include many genes related to cancers, as well as drugs that target the selected proteins. Thus, we were able to identify cancer drugs using only PPI. Because the selected proteins had more interactions, we replaced the selected proteins with hub proteins and found that hub proteins themselves could be used for drug repositioning. In contrast to hub proteins, which can only identify cancer drugs, TD-based unsupervised FE enables the identification of drugs for other diseases. In addition, TD-based unsupervised FE can be used to identify drugs that are effective in *in vivo* experiments, which is difficult when hub proteins are used. In conclusion, TD-based unsupervised FE is a useful tool for drug repositioning using only PPI without other information.

## 1 Introduction

The identification of drug-gene-disease information is critical for drug repositioning. Nevertheless, due to the fact that this does not involve a simple identification of paired information but rather of triplet information, identifying drug-gene-disease information is not an easy task. The identification of drug-gene-disease information is often performed in two steps. From the perspective of the genes (i.e. drug-centric approach), the identification of drug-gene relationships is performed first, followed by the extraction of gene-disease information by identifying diseases related to the selected genes. By contrast, when starting from the perspective of the diseases (i.e. disease-centric approach), the identification of gene-disease relations is performed first, followed by the extraction of drug-gene information via the identification of drugs that target the genes selected. Since the existence of a significant relationship at the second stage is not guaranteed (i.e., gene-disease information for the drug-centric approach or drug-gene information for the disease-centric approach), the identification of drug-gene-disease information is much more difficult than the identification of only gene-disease information or drug-gene information.

Several studies have reported on such drug-gene-disease relationships. Zickenrott et al. [1] attempted to predict disease–gene–drug relationships using differential network analysis, while Wong et al. [2] searched for gene-drug-disease interactions in pharmacogenomics by using GeneDive. Yu et al. [3] predicted drugs with opposing effects on disease genes using a directed network, Sun [4] investigated gene-gene, drug-drug, and disease-disease networks and studied their relationships, and Qahwaji et al. [5] reviewed the genetic approaches to drug development and therapy. Furthermore, Iida et al. [6] investigated network-based characterization of disease–disease relationships in terms of drugs and therapeutic targets. Lastly, Quan et al. [7] considered genetic disease genes as promising sources of drug targets. While these are only some examples, all focused on diseases, drugs, or both, and no studies have focused on genes without considering drugs and diseases. Thus, the relevant literature is not free from the use of a two-stage approach.

To address this difficulty, linear algebra has often been used to identify drug-genedisease information, since it enables the identification of drug-gene-disease information in neither a drug-centric nor disease-centric centric-manner. Wang et al. [8] applied matrix factorization to gene expression matrices for drug and disease treatments, while Kim and Cho [9] employed tensor decomposition (TD) to extract drug-gene-disease information starting from the product of ID embedding vectors for drugs, genes, and diseases. In these approaches, because the identification of drug-gene-disease information is performed without the need for a two-stage approach, these approaches can often avoid the difficulty associated with this approach.

Another advantage of these linear-algebra-based approaches is that they are fully unsupervised. Therein, there is no need for any drug- or disease-specific information (e.g., differential expression between healthy controls and patients), and we are free to identify drug-gene-disease information in a fully data-driven manner. We recently applied a TD-based unsupervised FE [10, 11] to integrate PPI with gene expression in cancer [12]. We found that integrated analysis enhanced the coincidence with clinical labels. In this study, we also found that PPI only contained information related to cancers, even though PPI themselves are unlikely to be associated with cancers. In this paper, we attempt to determine the degree of the relationship between PPI and cancers using TD-based unsupervised FE without integrating other information (e.g. gene expression). As a result, we found that applying TD-based unsupervised FE to PPI enabled the identification of cancer-related genes that were also used for drug repositioning. After identifying that the selected proteins were likely to be hub proteins in PPI, we replaced the proteins selected via TD-based unsupervised FE with hub proteins and found that the hub proteins could be used for drug repositioning. The distinction between hub proteins and those selected using TD-based unsupervised FE was the strength of the correlation among the interactions; that is, proteins selected using TD-based unsupervised FE were found to exhibit a greater number of shared interactions. Only proteins selected using TD-based unsupervised FE had hits in the *in vivo*-based drug database DrugMatrix. The identification of proteins that not only have more interactions (i.e., hub proteins) but also more shared interactions may be the key to identifying more promising proteins for drug repositioning.

## 2. Results

Figure 1 illustrates the analysis conducted in this study.

**Fig. 1.**
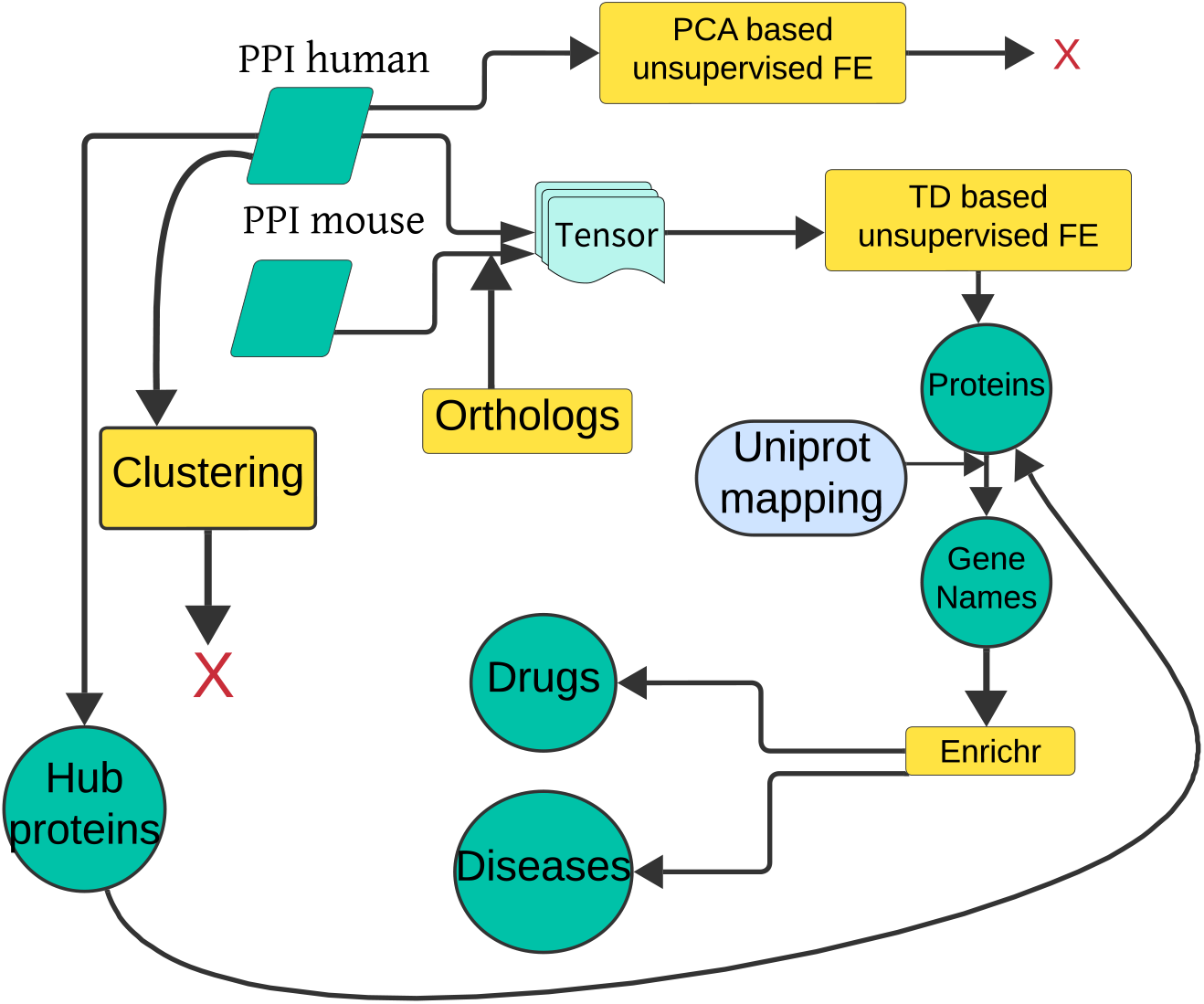
Flowchart of the analyses in this study. PCA-based unsupervised FE was applied to human PPI, but failed. TD-based unsupervised FE was applied to tensor generated from human and mouse PPI. Gene names associated with the identified proteins were uploaded to Enrichr to identify associated diseases and drugs. Hub proteins and proteins selected via cluster analyses were tested and used for the identification of diseases and drugs.

### 2.1 TD-based unsupervised FE

The first matrix to be integrated was human PPI, 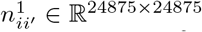 (for BioGRID) or 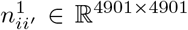, and the second was mouse PPI, 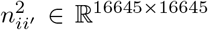 (for BioGRID) or 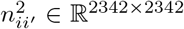 (for DIP). Since the number of common protein, *N*common, is 9688 (for BioGRID) or 1097 (for DIP), and the number of orthogonal protein, *N*ortho, is 4026 (for BioGRID) or and 432 (for DIP), the resulting integrated tensor is *n*_*ii*_*′*_*k*_ ∈ ℝ^*N ×N ×*2^ where *N* = 24875 + 16645 – 9688 – 4026 = 27806 (for BioGRID) or *N* = 4901 + 2342 – 1097 – 432 = 5714 (for DIP). After applying HOSVD to *n*_*ii*_*′*_*k*_, we obtained eq. (7). *P*_*i*_s were attributed to *i*s using*u*_2*i*_ (for BioGRID and DIP) or *u*_3*i*_ (only for DIP) using Eq. (6) with replacing *u*_*𝓁i*_ with 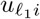 and are corrected using the BH criterion [10, 11] (The reason why *u*_3*i*_ was considered only for DIP was because genes selected by *u*_3*i*_ for DIP share the same enriched diseases, cancers, with those by *u*_2*i*_, which was not the case for BioGRID. For more details, see the latter part of this paper.) Thus, 195 (using *u*_2*i*_ for BioGRID) and 196 (using *u*_2*i*_ for DIP) and 59 (using *u*_3*i*_ for DIP) proteins were associated with adjusted *P* -values less than 0.01. The Uniprot accession numbers associated with these proteins were converted to 217 (using *u*_2*i*_ for BioGRID), 193 (using *u*_2*i*_ for DIP), and 57 (using *u*_3*i*_ for DIP) gene names by Uniprot ID mapping (see Supplementary Information for the list of proteins and gene names). 217, 193, and 57 gene names were uploaded to Enrichr [13] for evaluation purposes.

Next, the types of diseases associated with three sets of gene names were identified. Firstly, we considered the category of “Jensen Diseases”.

For the 217 gene names selected by *u*_2*i*_ on BioGrid (Table S1 and Fig. 2), not only were there highly significant diseases, but most were cancers or tumors (“cancer,” “stomach cancer,” “adenoma,” “immune system cancer,” “ovarian cancer,” “ductal carcinoma in situ,” “esophageal carcinoma,” and “lymphoid leukemia”).

**Fig. 2.**
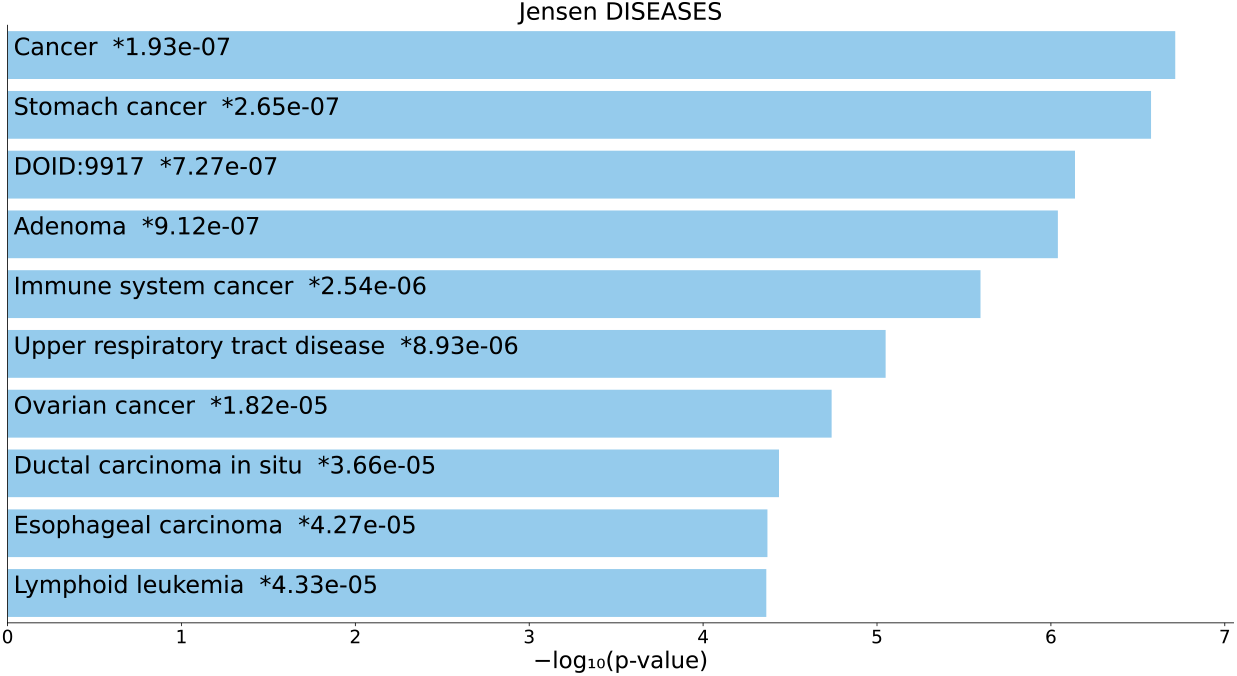
Top 10 diseases in the “Jensen Diseases” category of Enrichr for 217 gene names selected by *u*_2*i*_ for BioGrid. Blue: *P <* 0.05, *: adjusted *P <* 0.05.

For the 193 gene names selected by *u*_2*i*_ for the DIP (see Table S2 and Fig. 3), not only were there highly significant diseases, but more than half were cancers or tumors (“lymphoid leukemia,” “immune system cancer,” “cancer,” “biliary tract cancer,” and “ovarian cancer”). Additionally, “familial adenomatous polyposis” is known to develop into cancer [14]. For the 57 gene names selected by *u*_2*i*_ for DIP (Table S3 and Fig. 4), not only were there highly significant diseases, but more than half were cancer or tumors (“ovarian cancer,” “immune system cancer,” “breast cancer,” “biliary tract cancer,” “ductal carcinoma in situ,” and “hereditary breast ovarian cancer”). In addition, “Li-Fraumeni syndrome” is known to be the cause of several cancers [15].

**Fig. 3.**
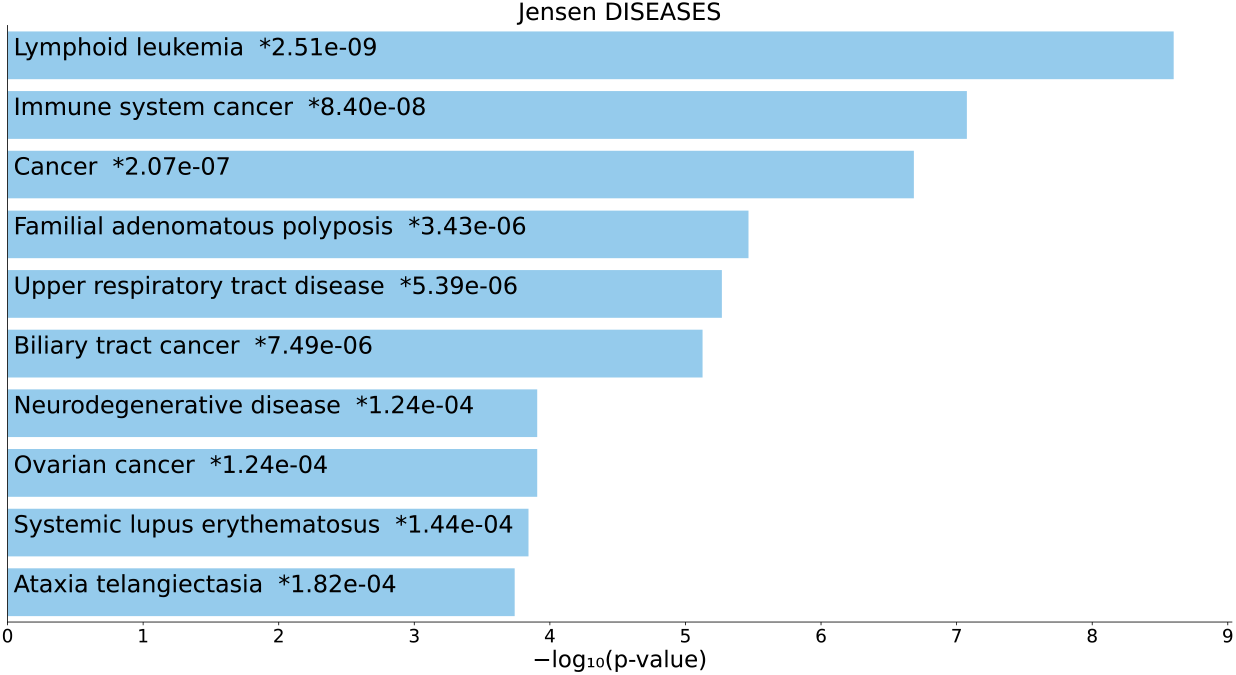
Top 10 diseases in the “Jensen Diseases” category of Enrichr for 193 gene names selected by *u*_2*i*_ for DIP. Blue: *P <* 0.05, *: adjusted *P <* 0.05.

**Fig. 4.**
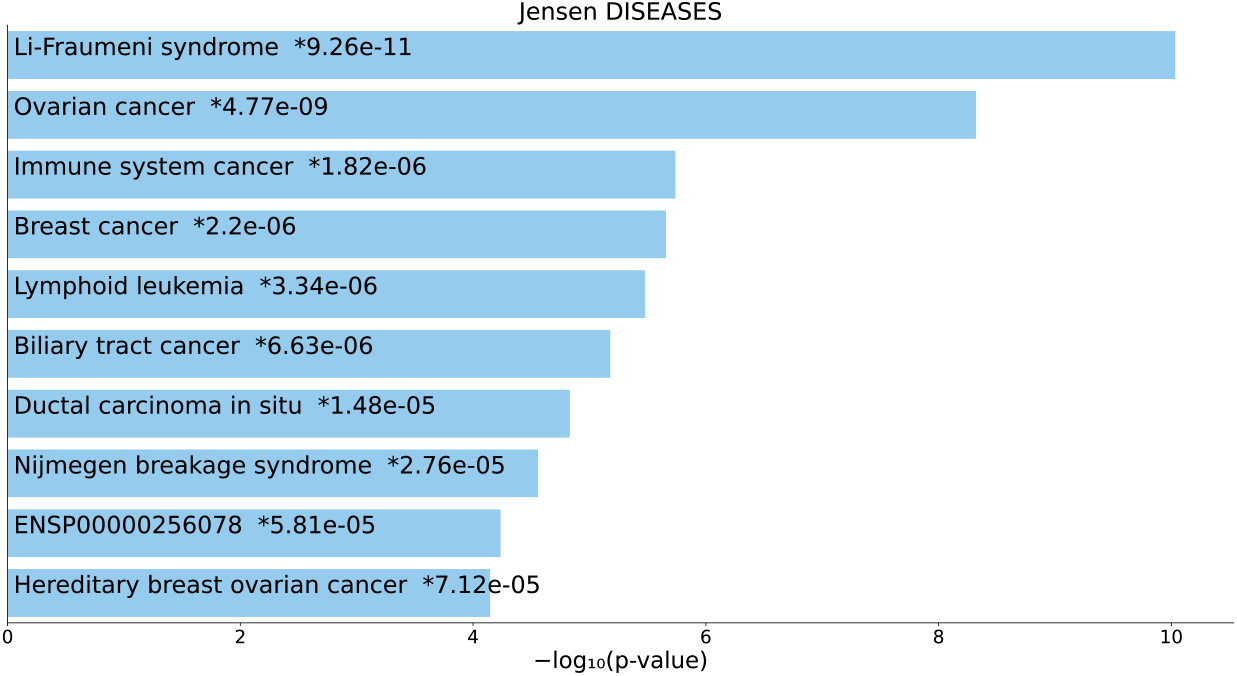
Top 10 diseases in the “Jensen Diseases” category of Enrichr for 57 gene names selected by *u*_3*i*_ for DIP. Blue: *P <* 0.05, *: adjusted *P <* 0.05.

Next, we considered the “OMIM Diseases” category of Enrichr. For the 217 gene names selected by *u*_2*i*_ for BioGrid (Table S4 and Fig. 5), six diseases were cancer or tumors (“breast cancer,” “ovarian cancer,” “thyroid carcinoma,” “prostate cancer,” “colorectal cancer,” “pancreatic cancer,” and “gastric cancer”), although statistical (i.e., associated with adjusted *P* -values less than 0.05) significance was only observed for the top two. In contrast to BioGRID, which failed to identify sufficiently large significant diseases, 193 gene names selected by *u*_2*i*_ for DIP (Table S5 and Fig. 6), not only were there highly significant diseases, but four diseases were related to cancers or tumors (“colorectal cancer,” “breast cancer,” “leukemia,” and “lymphoma”). Additionally, “Fanconi anemia” is also often associated with cancer [16]. For 57 gene names selected by *u*_3*i*_ for DIP (Table S6 and Fig. 7), not only there were highly significant diseases, but also more than half of the diseases were cancers or tumors (“pancreatic cancer,” “ovarian cancer,” “colorectal cancer,” “breast cancer,” “prostate cancer,” and “melanoma”). In addition, “Fanconi anemia” was also identified. In conclusion, these three sets of proteins were found to be highly related to cancers and tumors regardless of the PPI datasets used, excluding the 217 gene names selected by *u*_2*i*_ for BioGRID. Next, we considered potential drug repositioning using these gene sets. Multiple categories can be used for drug repositioning in Enrichr: “LINCS L1000 Chem Pert Consensus Sigs,” “DSigDB,” “DrugMatrix,” “Drug Perturbations from GEO down,” and “Drug Perturbations from GEO up.” All of these categories return a list of compounds that are supposed to significantly target a set of uploaded genes. Because three sets of gene names are supposed to be deeply related to cancers and tumors, drugs that target these genes can be used to treat tumors and cancers. A list of the top 10 compounds in the “LINCS L1000 Chem Pert Consensus Sigs” category for 217 gene names selected by *u*_2*i*_ for BioGrid and 193 gene names selected by *u*_2*i*_ for DIP (no significant hit for 57 gene names selected by *u*_3*i*_ for DIP is provided in Tables S7 and S8 and Figs. 8 and 9). Tables S9, S10, and S11 and Figs. 10, 11, and 12 list the top 10 compounds in the “DSigDB” category for 217 gene names selected by *u*_2*i*_ for BioGRID, 193 gene names selected by *u*_2*i*_ for DIP, and 57 gene names selected by *u*_3*i*_ for DIP. Tables S12 and S13 and Figs. 13 and 14 list the top 10 compounds in the “DrugMatrix” category for 217 gene names selected by *u*_2*i*_ for BioGRID and 193 gene names selected by *u*_2*i*_ for DIP (no significant hits for 57 gene names selected by *u*_3*i*_ for DIP). Tables S14, S15, and S16 and Figs. 15, 16, and 17 list the top 10 compounds in the “Drug Perturbations from GEO down” category for 217 gene names selected by *u*_2*i*_ for BioGRID, 193 gene names selected by *u*_2*i*_ for DIP, and 57 gene names selected by *u*_3*i*_ for DIP. Tables S17, S18, and S19 and Figs. 18, 19, and 20 list the top 10 compounds in the “Drug Perturbations from GEO up” category for 217 gene names selected by *u*_2*i*_ for BioGRID, 193 gene names selected by *u*_2*i*_ for DIP, and 57 gene names selected by *u*_3*i*_ for DIP.

**Fig. 5.**
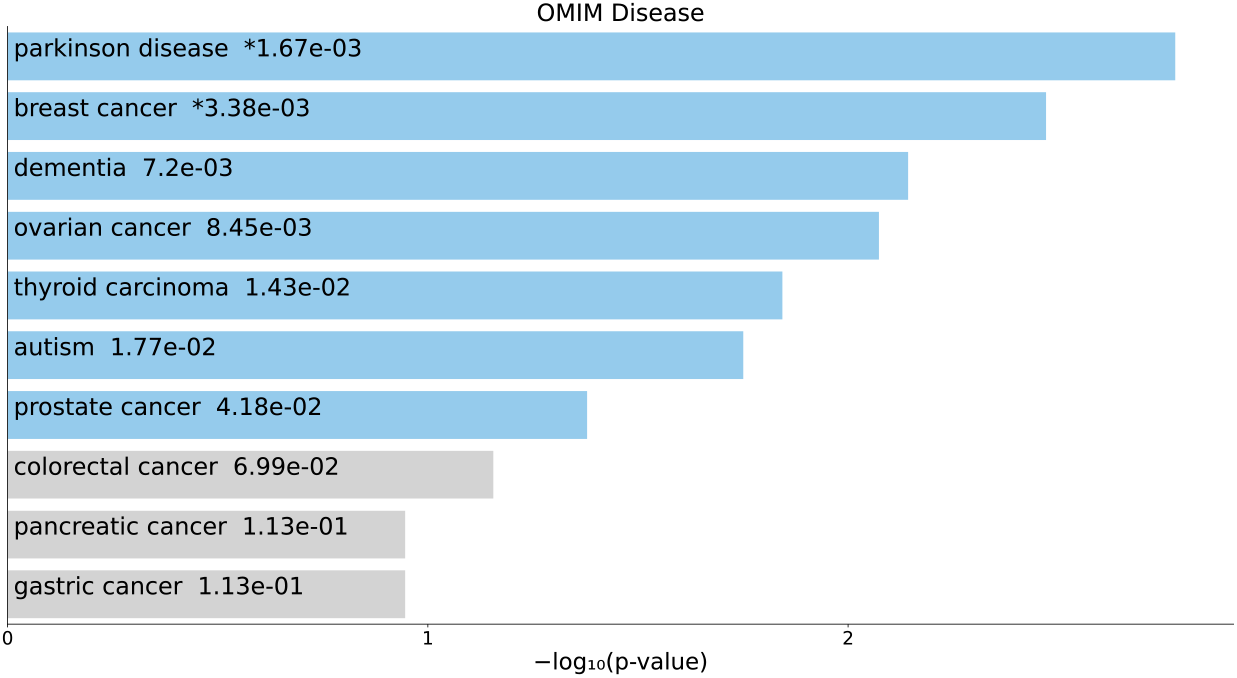
Top 10 diseases in the “OMIM Diseases” category of Enrichr for 217 gene names selected by *u*_2*i*_ for BioGrid. Blue: *P <* 0.05, *: adjusted *P <* 0.05.

**Fig. 6.**
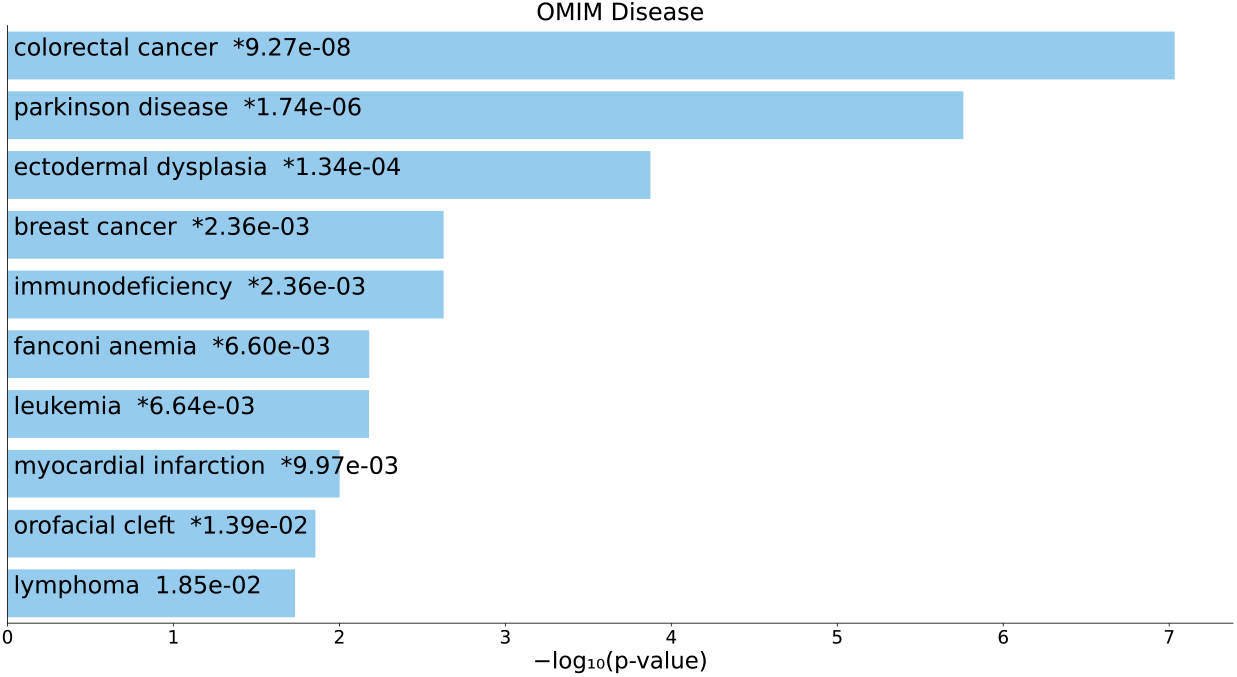
Top 10 diseases in the “OMIM Diseases” category of Enrichr for 193 gene names selected by *u*_2*i*_ for DIP. Blue: *P <* 0.05, *: adjusted *P <* 0.05.

**Fig. 7.**
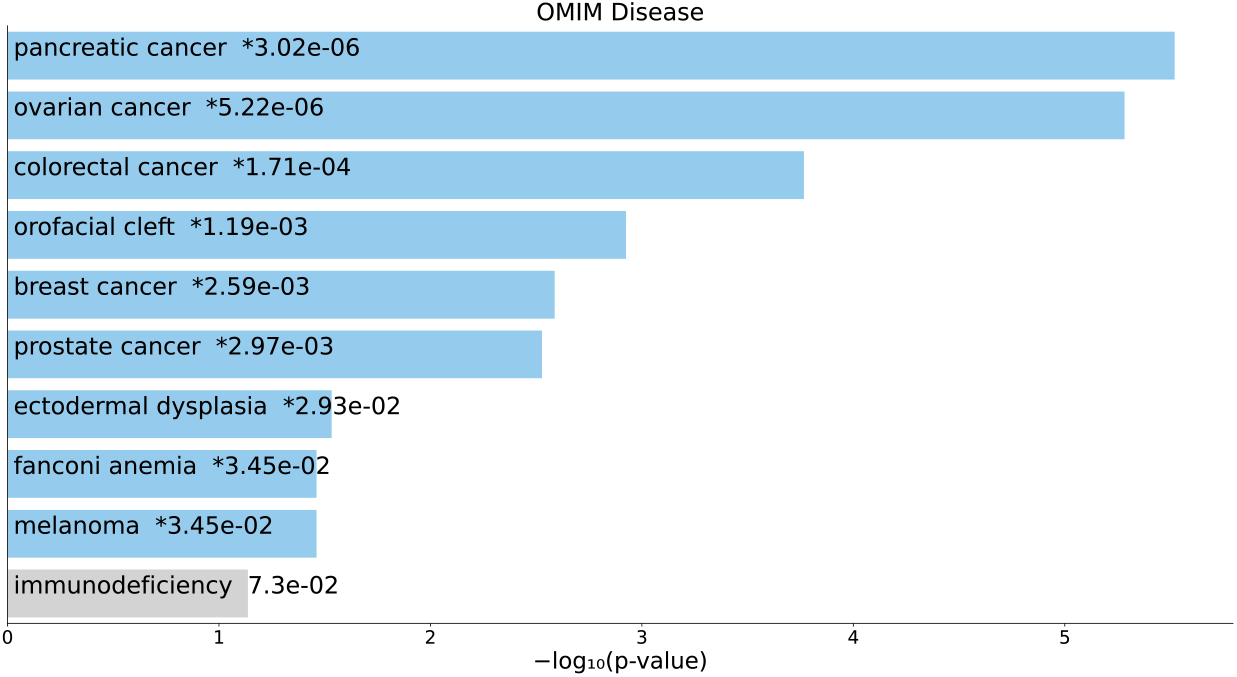
Top 10 diseases in the “OMIM Diseases” category of Enrichr for 57 gene names selected by *u*_3*i*_ for DIP. Blue: *P <* 0.05, *: adjusted *P <* 0.05.

**Fig. 8.**
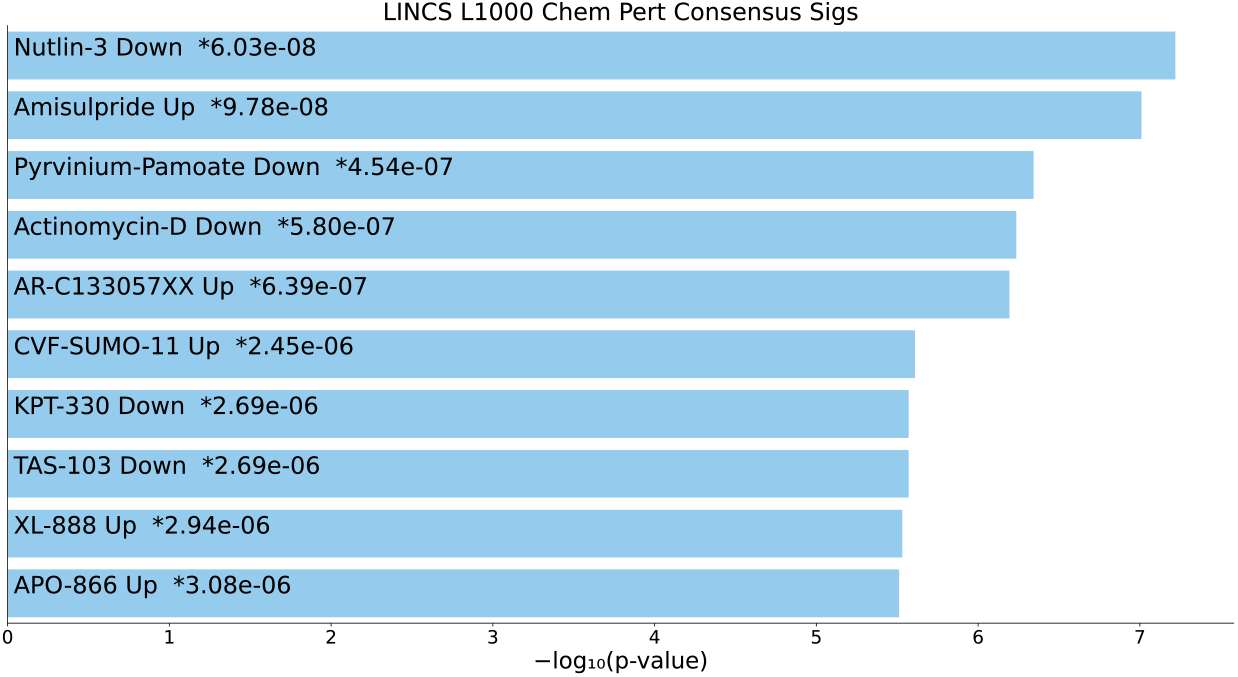
Top 10 drugs in the “LINCS L1000 Chem Pert Consensus Sigs” category of Enrichr for 217 gene names selected by *u*_2*i*_ for BioGrid. Blue: *P <* 0.05, *: adjusted *P <* 0.05.

**Fig. 9.**
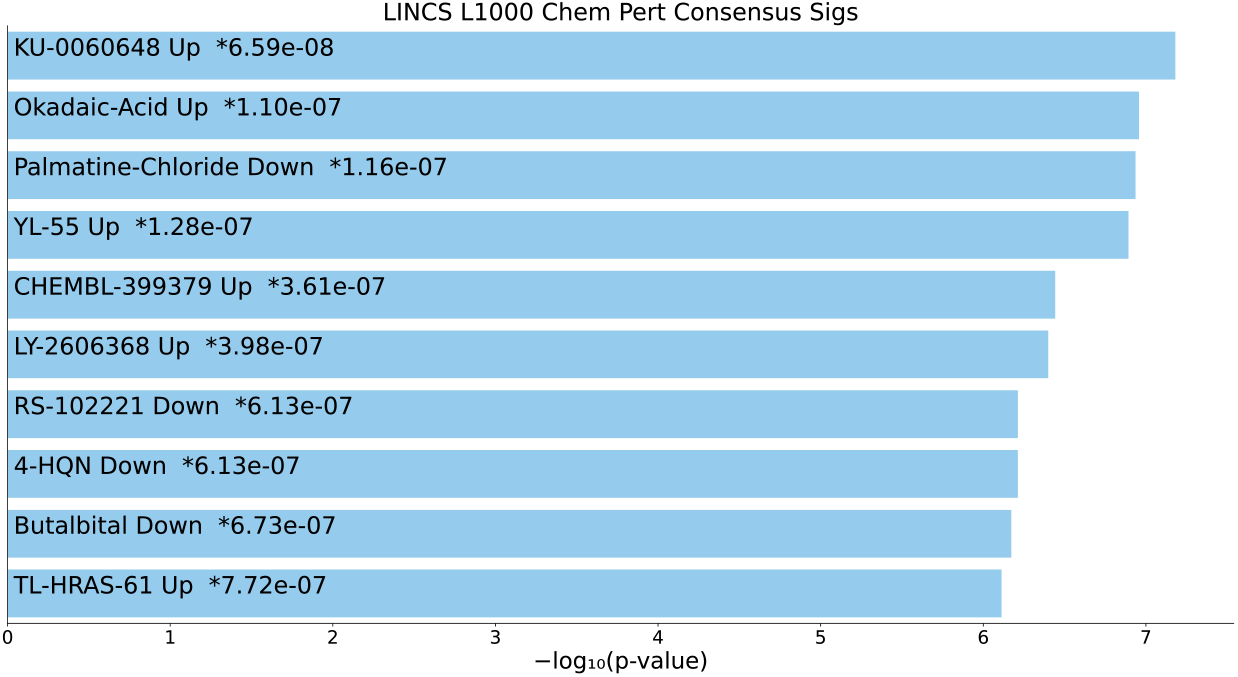
Top 10 drugs in the “LINCS L1000 Chem Pert Consensus Sigs” category of Enrichr for 193 gene names selected by *u*_2*i*_ for DIP. Blue: *P <* 0.05, *: adjusted *P <* 0.05.

**Fig. 10.**
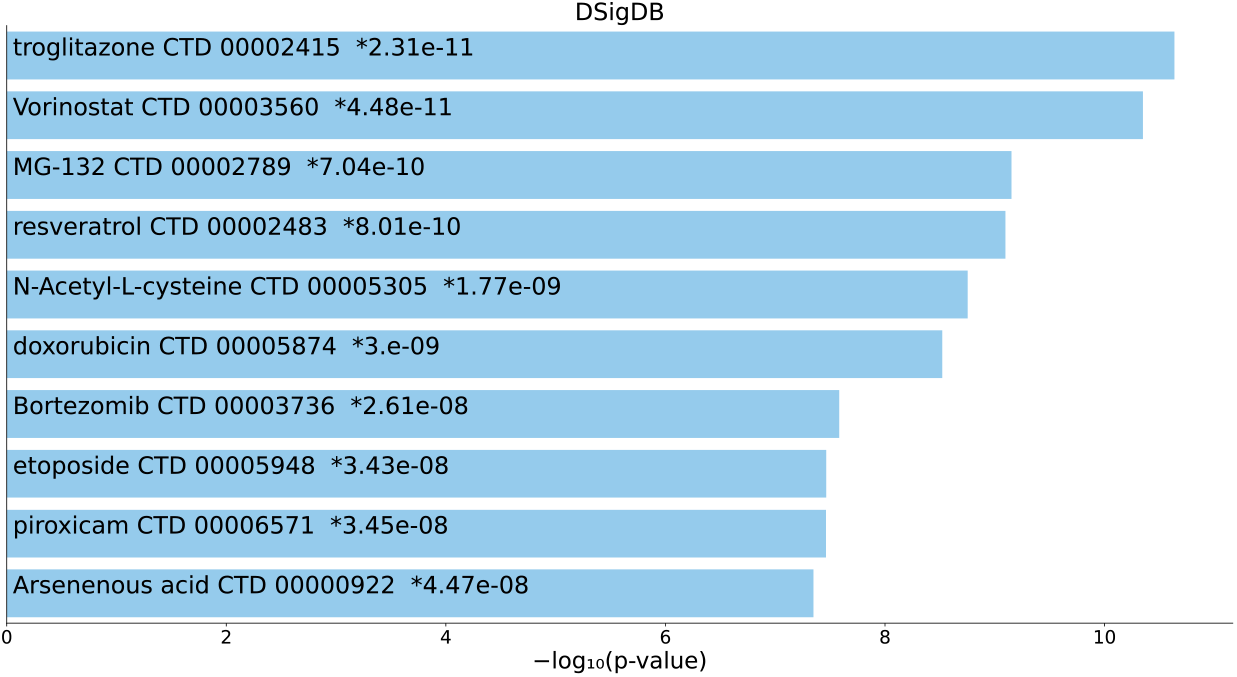
Top 10 drugs in the “DSigDB” category of Enrichr for 217 gene names selected by *u*_2*i*_ for BioGrid. Blue: *P <* 0.05, *: adjusted *P <* 0.05.

**Fig. 11.**
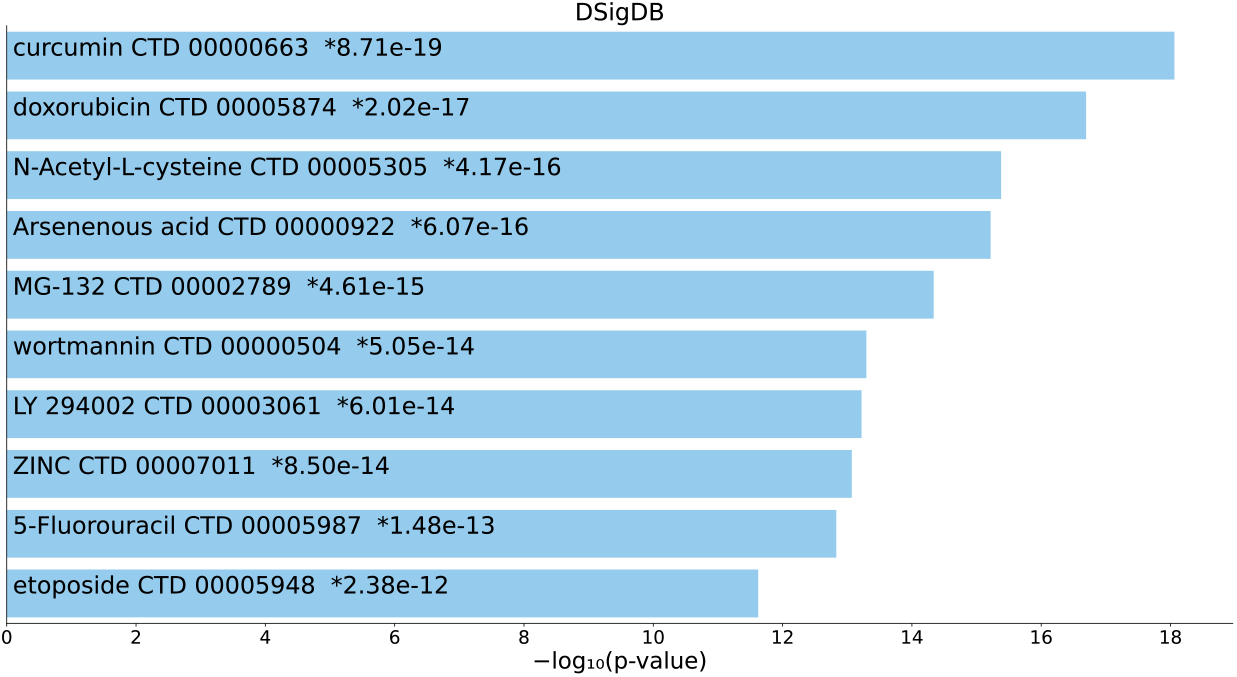
Top 10 drugs in the “DSigDB” category of Enrichr for 193 gene names selected by *u*_2*i*_ for DIP. Blue: *P <* 0.05, *: adjusted *P <* 0.05.

**Fig. 12.**
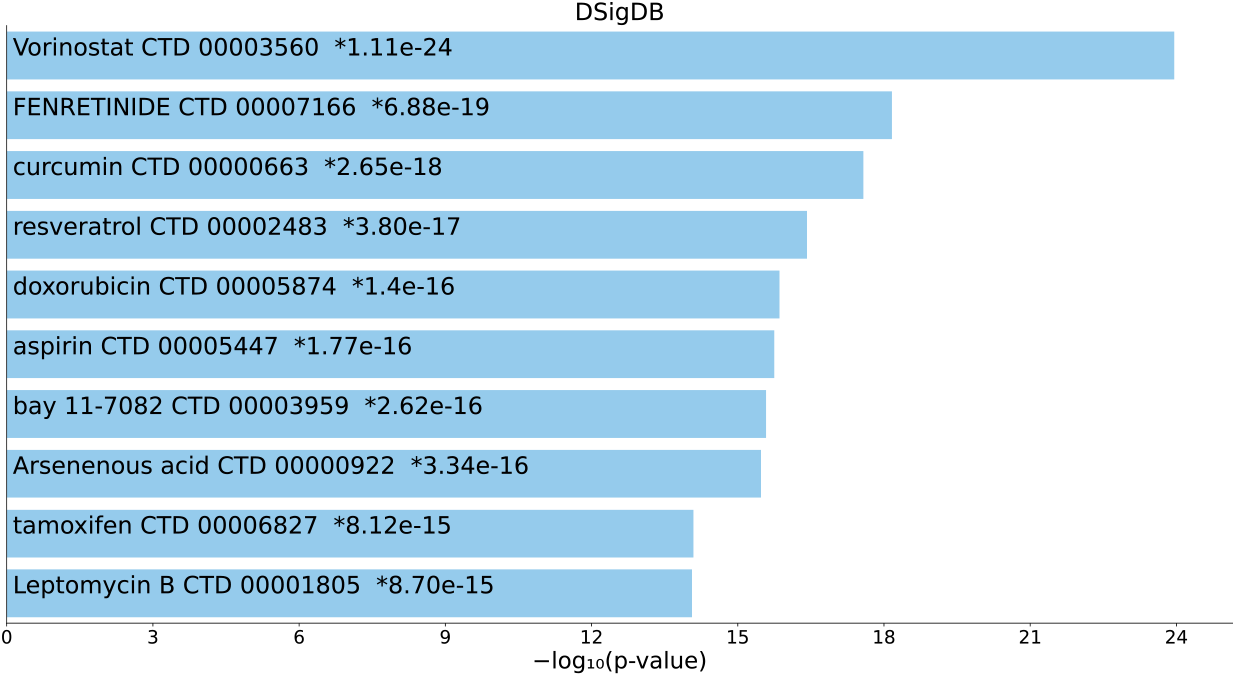
Top 10 drugs in the “DSigDB” category of Enrichr for 57 gene names selected by *u*_3*i*_ for DIP. Blue: *P <* 0.05, *: adjusted *P <* 0.05.

**Fig. 13.**
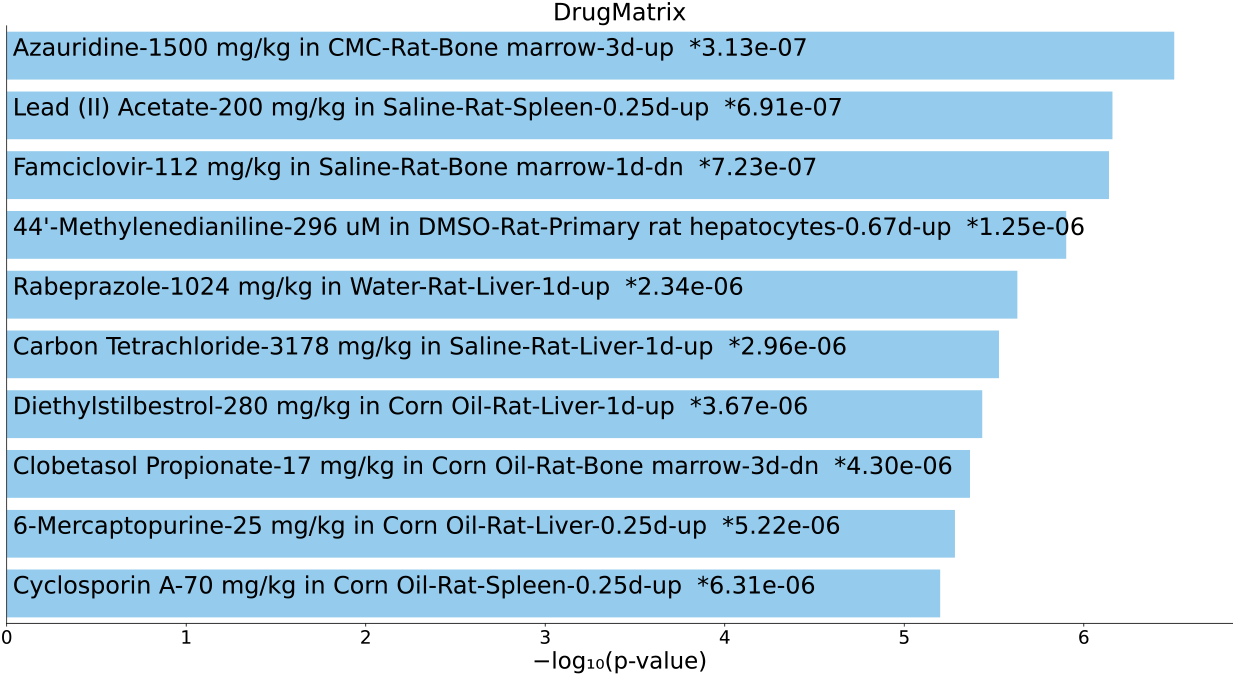
Top 10 drugs in the “DrugMatrix” category of Enrichr for 217 gene names selected by *u*_2*i*_ for BioGrid. Blue: *P <* 0.05, *: adjusted *P <* 0.05.

**Fig. 14.**
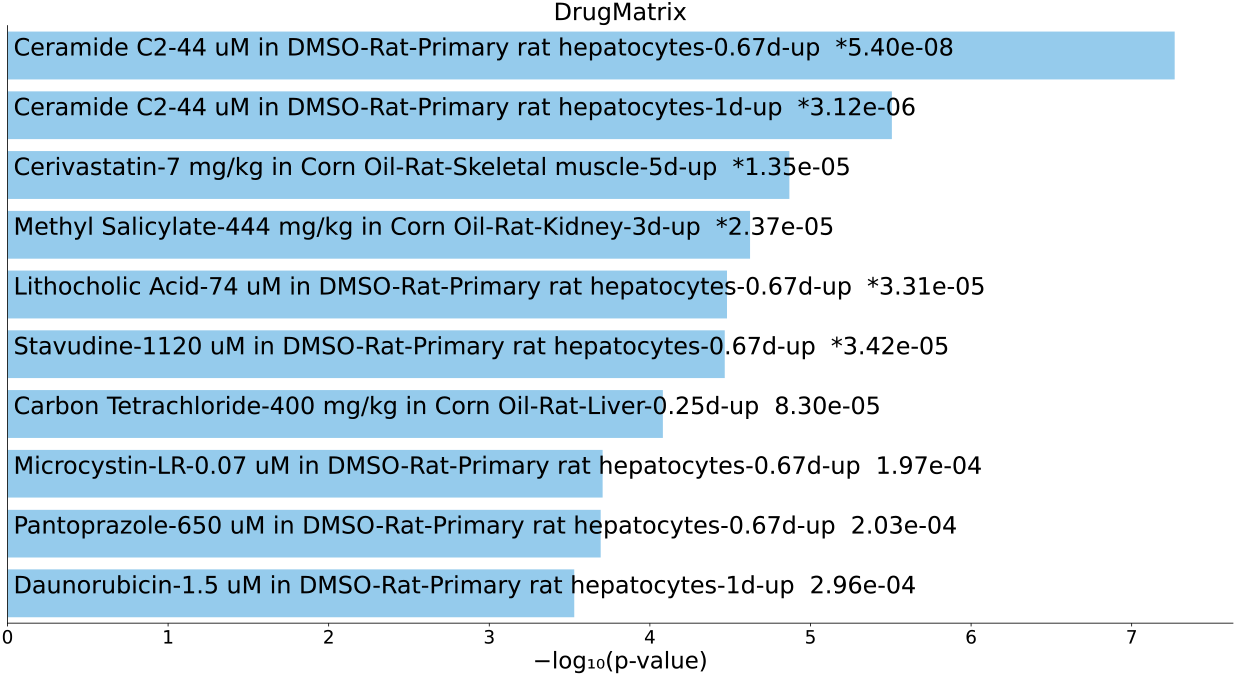
Top 10 drugs in the “DrugMatrix” category of Enrichr for 193 gene names selected by *u*_2*i*_ for DIP. Blue: *P <* 0.05, *: adjusted *P <* 0.05.

**Fig. 15.**
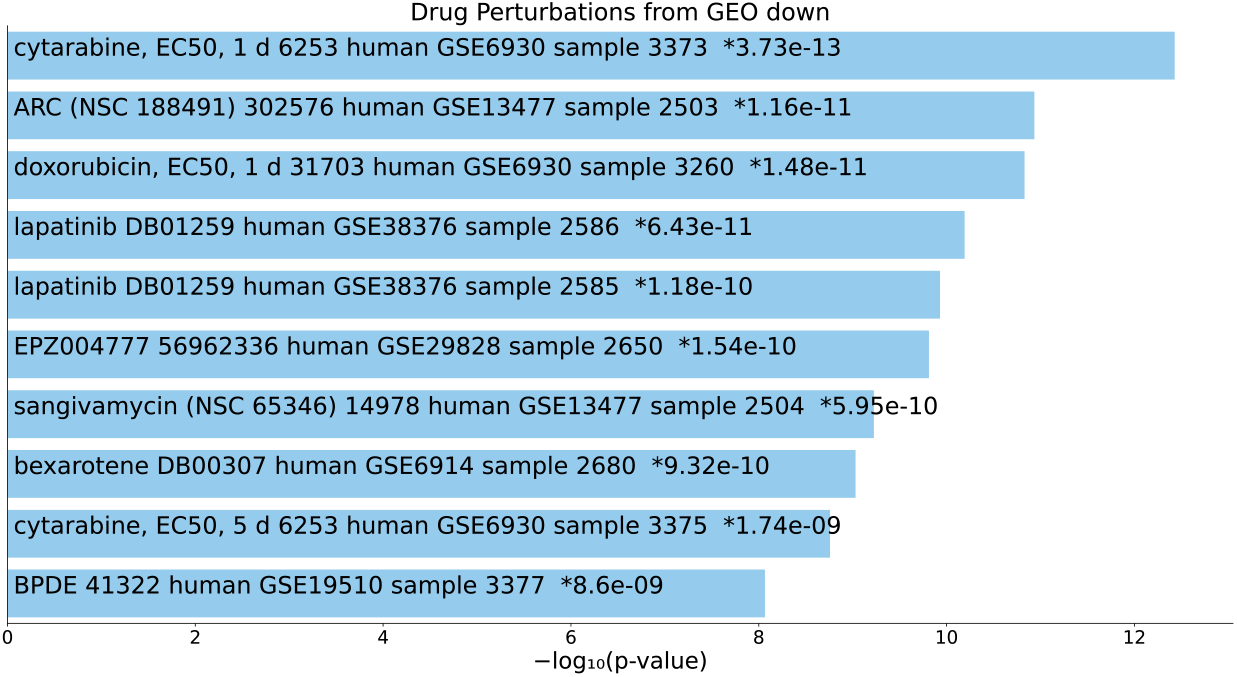
Top 10 drugs in the “Drug Perturbations from GEO down” category of Enrichr for 217 gene names selected by *u*_2*i*_ for BioGrid. Blue: *P <* 0.05, *: adjusted *P <* 0.05.

**Fig. 16.**
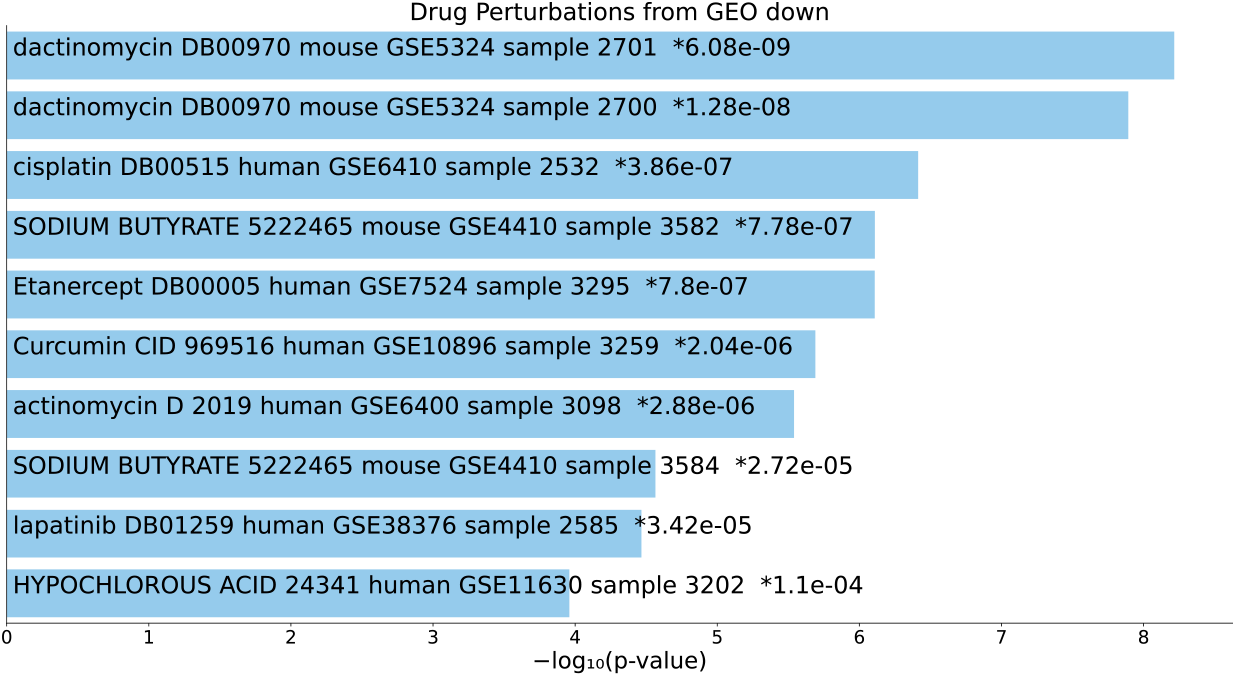
Top 10 drugs in the “Drug Perturbations from GEO down” category of Enrichr for 193 gene names selected by *u*_2*i*_ for DIP. Blue: *P <* 0.05, *: adjusted *P <* 0.05.

**Fig. 17.**
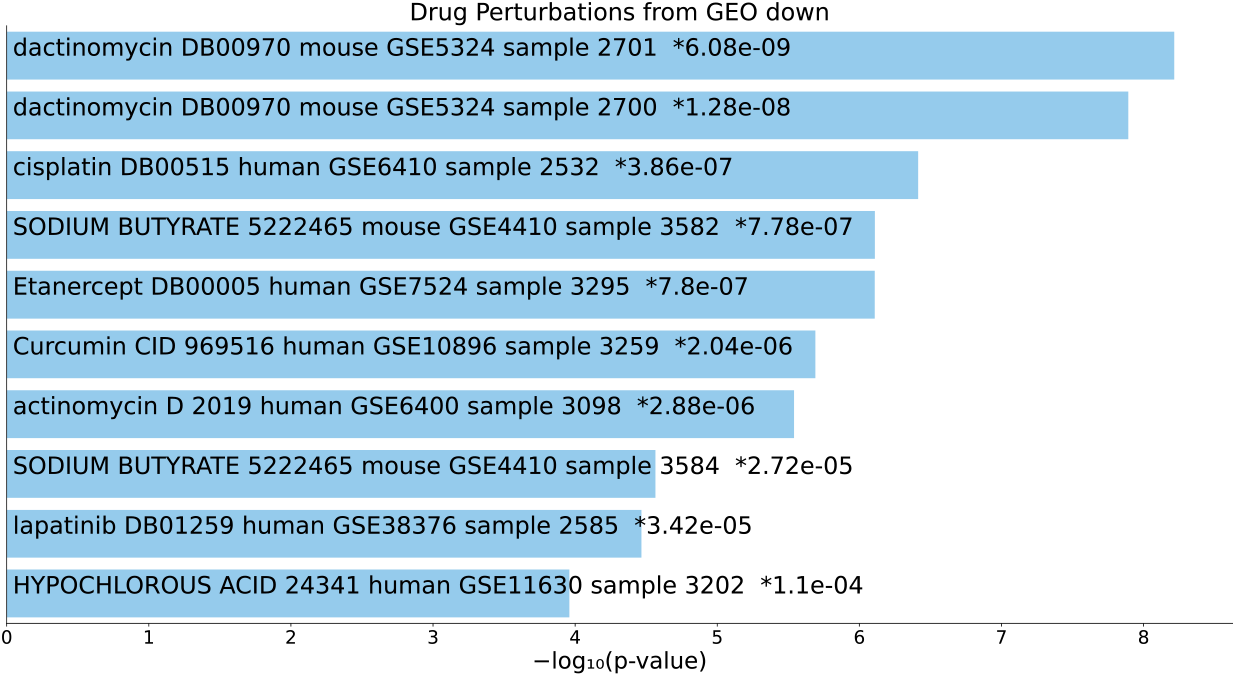
Top 10 drugs in the “Drug Perturbations from GEO down” category of Enrichr for 57 gene names selected by *u*_3*i*_ for DIP. Blue: *P <* 0.05, *: adjusted *P <* 0.05.

**Fig. 18.**
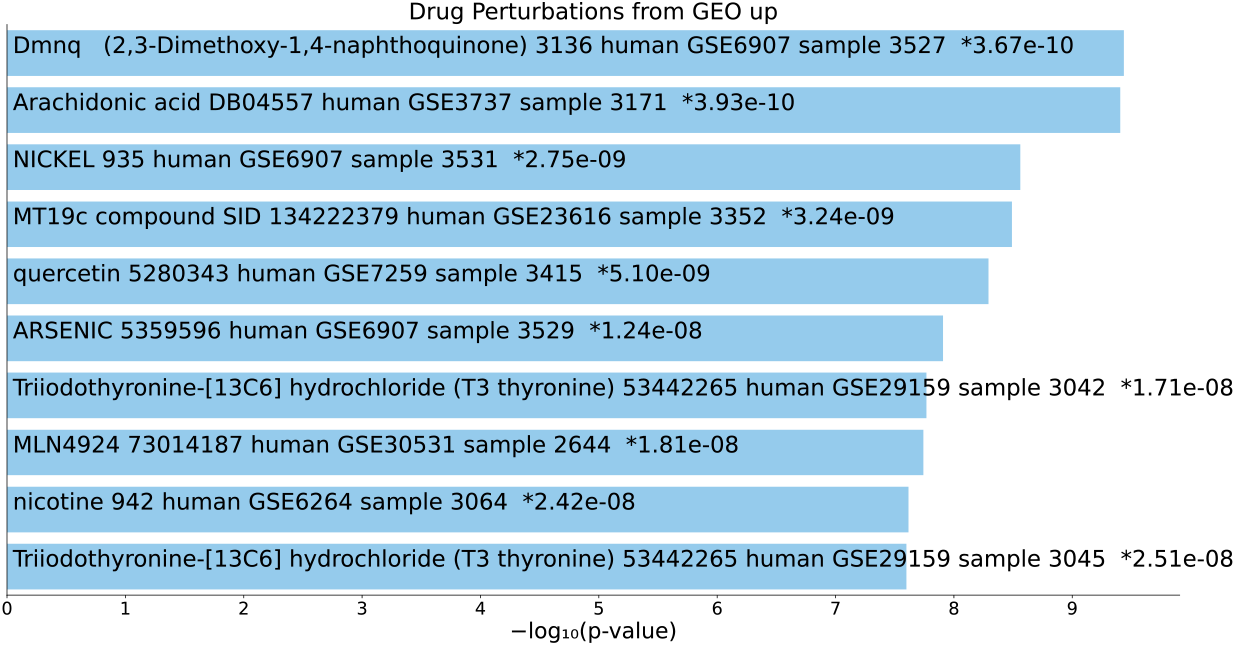
Top 10 drugs in the “Drug Perturbations from GEO up” category of Enrichr for 217 gene names selected by *u*_2*i*_ for BioGrid. Blue: *P <* 0.05, *: adjusted *P <* 0.05.

**Fig. 19.**
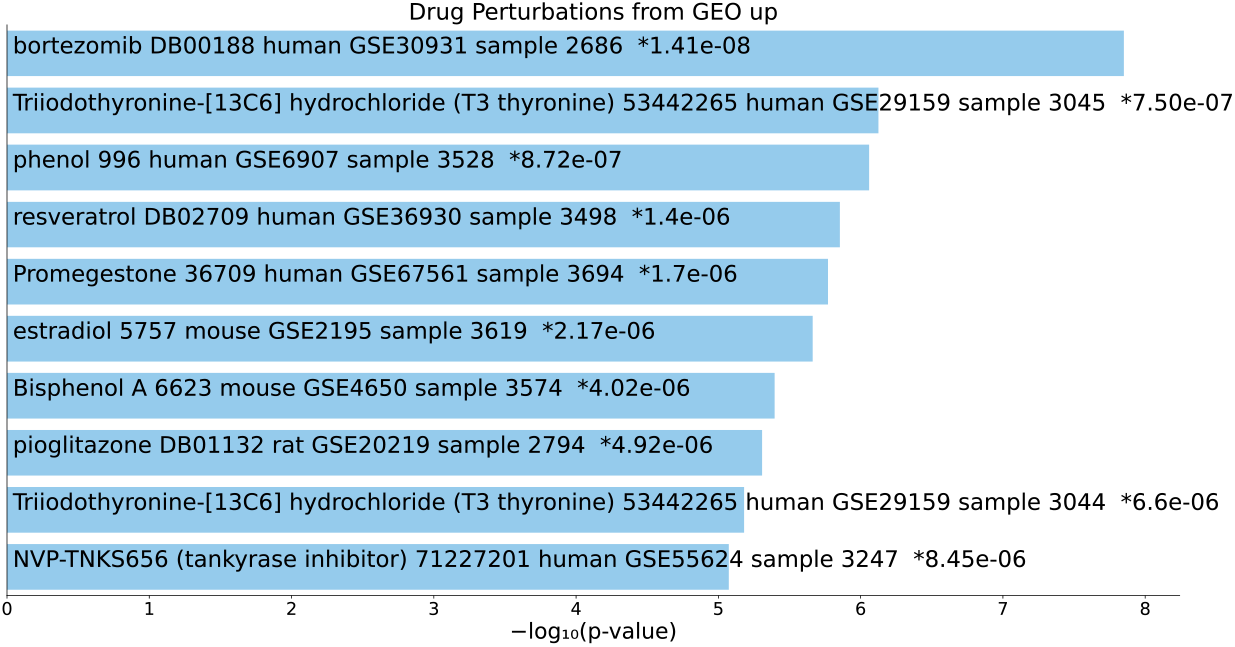
Top 10 drugs in the “Drug Perturbations from GEO up” category of Enrichr for 193 gene names selected by *u*_2*i*_ for DIP. Blue: *P <* 0.05, *: adjusted *P <* 0.05.

**Fig. 20.**
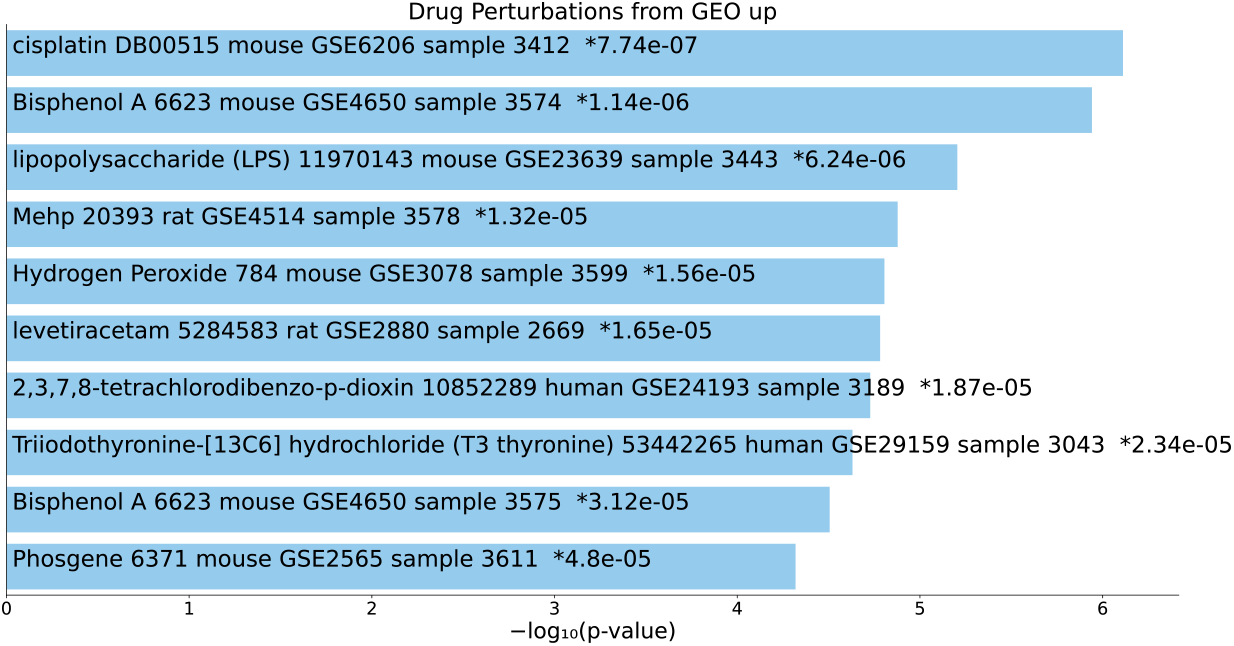
Top 10 drugs in the “Drug Perturbations from GEO up” category of Enrichr for 57 gene names selected by *u*_3*i*_ for DIP. Blue: *P <* 0.05, *: adjusted *P <* 0.05.

As can be seen above, although it is not true for all categories, three sets of gene names were often associated with lists of drugs that significantly target these sets of gene names. Thus, we can conclude that our proposed gene-centric drug repositioning strategy can be performed employing TD-based unsupervised FE to select a set of proteins based on PPI networks, regardless of the PPI dataset employed.

### 2.2 PCA-based unsupervised FE

One might wonder why we needed to integrate human and mouse PPI. Simply using only human PPI might result in a similar or even better performance. To address this problem, we applied PCA-based unsupervised FE [10, 11] to human PPI and attempted to obtain a set of proteins associated with adjusted *P* -values less than 0.01. Nevertheless, when considering the *u*_2*i*_ for BioGRID, although 158 proteins were associated with adjusted *P* -values less than 0.01, the “DrugMatrix” category failed to identify enriched drug for gene names associated with these proteins. Similarly, when considering *u*_2*i*_ for DIP, only one protein was found to be associated with adjusted *P* -values of less than 0.01. This results suggest that TD-based unsupervised FE is superior to PCA-based unsupervised FE and thereby worth testing.

### 2.3 Cluster analyses

However, another concern is whether TD-based unsupervised FE is required when selecting a set of proteins since several methods exist with which to select sub-clusters within large networks. Such methods can replace TD-based unsupervised FE in the selection of a set of proteins. Following Zhao et al [17], we employed three methods, multi-level [18], label-progration [19], and edge-betweenness [20], which enabled us to identify sub-clusters. When considering PPI human for BioGrid, edge-betweenness failed to converge after waiting more than eight hours (i.e longer than the ten minutes required for TD-based unsupervised FE, which includes the time consuming process of generating tensor) *n*_*ii*_*′*_*k*_ ∈ ℝ^*N ×N ×*2^), and the first and the second clusters generated by multi-level were too large (6406 and 2776). The first and the second clusters generated by label-progration were 24501 and 1, which was not reasonable at all. However, when human PPI for DIP was considered, the results improved. Table 1 lists the performances for the identification of diseases and drugs significantly associated with selected proteins in individual identified sub-clusters (label-progration could not identify large enough clusters to be used for enrichment analyses). It is clear that the tensor method outperforms the clustering methods, even for DIP. In conclusion, in contrast to conventional cluster analysis in which cluster sizes varied heavily depending on the PPI dataset used, using TD-based unsupervised FE can provide a stable and reasonable size for a set of genes. Thus, TD-based unsupervised FE appears to be superior to conventional cluster analysis.

**Table 1.** Performances of various protein selection methods for DIP. Os denotes cases with more than or equal to 10 drugs/associated with adjusted *P* -values less than 0.05. Otherwise, the corresponding practical numbers are presented.

| Methods<br>Order | Tensor |  | multi-level |  | edge-betweenness |  |
| --- | --- | --- | --- | --- | --- | --- |
|  | 2nd | 3rd | 1st | 2nd | 1st | 2nd |
| # of proteins | 195 | 55 | 144 | 138 | 39 | 82 |
| # of genes | 193 | 57 | 128 | 164 | 30 | 111 |
| Jensen Diseases | $\bigcirc$ | $\bigcirc$ | $\bigcirc$ | $\bigcirc$ | 7 | 5 |
| OMIM Disease | 9 | 9 | 8 | 0 | 0 | 0 |
| LINCS L1000 Chem Pert Consensus Sigs | $\bigcirc$ | 0 | 4 | 1 | 0 | $\bigcirc$ |
| DSigDB | $\bigcirc$ | $\bigcirc$ | $\bigcirc$ | $\bigcirc$ | 0 | $\bigcirc$ |
| DrugMatrix | 6 | 0 | 0 | 0 | 0 | 0 |
| Drug Perturbations from GEO down | $\bigcirc$ | $\bigcirc$ | 0 | 0 | 0 | 2 |
| Drug Perturbations from GEO up | $\bigcirc$ | $\bigcirc$ | 6 | 0 | 1 | 0 |

### 2.4 Other species

Although we only used human and mouse PPI, it is possible to employ other combinations because BioGRID and DIP to include more species-specific PPI. To validate this, we initially examined a combination of human and rat PPI for BioGRID. However, this outcome was not very promising. Next, 271 proteins associated with adjusted *P* values less than 0.01 were selected, and the gene names associated with these proteins were uploaded. The “Jensen Diseases” and “OMIM Diseases” categories identified two and zero diseases enriched with the uploaded gene names, respectively. Because the identification of enriched diseases is the starting point, if the process fails at this point, there are no way to proceed. The reason for this is the inadequateness of rat PPI, which includes too few PPI; integrating human and rat PPI results in the wrong conclusion, namely that missing PPI in rats indicates a lack of interactions, although this may simply mean that PPI in rats has not been thoroughly investigated. Thus, combinations other than that between human and mouse PPI will not work until a more comprehensive PPI for species close to humans can be obtained.

In conclusion, PPI can be a useful source of information for drug repositioning when used in combination with TD.

## 3 Discussion

To understand the set of genes selected in this analysis, we computed the mean number of bindings of individual proteins, defined as

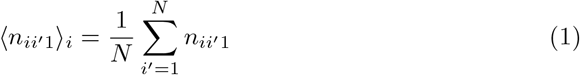

and compared ⟨*n*_*ii*_*′*_1_⟩_*i*_ between the selected and non-selected proteins for the human PPI. As can be seen in Table 2, ⟨*n*_*ii*_*′*_1_⟩_*i*_ for 217 proteins selected by *u*_2*i*_ for BioGRID, 195 proteins selected by *u*_2*i*_ for DIP, and 55 proteins selected by *u*_3*i*_ for DIP were always significantly larger than those of other (i.e. not selected) proteins. These results suggest that TD-based unsupervised FE selects proteins with more interactions in a fully data-driven and unsupervised manner.

**Table 2.** To compare ⟨*n*_*ii*_*′*_1_⟩_*i*_ between the selected and not selected proteins when using various 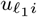 to select proteins, *P* -values were computed using the *t* test based on the alternative hypothesis that the mean ⟨*n*_*ii*_*′*_1_⟩_*i*_ of the selected proteins was larger that that of not selected proteins.

| $u_{\ell_1 i}$s used | $u_{2i}$ | | $u_{3i}$ | | $u_{6i}$ | |
| --- | --- | --- | --- | --- | --- | --- |
|  | Selected | Not selected | Selected | Not selected | Selected | Not selected |
| BioGRID |  |  |  |  |  |  |
| Mean $\langle n_{ii'/1} \rangle_i$ | $3.63 \times 10^{-2}$ | $1.84 \times 10^{-3}$ | $1.97 \times 10^{-2}$ | $1.78 \times 10^{-3}$ | — | — |
| $P$-value | $5.32 \times 10^{-37}$ | | $2.53 \times 10^{-59}$ | | — | |
| DIP |  |  |  |  |  |  |
| Mean $\langle n_{ii'/1} \rangle_i$ | $1.12 \times 10^{-3}$ | $4.17 \times 10^{-4}$ | $3.12 \times 10^{-3}$ | $4.13 \times 10^{-4}$ | $2.14 \times 10^{-3}$ | $4.26 \times 10^{-4}$ |
| $P$-value | $2.4 \times 10^{-4}$ | | $5.3 \times 10^{-5}$ | | $2.04 \times 10^{-13}$ | |

Since proteins selected via TD-based unsupervised FE have more interactions, we proceeded to verify whether proteins were being selected for use in drug repositioning simply due to their greater number of interactions (hereafter, denoted as hub proteins). To verify this, we selected the top 200 and 50 proteins with more interactions (these numbers were selected to be close to the number of proteins selected using *u*_2*i*_ and *u*_3*i*_. See Supplementary Information for a list of proteins and gene names) using human PPI. Table 3 lists the confusion matrices for the selected proteins. Although they significantly overlapped because the majority were not shared, distinct sets of proteins were identified.

**Table 3.**
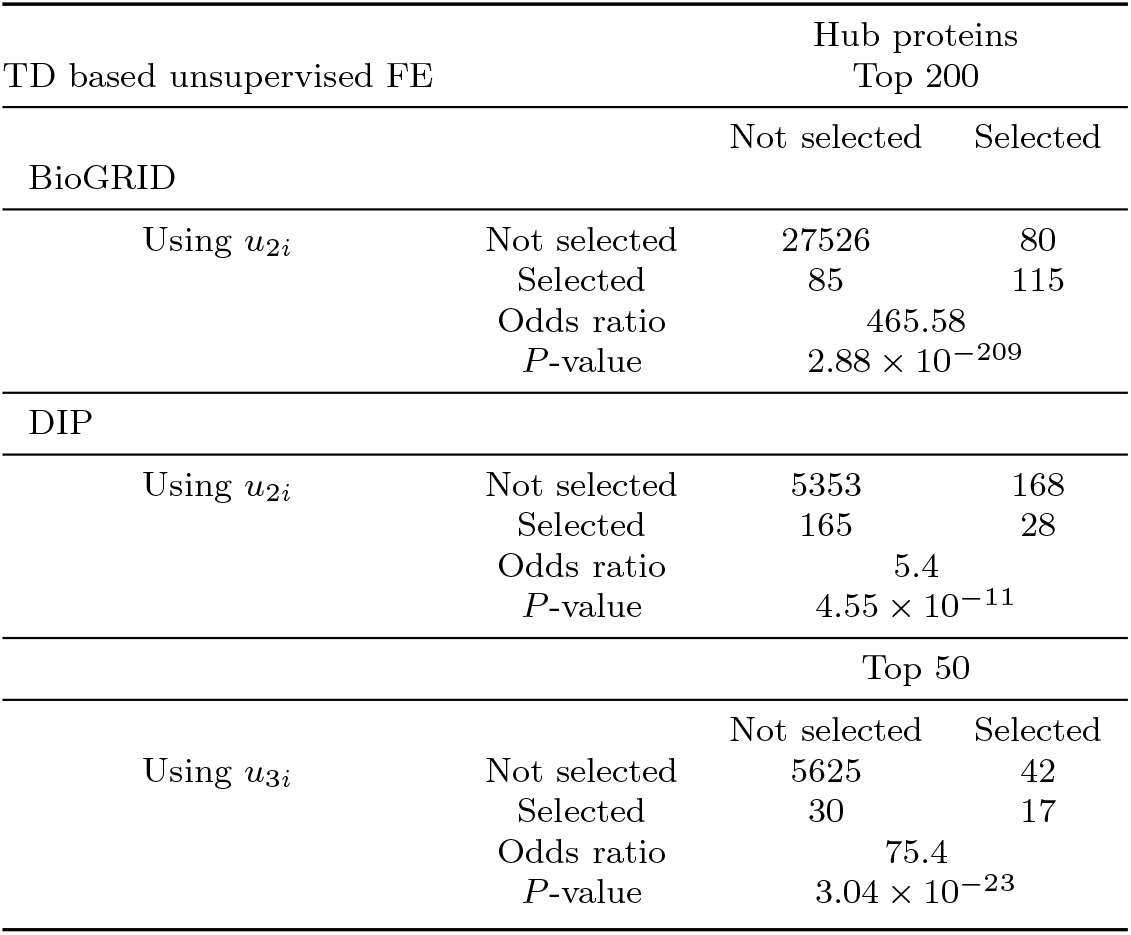
Confusion matrix of selected proteins between TD-based unsupervised FE and hub proteins.

We uploaded the associated gene names to Enrichr to evaluate the top 200 and 50 hub proteins. Table 4 lists the performance achieved by the hub proteins. (for details on the top diseases and drugs in individual categories: see Tables S20 to S26 for the top 200 hub proteins for BioGRID, Tables S29 to S33 for the top 200 hub proteins for DIP, and Tables S34 to S40 for the top 50 hub proteins for DIP). Their performances were essentially the same as those of TD-based unsupervised FE, excluding DrugMatrix for DIP.

**Table 4.**
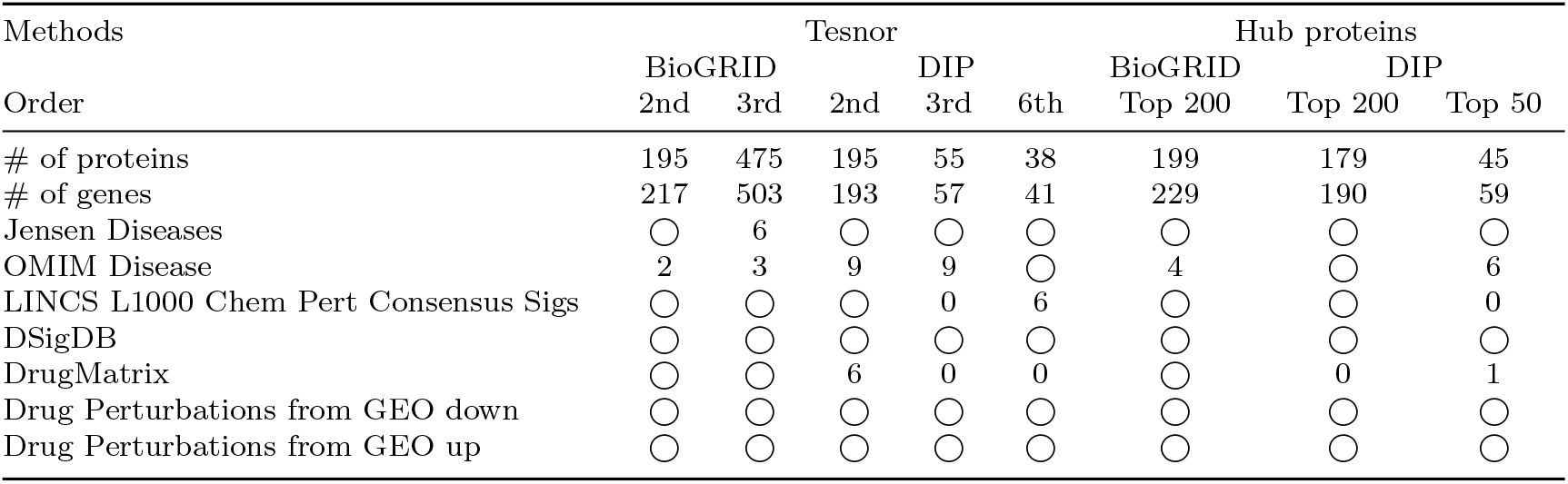
Performances achieved by hub proteins. Os denote case with more than or equal to 10 drugs/associated with adjusted *P* -values less than 0.05. Otherwise, the corresponding practical numbers are presented.

Comparing the performance of TD-based unsupervised FE with hub proteins, to the best of our knowledge, there are no other studies that use hub proteins derived from only PPI for drug repositioning without using other information. This is because hub proteins derived from PPI themselves, without any other information, cannot be used for drug repositioning for cancer. This was clear from the fact that proteins selected by TD-based unsupervised FE were hub proteins. Without TD-based unsupervised FE, we could not identify hub proteins derived only from PPI that could be used for drug repositioning for cancer. Secondly, the hub proteins for DIP failed to identify hits in DrugMatrix. DrugMatrix is an *in vivo* specific database. Thus, the hub proteins for DIP do not have the ability to identify drugs that may be useful in *in vivo* experiments. Thus, even if hub proteins can achieve a performance similar to that of the *in vitro* experiments, TD-based unsupervised FE is still useful, at least for DIP. Third, using hub proteins, we cannot target diseases other than cancers because hub proteins are enriched mainly in cancers. For the top 200 hub proteins for BioGRID, seven cancers were identified (“stomach cancer,” “adenoma,” “cancer,” “immune system cancer,” “esophageal carcinoma,” “intestinal benign neoplasm,” and “lymphoid leukemia”) within the top ranked diseases in the “Jensen Diseases” category and as many as five cancers (“ovarian cancer,” “breast cancer,” “colorectal cancer,” “thyroid carcinoma,” and “prostate cancer”) within the top ranked diseases in the “OMIM Diseases” category (Table S20 and S21). For the top 200 hub proteins for DIP, as many as six cancers were identified (“cancer,” “lymphoid leukemia,” “intestinal benign neoplasm,” “immune system cancer,” “biliary tract cancer,” and “stomach cancer”) within the top ranked diseases in the “Jensen Diseases” category and as many as seven (“colorectal cancer,” “ovarian cancer,” “breast cancer,” “leukemia,” “lung cancer,” “melanoma,” and “pancreatic cancer”) within the top ranked diseases in the “OMIM Diseases” category (Table S27 and S28). For the top 50 hub proteins for DIP, as many as five cancers (“cancer,” “DOID:9917” (pleural cancer), “thyroid cancer”, “esophageal carcinoma”, and “intestinal benign neoplasms”) were identified within the top ranked diseases in the “Jensen Diseases” category and as many as five (“colorectal cancer,” “ovarian cancer,” “breast cancer,” “lung cancer,” and “pancreatic cancer”) within the top ranked diseases in the “OMIM Diseases” category (Table S34 and S35). Nevertheless, because TD-based unsupervised FE has the freedom to select 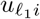, it can provide a set of proteins enriched in diseases other than cancer. Tables S41 and S42, and Figs. 21 and 22 list the diseases enriched in proteins selected by *u*_3*i*_ for BioGRID, while Tables S43 and S44, and Figs. 23 and 24 list the diseases enriched in proteins selected by *u*_6*i*_ for DIP (see Supplementary Information for the list of proteins and gene names); they are mainly distinct from cancers (only one cancer (“stomach cancer”) for BioGRID and only one cancer (“cancer”) for DIP within the top 10 diseases in the “Jensen Diseases” category (Tables S41 and S43 and Figs. 21 and **??**) and Three cancers (“thyroid carcinoma,” “gastric cancer,” and “ovarian cancer “) for BioGRID and three cancers (“pancreatic cancer,” “melanoma,” and “lung cancer”) for DIP were identified within the top 10 diseases in the “OMIM Diseases” category (Tables S42 and S44 and Figs. 22 and 24). For these cancers, sufficient drugs exist (for detailed drug names, see Table S22–S31). Thus, the results of TD-based unsupervised FE are useful even if hub proteins can be used to identify drugs for cancers.

**Fig. 21.**
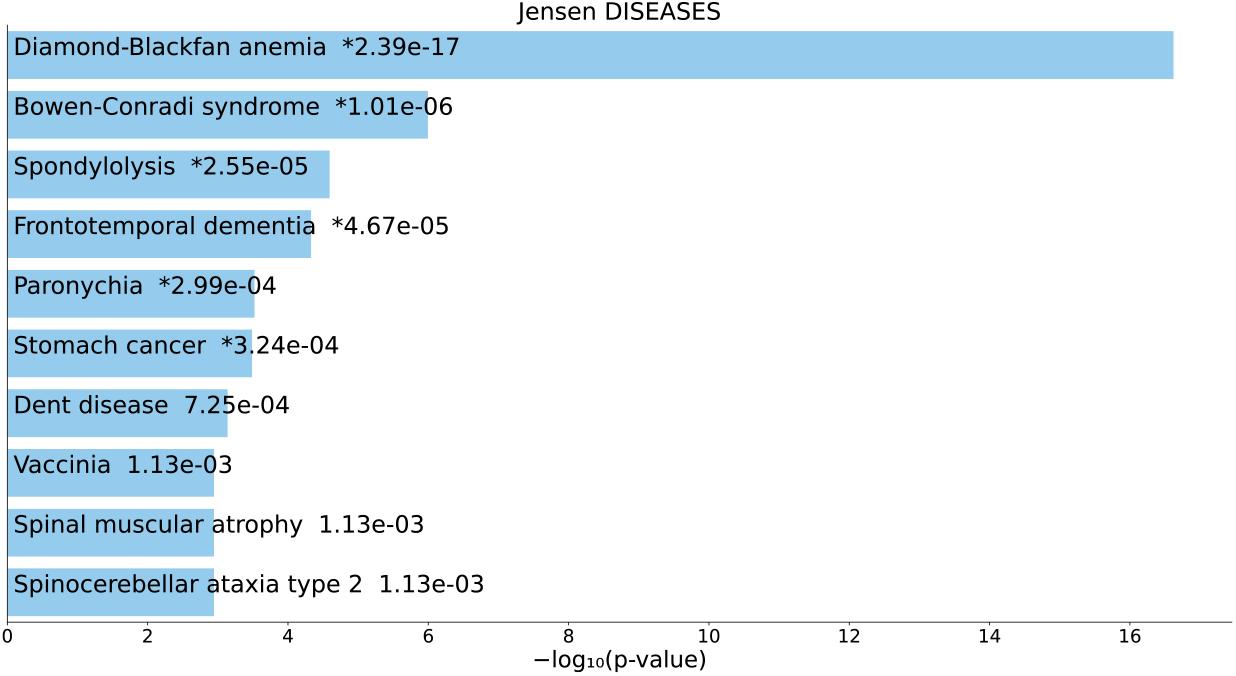
Top 10 diseases in the “Jensen Diseases” category of Enrichr for 502 gene names selected by *u*_3*i*_ for BioGrid. Blue: *P <* 0.05, *: adjusted *P <* 0.05.

**Fig. 22.**
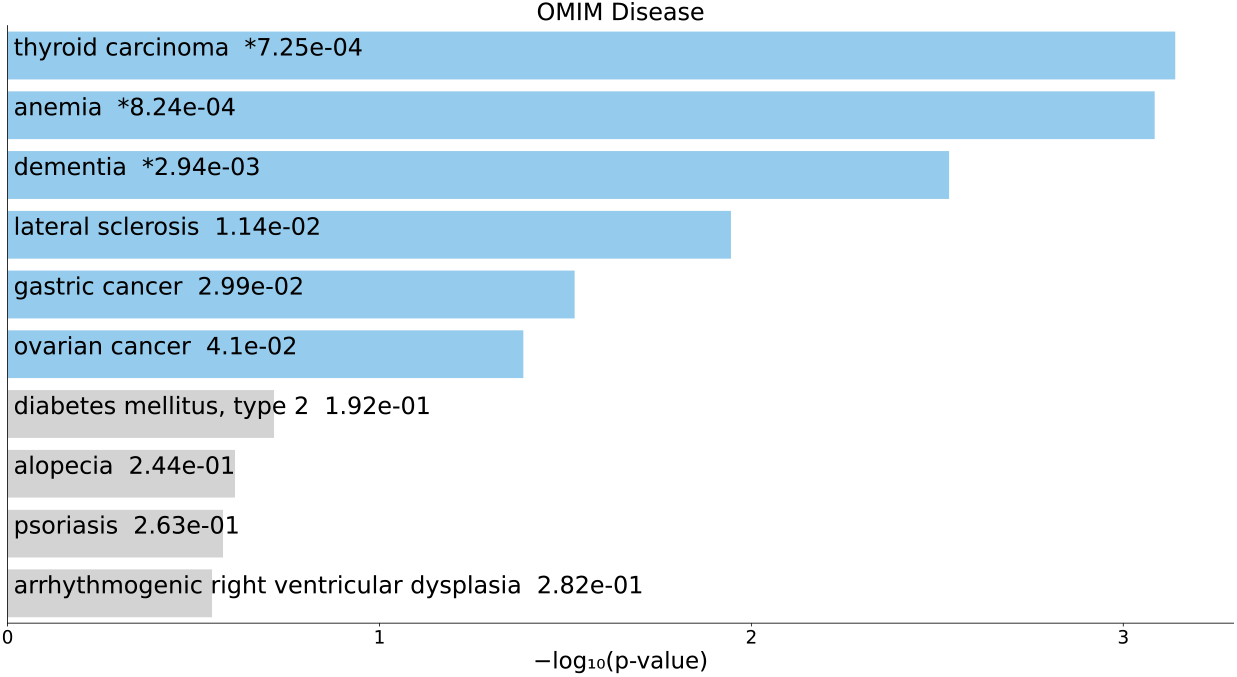
Top 10 diseases in the “OMIM Diseases” category of Enrichr for 502 gene names selected by *u*_3*i*_ for BioGrid. Blue: *P <* 0.05, *: adjusted *P <* 0.05.

**Fig. 23.**
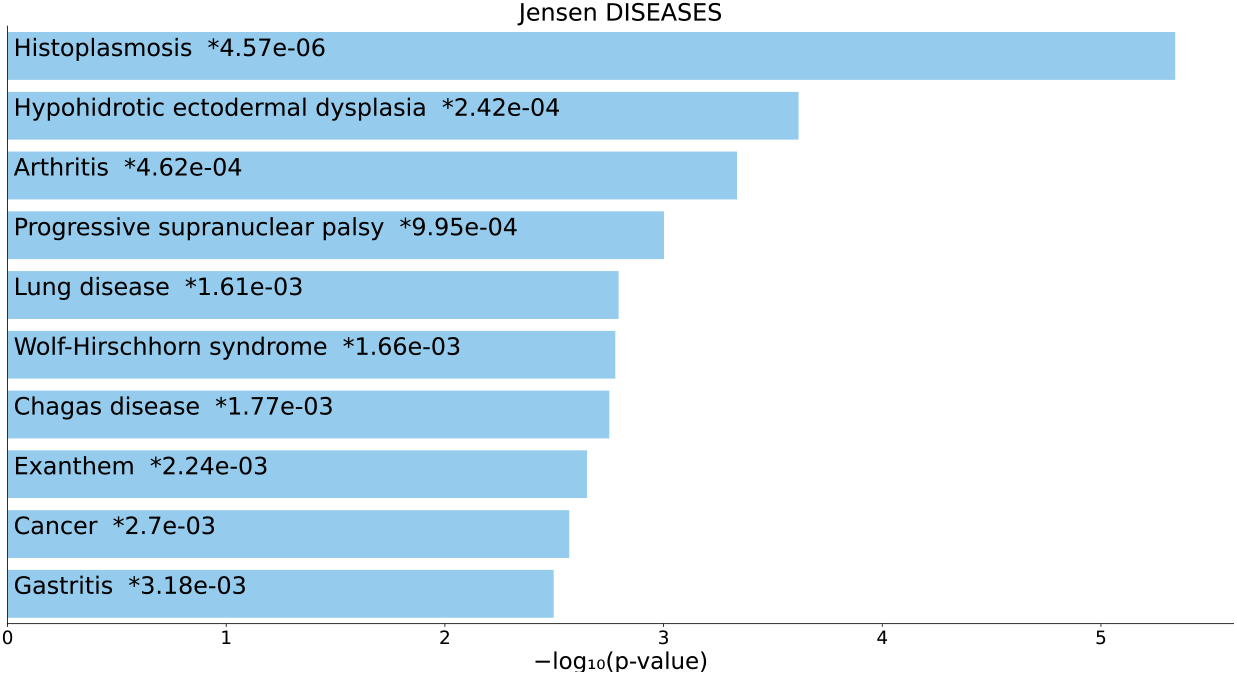
Top 10 diseases in the “Jensen Diseases” category of Enrichr for 41 gene names selected by *u*_6*i*_ for DIP. Blue: *P <* 0.05, *: adjusted *P <* 0.05.

**Fig. 24.**
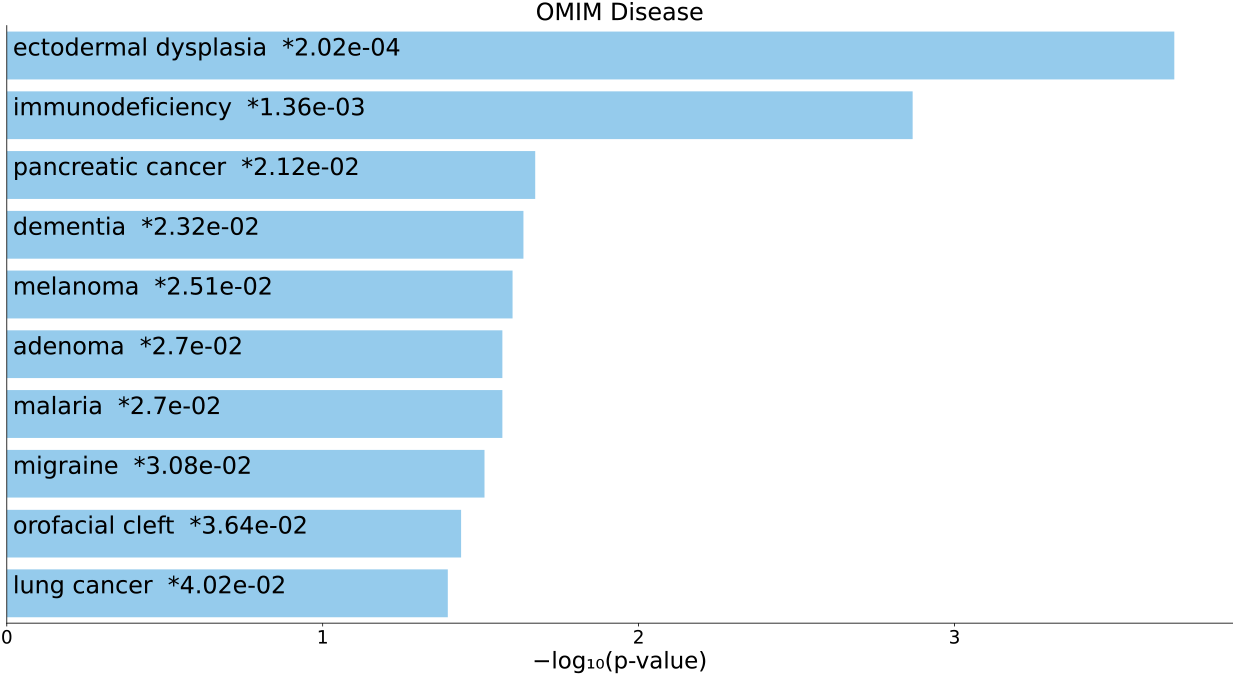
Top 10 diseases in the “OMIM Diseases” category of Enrichr for 41 gene names selected by *u*_6*i*_ for DIP. Blue: *P <* 0.05, *: adjusted *P <* 0.05.

It is worth considering the differences between hub proteins and the proteins selected by TD-based unsupervised FE if they are not identical. To determine this, the mean correlation coefficients between proteins where *r*(*n*_*ii*_*′*_1_, *n*_*ii*”1_) is Pearson’s correlation coefficient between {*n*_*ii*_*′*_1_|1 ≤ *i* ≤ *N* } and {*n*_*ii*_*′*_1_|1 ≤ *i* ≤ *N* }, were computed with human PPI. Table 5 lists the values of ⟨*r*⟩. Because proteins selected by TD-based unsupervised FE for DIP had a substantially larger ⟨*r*⟩, the distinctions between the proteins selected by TD-based unsupervised FE and the hub proteins were correlated. The fact that a larger ⟨*r*⟩ means that proteins share the proteins with which they interact suggests that proteins that share interactions may be the key to identifying effective drugs in *in vivo* experiments. In reality, the top 200 hub proteins for BioGRID have a larger ⟨*r*⟩ and many enrichments in the “DrugMatrix” category (Table S12 and Fig. 13). Thus, we can guarantee a high correlation for hub proteins without consulting TD-based unsupervised FE, which allows us to identify genes that can be used to identify effective drugs, even in *in vivo* experiments.

**Table 5.** ⟨*r*⟩s and *P* -values computed using a one way *t* test

| | $\langle r \rangle$ | $P$-values |
| --- | --- | --- |
| BioGRID |  |  |
| Proteins selected using $u_{2i}$ | $1.97 \times 10^{-1}$ | 0 |
| Proteins selected using $u_{3i}$ | $1.60 \times 10^{-1}$ | 0 |
| Top 200 proteins | $2.02 \times 10^{-1}$ | 0 |
| DIP |  |  |
| Proteins selected using $u_{2i}$ | $3.29 \times 10^{-1}$ | 0 |
| Proteins selected using $u_{3i}$ | $9.64 \times 10^{-2}$ | $7.70 \times 10^{-211}$ |
| Proteins selected using $u_{6i}$ | $8.14 \times 10^{-2}$ | $5.84 \times 10^{-70}$ |
| Top 200 proteins | $7.49 \times 10^{-3}$ | $3.97 \times 10^{-128}$ |
| Top 50 proteins | $1.19 \times 10^{-3}$ | $6.84 \times 10^{-18}$ |

Because we employed a gene-centric strategy, the drugs and diseases identified were associated with common genes. Nevertheless, one might wonder whether this always guarantees the effectiveness of the identified drugs against the diseases. The so-called docking simulation is not a good idea for validation because Enrichr is used to relate genes to drugs or diseases in order to identify relationships based on gene expression. Thus, this does not always mean that the selected genes are direct targets of drugs, but simply those whose expression is altered by drug treatment, since the genes are in the downstream pathway. However, to the best of our knowledge, search engines specifically for studies on drug-disease pairs within the proposed list of drugs and diseases do not exist. Alternatively, to achieve this, we employed a large language model (LLM). We sought to identify pairs of drugs and diseases for which research has been reported. To avoid the wrong relationships being reported by LLM due to hallucinations, we verified whether other papers existed in support of the relationship reported our LLM.

Table 6 lists the reported and validated combination of drugs and diseases between the “Jensen Diseases” category by *u*_3*i*_ for BioGRID (Table S41 and Fig. 21) and the “DSigDB” category or “DrugMatrix” category by *u*_3*i*_ for BioGRID (Table S23 or S24). The reason we employ this is simply because it is difficult to validate the correspondence between diseases and drugs if diseases are mostly composed of cancers because many cancers share effective drugs. Therefore, it is preferable to use a set of diseases other than cancer. It is evident that the combinations of drugs and diseases reported by TD-based unsupervised FE are associated with previously published studies, despite the diverse types of diseases. Thus, we concluded that the pairs of diseases and drugs identified in this study are likely to include promising pairs.

**Table 6.**
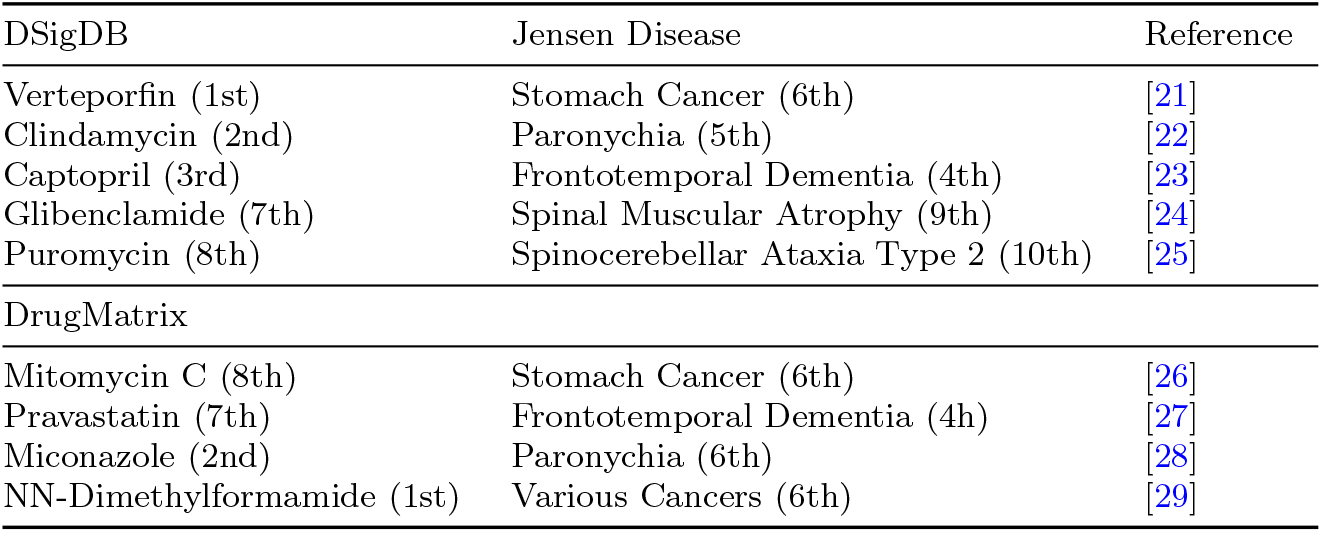
Diseases and drugs whose relation is reported by LLM. Their corresponding ranks in categories appear in parentheses.

The limitation of our present methods is that we cannot always get enough number of PPI information for most species. As can be seen above, when we considered rat, we could not get any good results because of the lack of enough PPI information. We expect that more advanced methods for PPI identification will be developed.

## 4 Methods

### 4.1 Protein-protein interactions (PPI)

#### 4.1.1 BioGRID

The PPI data were downloaded from BioGRID [30]. PPI classified by species, BIOGRID-ORGANISM-4.4.236.tab3.zip, were downloaded. Three species-specific files, BIOGRID-ORGANISM-Homo sapiens-4.4.236.tab3.txt, BIOGRID-ORGANISM-Mus musculus-4.4.236.tab3.txt, and BIOGRID-ORGANISM-Rattus norvegicus-4.4.236.tab3.txt were extracted for analysis.

#### 4.1.2.DIP

The PPI data were also downloaded from DIP [31]. The datasets used were species-specific sets for *Homo Sapiens*, Hsapi20170205, and *Mus musculus*, Mmusc20170205. “full [.gz]” were downloaded as tab-limited files.

#### 4.1.3 Tensor format

The PPI files were further loaded into R using the read_csv command as 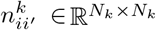 (*k* = 1: human, *k* = 2: mouse, and *k* = 3: rat (only for BioGRID)), which takes 1 when the *i*th and *i*^*′*^th proteins interact with each other; otherwise, 0. As these matrices were sparse, they were stored in a sparse matrix format using the Matrix [32] package.

### 4.2 Principal component analysis (PCA)-based unsupervised feature extraction (FE)

Singular value decomposition was applied to the irlba function in the irlba package [33] for human PPI 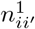. As a result, we obtained

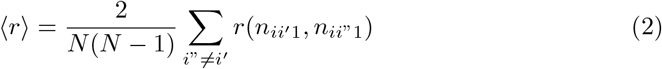

where *λ*_*𝓁*_ is a singular value, and *u*_*𝓁i*_ ∈ ℝ^*N ×N*^ is the singular value vector and orthogonal vector.

*P* -values are attributed to *i*th protein as

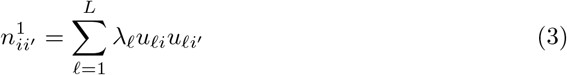

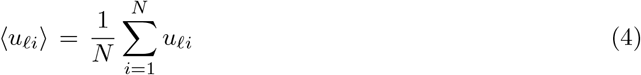

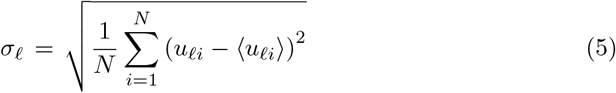

Where 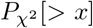 is the cumulative *χ* distribution, the argument is larger than *x* and ⟨*u*_*𝓁i*_⟩, and *σ*_*𝓁*_ are the mean and standard deviation, respectively. Thus, we assumed the following null hypothesis: *u*_*𝓁i*_ follows a Gaussian distribution. *P* -values were corrected using the BH criterion, and proteins associated with adjusted *P* -values less than 0.01 were selected.

### 4.3 Tensor decomposition (TD)-based unsupervised FE

To apply TD-based unsupervised FE to PPI, an integrated tensor that stores multiple PPIs must be constructed. Suppose we have two PPI matrices, 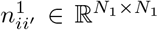 and

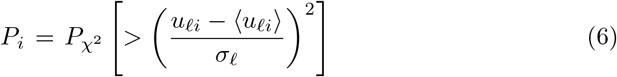

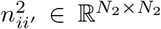. It is also assumed that there are *N*common common proteins and *N*ortho orthogonal proteins between the two datasets. Then, we merge these datasets into one tensor, *n*_*ii*_*′*_2_ ∈ ℝ^*N ×N ×*2^ where *N* = *N*_1_ + *N*_2_ – *N*common – *N*otrho as shown in (Fig. 25). 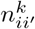, (1 ≤ *i, i*^*′*^ ≤ *N*common + *N*ortho) are placed in *n*_*ii*_*′*_*k*_ as is (yellow and red regions in Fig. 25). 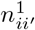, (*N*common + *N < i, i*^*′*^ ≤ *N*ortho) are placed in *n*_*ii*_*′*_1_ (the blue region in Fig. 25). 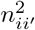, (*N*common + *N < i, i*^*′*^ ≤ *N*_2_) are placed in *n*_*ii*_*′*_2_ in a fragmented manner (green region in Fig. 25). As a result, the blue region of *n*_*ii*_*′*_2_ and green region of *n*_*ii*_*′*_1_ are blank. The shaded region is blank in *n*_*ii*_*′*_*k*_.

**Fig. 25.**
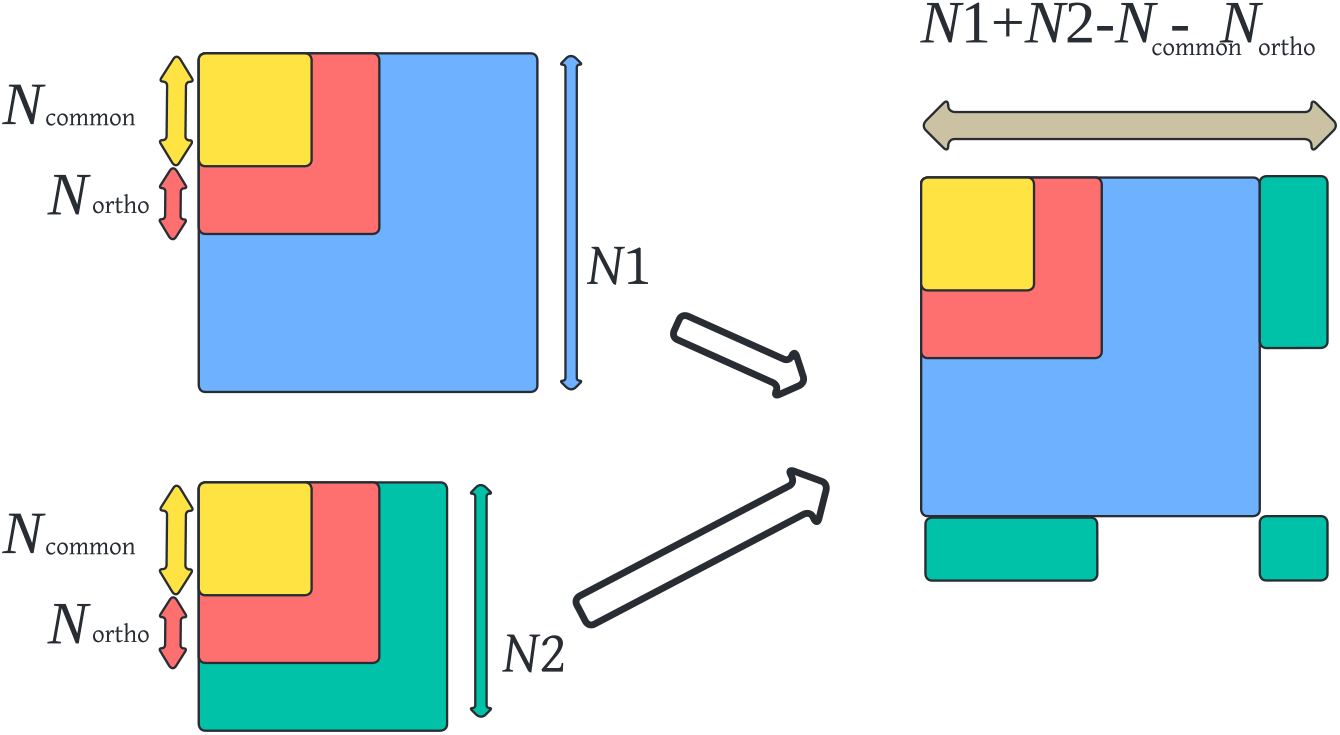
How to merge two PPI matrices into one tensor.

HOSVD was applied to *n*_*ii*_*′*_*k*_, resulting in

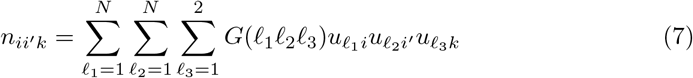

where *G* is a core tensor representing the weight (contribution) of 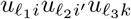 to 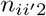 and 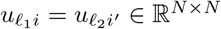 and 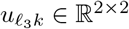 are singular value and orthogonal matrices, respectively. The attribution of *P*_*i*_s to *i*s and the selection of proteins were performed by replacing *u*_*𝓁i*_ where 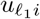 in eq. (6).

HOSVD was performed using the irlba function by applying SVD to unfolded matrices because the usual R function that can perform HOSVD does not accept a sparse matrix as input.

### 4.4 Identification of orthogonal proteins

Orthologs between human and mouse were retrieved by querying “((organism id:10090) OR ((organism id:9606) AND (reviewed:true))) AND (database:orthodb)” in uniport search (https://www.uniprot.org/). Human Uniprot accessions were converted to gene name as “XXX HUMAN” using the above retrieved information. Then, after seeking “XXX MOUSE” in the above retrieved information (using the first hit if multiple hits are found), “XXX MOUSE” was converted into the corresponding mouse Uniprot accession numbers in the information retrieved above. This results in the corresponding table of Uniprot ortholog (the above retrieved information is provided as Supplementary Material).

### 4.5 Gene ID conversion

The obtained Uniprot accession numbers attributed to proteins were converted to gene names using Uniprot ID mapping (https://www.uniprot.org/id-mapping).

### 4.6 Relating genes to drugs and diseases through enrichment analysis

Enrichr enables the validation of overlaps between two sets of genes using statistical tests that measure the probability that the observed overlap occurs by chance (the so-called *P* -value). If *P* -values are sufficiently small, the two sets of genes are significantly related. In Enrichr, one set of genes was provided by the researchers, and its overlap with the prepared sets of genes was evaluated. If the uploaded set of genes significantly overlaps with one of the prepared sets of genes, the uploaded set of genes can be said to be associated with the properties of the overlapping set of genes. In this analysis, we considered two sets of genes, diseases and drugs. For the disease gene sets, the uploaded set of genes was evaluated if it overlapped with a set of genes known to be related to diseases. The “Jensen Diseases” and “OMIM disease” are independent categories that collect the genes related to diseases. Thus, if the uploaded gene sets have significant overlap with sets of genes in either “Jensen Diseases” or “OMIM disease,” we can regard the uploaded gene sets as being related to diseases. For drug gene sets, the uploaded set of genes was evaluated if it overlapped with a set of genes whose expression was altered by drug treatment. If the uploaded set of genes has a significant overlap with the gene sets whose expression is known to be altered by some drugs, we can consider that the expression of the uploaded gene set is also altered by drug treatment. Thus, if the uploaded gene set significantly overlaps with the drug and disease gene sets simultaneously, we can expect that drug treatment can significantly alter the expression of genes whose expression is known to be altered by drug treatment. Consequently, drugs related to diseases through genes are potential drug compounds, although they are not always effective against specific diseases.

### 4.7 Query of drug-disease relation by LLM

The actual queries were performed using Microsoft Copilot. The basic structure of the prompt is provided in Supplementary Information.

The actual query prompts and LLMs replies are accessible through the following two URLs: https://copilot.microsoft.com/sl/dCHesf9R01c and https://copilot.microsoft.com/sl/jCQxrKJzZoO.

## 5 Conclusions

In this study, we demonstrated the usefulness of drug repositioning TD-based unsupervised FE applied to PPI. Although it is unlikely that only the information retrieved from PPI will be useful for disease-specific drug repositioning, our findings indicate that it works in a practical sense. Thus, TD-based unsupervised FE applied to PPI is likely to be useful for drug repositioning. Further studies will be needed to understand the extent to which this strategy is effective.

## Author Contributions

Y.-H.T. planned the study and performed the analyses. Y.-H.T. and T.T. evaluated the results, discussions, and outcomes and wrote and reviewed the manuscript. All the authors have read and agreed to the published version of this manuscript.

## Funding

This work was supported by KAKENHI [grant number 24K15168] and Chuo Universiy Tokutei Kadai kenkyuu.

## Data and Availability

All the data used in this study can be downloaded from BioGRID [30] and DIP [31]. Sample code for data processing is in https://github.com/tagtag/TDbasedUFEPPI.

## Declarations

Not applicable.

### Ethics approval and consent to participate

Not applicable.

### Consent for publication

Not applicable.

### Competing interests

The authors declare no conflict of interest.

## Notes

### Competing Interest Statement

The authors have declared no competing interest.

### Summary of Updates

Revision due to reviewer's comment. Most of tables were replaced with graphs. Tow illustrations were re-drawn by new tool. Github site was added.

## References

[1] S, Z., V, E.A., B, B.U., A, d.S.: Prediction of disease–gene–drug relationships following a differential network analysis. Cell Death & Disease 7(1), 2040–2040 (2016) 10.1038/cddis.2015.393

[2] Wong, M., Previde, P., Cole, J., Thomas, B., Laxmeshwar, N., Mallory, E., Lever, J., Petkovic, D., Altman, R.B., Kulkarni, A.: Search and visualization of gene-drug-disease interactions for pharmacogenomics and precision medicine research using genedive. Journal of Biomedical Informatics 117, 103732 (2021) 10.1016/j.jbi.2021.103732

[3] Yu, H., Choo, S., Park, J., Jung, J., Kang, Y., Lee, D.: Prediction of drugs having opposite effects on disease genes in a directed network. BMC Systems Biology 10(Suppl 1), 2 (2016) 10.1186/s12918-015-0243-2

[4] Sun, P.G.: The human drug–disease–gene network. Information Sciences 306, 70–80 (2015) 10.1016/j.ins.2015.01.036

[5] Qahwaji, R., Ashankyty, I., Sannan, N.S., Hazzazi, M.S., Basabrain, A.A., Mobashir, M.: Pharmacogenomics: A genetic approach to drug development and therapy. Pharmaceuticals 17(7) (2024) 10.3390/ph17070940

[6] Iida, M., Iwata, M., Yamanishi, Y.: Network-based characterization of disease–disease relationships in terms of drugs and therapeutic targets. Bioinformatics 36(Supplement 1), 516–524 (2020) 10.1093/bioinformatics/btaa439 https://academic.oup.com/bioinformatics/article-pdf/36/Supplement 1/i516/56702295/bioinformatics 36 supplement1 i516.pdf

[7] Quan, Y., Luo, Z.-H., Yang, Q.-Y., Li, J., Zhu, Q., Liu, Y.-M., Lv, B.-M., Cui, Z.-J., Qin, X., Xu, Y.-H., Zhu, L.-D., Zhang, H.-Y.: Systems chemical genetics-based drug discovery: Prioritizing agents targeting multiple/reliable disease-associated genes as drug candidates. Frontiers in Genetics 10 (2019) 10.3389/fgene.2019.00474

[8] Wang, L., Wang, Y., Hu, Q., Li, S.: Systematic analysis of new drug indications by drug-gene-disease coherent subnetworks. CPT: Pharmacometrics & Systems Pharmacology 3(11), 146 (2014) 10.1038/psp.2014.44 https://ascpt.onlinelibrary.wiley.com/doi/pdf/10.1038/psp.2014.44

[9] Kim, Y., Cho, Y.-R.: Predicting drug–gene–disease associations by tensor decomposition for network-based computational drug repositioning. Biomedicines 11(7) (2023) 10.3390/biomedicines11071998

[10] Taguchi, Y.-h.: Unsupervised Feature Extraction Applied to Bioinformatics: A PCA Based and TD Based Approach, 1st edn. Springer, ??? (2020). 10.1007/978-3-030-22456-1. http://dx.doi.org/10.1007/978-3-030-22456-1

[11] Taguchi, Y.-h.: Unsupervised Feature Extraction Applied to Bioinformatics: A PCA Based and TD Based Approach, 2nd edn. Springer, ??? (2024)

[12] Taguchi, Y.-H., Turki, T.: Integrated analysis of gene expression and protein–protein interaction with tensor decomposition. Mathematics 11(17) (2023) 10.3390/math11173655

[13] Xie, Z., Bailey, A., Kuleshov, M.V., Clarke, D.J.B., Evangelista, J.E., Jenkins, S.L., Lachmann, A., Wojciechowicz, M.L., Kropiwnicki, E., Jagodnik, K.M., Jeon, M., Ma’ayan, A.: Gene set knowledge discovery with enrichr. Current Protocols 1(3), 90 (2021) 10.1002/cpz1.90 https://currentprotocols.onlinelibrary.wiley.com/doi/pdf/10.1002/cpz1.90

[14] Carr, S., Kasi, A.: Familial Adenomatous Polyposis. StatPearls Publishing, Treasure Island (FL) (2024). Updated 2023 Feb 25. https://www.ncbi.nlm.nih.gov/books/NBK538233/

[15] Aedma, S.K., Kasi, A.: Li-Fraumeni Syndrome. StatPearls Publishing, Treasure Island (FL) (2024). PMID: 30335319. https://pubmed.ncbi.nlm.nih.gov/30335319/

[16] Bhandari, J., Thada, P.K., Puckett, Y.: Fanconi Anemia. StatPearls Publishing, Treasure Island (FL) (2024). Last Update: August 10, 2022. https://www.ncbi.nlm.nih.gov/books/NBK559133/

[17] Zhao, Q., Zhang, Y., Shao, S., Sun, Y., Lin, Z.: Identification of hub genes and biological pathways in hepatocellular carcinoma by integrated bioinformatics analysis. PeerJ 9, 10594 (2021) 10.7717/peerj.10594

[18] Blondel, V.D., Guillaume, J.-L., Lambiotte, R., Lefebvre, E.: Fast unfolding of communities in large networks. Journal of Statistical Mechanics: Theory and Experiment 2008(10), 10008 (2008) 10.1088/1742-5468/2008/10/P10008

[19] Raghavan, U.N., Albert, R., Kumara, S.: Near linear time algorithm to detect community structures in large-scale networks. Phys. Rev. E 76, 036106 (2007) 10.1103/PhysRevE.76.036106

[20] Newman, M.E.J., Girvan, M.: Finding and evaluating community structure in networks. Phys. Rev. E 69, 026113 (2004) 10.1103/PhysRevE.69.026113

[21] Kang, M.-H., Seok Jeong, G., Smoot, D.T., Ashktorab, H., Mo Hwang, C., Sik Kim, B., Sung Kim, H., Park, Y.-Y.: Verteporfin inhibits gastric cancer cell growth by suppressing adhesion molecule fat1. Oncotarget 8(58), 98887–98897 (2017) 10.18632/oncotarget.21946

[22] Rigopoulos, D., Larios, G., Gregoriou, S., Alevizos, A.: Acute and chronic paronychia. Am. Fam. Physician 77(3), 339–346 (2008)

[23] Arjmand Abbassi, Y., Mohammadi, M.T., Sarami Foroshani, M., Raouf Sarshoori, J.: Captopril and valsartan may improve cognitive function through potentiation of the brain antioxidant defense system and attenuation of oxidative/nitrosative damage in stz-induced dementia in rat. Adv Pharm Bull 6(4), 531–539 (2016) 10.15171/apb.2016.067 https://apb.tbzmed.ac.ir/PDF/APB416820160423171936

[24] Michela, C., Antonietta, M., Domenico, T.: Effects of the antidiabetic drugs on the age-related atrophy and sarcopenia associated with diabetes type ii. Current Diabetes Reviews 10(4), 231–237 (2014) 10.2174/1573399810666140918121022

[25] Ohno, T., Nakane, T., Akase, T., Kurasawa, H., Aizawa, Y.: Development of an isogenic human cell trio that models polyglutamine disease. Genes & Genetic Systems 98(4), 179–189 (2023) 10.1266/ggs.22-00030

[26] Giuliani, F., Molica, S., Maiello, E., Battaglia, C., Gebbia, V., Bisceglie, M.D., Vinciarelli, G., Gebbia, N., Colucci, G.: Irinotecan (cpt-11) and mitomycin-c (mmc) as second-line therapy in advanced gastric cancer: A phase ii study of the gruppo oncologico dell’ italia meridionale (prot. 2106). American Journal of Clinical Oncology 28(6), 581–585 (2005) 10.1097/01.coc.0000190398.52142.7f

[27] Zhu, X.-C., Dai, W.-Z., Ma, T.: Overview the effect of statin therapy on dementia risk, cognitive changes and its pathologic change: a systematic review and metaanalysis. Annals of Translational Medicine 6(22) (2018)

[28] Billingsley, E., Vidimos, A.: Paronychia Treatment & Management. Medscape, 1106062 (2022). https://emedicine.medscape.com/article/1106062-treatment?form=fpf

[29] Yoon, J.-H., Yoo, C.-I., Ahn, Y.-S.: N,n-dimethylformamide: evidence of carcinogenicity from national representative cohort study in south korea. Scandinavian Journal of Work, Environment & Health (4), 396–401 (2019) 10.5271/sjweh.3802

[30] Oughtred, R., Rust, J., Chang, C., Breitkreutz, B.-J., Stark, C., Willems, A., Boucher, L., Leung, G., Kolas, N., Zhang, F., Dolma, S., Coulombe-Huntington, J., Chatr-aryamontri, A., Dolinski, K., Tyers, M.: The BioGRID database: A comprehensive biomedical resource of curated protein, genetic, and chemical interactions. Protein Science 30(1), 187–200 (2021) 10.1002/pro.3978 https://onlinelibrary.wiley.com/doi/pdf/10.1002/pro.3978

[31] Xenarios, I., Rice, D.W., Salwinski, L., Baron, M.K., Marcotte, E.M., Eisenberg, D.: DIP: the Database of Interacting Proteins. Nucleic Acids Research 28(1), 289–291 (2000) 10.1093/nar/28.1.289 https://academic.oup.com/nar/article-pdf/28/1/289/9895160/280289.pdf

[32] Bates, D., Maechler, M., Jagan, M.: Matrix: Sparse and Dense Matrix Classes and Methods. (2024). R package version 1. 7-0. https://CRAN.R-project.org/package=Matrix

[33] Baglama, J., Reichel, L., Lewis, B.W.: Irlba: Fast Truncated Singular Value Decomposition and Principal Components Analysis for Large Dense and Sparse Matrices. (2022). R package version 2.3.5.1. https://CRAN.R-project.org/package=irlba

