## Supplementary material for "Novel artificial intelligence-based identification of drug-gene-disease interaction using protein-protein interaction": README.txt

Genes and enrichment analysis
DIP
gene_sym_tensor.ods : using u2i
gene2_sym_tensor.ods : using u3i
genes_sym_PC6.ods   : using u6i
gene_g2_com_eb_1.ods  : 1st cluster using edge-betweennes 
gene_g2_com_eb_2.ods  : 2nd cluster using edge-betweennes            
gene_g2_com_mul_1.ods : 1st cluster using multi-level              
gene_g2_com_mul_2.ods : 2nd cluster using multi-level 
genes_top_50.ods : using top 50 hub proteins
genes_top_200.ods : using top 200 hub proteins

BIOGRID
genes.ods : using u2i
genes2.ods : using u3i
genes_PCA.ods  : using PCA 
genes_top_200.ods  : using top 200 hub proteins
genes_vs_rat.ods : using rat genome

Orthologs between human and mouse
uniprotkb_organism_id_10090_OR_organism_2024_05_17.tsv.gz
