## Supplementary material for "Novel artificial intelligence-based identification of drug-gene-disease interaction using protein-protein interaction": Supplementary_prompt.txt

Find the studies that relate drugs in the drug list and diseases in the disease list: They are not pairwise list. The order is not decided. 

Drug lists: 
(Here comes the list of drugs one per line)

Disease list:
(Here comes the list of diseases one per line)}
