## Supplementary material for "Novel artificial intelligence-based identification of drug-gene-disease interaction using protein-protein interaction": Supplemrntary_Tables.pdf

### Supplementary document Selecting genes that can be used for the identification of drug-gene-disease interaction based upon only protein-protein interaction

Y-h. Taguchi and Turki Turki

Auguset 2024

Table S1: Top 10 diseases in the "Jensen Diseases" category of Enrichr for 217 gene names selected by $u_{2i}$ for BioGrid

| Term | Overlap | P-value | Adjusted P-value |
|---|---|---|---|
| Cancer | 16/300 | $1.93 \times 10^{-7}$ | $4.53 \times 10^{-5}$ |
| Stomach cancer | 5/14 | $2.65 \times 10^{-7}$ | $4.53 \times 10^{-5}$ |
| DOID:9917 | 6/30 | $7.27 \times 10^{-7}$ | $7.77 \times 10^{-5}$ |
| Adenoma | 4/8 | $9.12 \times 10^{-7}$ | $7.77 \times 10^{-5}$ |
| Immune system cancer | 10/137 | $2.54 \times 10^{-6}$ | $1.73 \times 10^{-4}$ |
| Upper respiratory tract disease | 4/13 | $8.93 \times 10^{-6}$ | $5.07 \times 10^{-4}$ |
| Ovarian cancer | 6/51 | $1.82 \times 10^{-5}$ | $8.89 \times 10^{-4}$ |
| Ductal carcinoma in situ | 4/18 | $3.66 \times 10^{-5}$ | $1.48 \times 10^{-3}$ |
| Esophageal carcinoma | 3/7 | $4.27 \times 10^{-5}$ | $1.48 \times 10^{-3}$ |
| Lymphoid leukemia | 9/152 | $4.33 \times 10^{-5}$ | $1.48 \times 10^{-3}$ |

Table S2: Top 10 diseases in the "Jensen Diseases" category of Enrichr for 193 gene names selected by $u_{2i}$ for DIP

| Term | Overlap | P-value | Adjusted P-value |
|---|---|---|---|
| Lymphoid leukemia | 13/152 | $2.51 \times 10^{-9}$ | $8.33 \times 10^{-7}$ |
| Immune system cancer | 11/137 | $8.40 \times 10^{-8}$ | $1.39 \times 10^{-5}$ |
| Cancer | 15/300 | $2.07 \times 10^{-7}$ | $2.29 \times 10^{-5}$ |
| Familial adenomatous polyposis | 5/25 | $3.43 \times 10^{-6}$ | $2.85 \times 10^{-4}$ |
| Upper respiratory tract disease | 4/13 | $5.39 \times 10^{-6}$ | $3.58 \times 10^{-4}$ |
| Biliary tract cancer | 4/14 | $7.49 \times 10^{-6}$ | $4.14 \times 10^{-4}$ |
| Neurodegenerative disease | 11/293 | $1.24 \times 10^{-4}$ | $5.15 \times 10^{-3}$ |
| Ovarian cancer | 5/51 | $1.24 \times 10^{-4}$ | $5.15 \times 10^{-3}$ |
| Systemic lupus erythematosus | 6/83 | $1.44 \times 10^{-4}$ | $5.31 \times 10^{-3}$ |
| Ataxia telangiectasia | 4/30 | $1.82 \times 10^{-4}$ | $6.04 \times 10^{-3}$ |

Table S3: Top 10 diseases in the "Jensen Diseases" category of Enrichr for 57 gene names selected by $u_{3i}$ for DIP

| Term | Overlap | P-value | Adjusted P-value |
|---|---|---|---|
| Li-Fraumeni syndrome | 5/12 | $9.26 \times 10^{-11}$ | $1.38 \times 10^{-8}$ |
| Ovarian cancer | 6/51 | $4.77 \times 10^{-9}$ | $3.56 \times 10^{-7}$ |
| Immune system cancer | 6/137 | $1.82 \times 10^{-6}$ | $8.19 \times 10^{-5}$ |
| Breast cancer | 9/434 | $2.20 \times 10^{-6}$ | $8.19 \times 10^{-5}$ |
| Lymphoid leukemia | 6/152 | $3.34 \times 10^{-6}$ | $9.94 \times 10^{-5}$ |
| Biliary tract cancer | 3/14 | $6.63 \times 10^{-6}$ | $1.65 \times 10^{-4}$ |
| Ductal carcinoma in situ | 3/18 | $1.48 \times 10^{-5}$ | $3.14 \times 10^{-4}$ |
| Nijmegen breakage syndrome | 3/22 | $2.76 \times 10^{-5}$ | $5.15 \times 10^{-4}$ |
| ENSP00000256078 | 3/28 | $5.81 \times 10^{-5}$ | $8.91 \times 10^{-4}$ |
| Hereditary breast ovarian cancer | 2/5 | $7.12 \times 10^{-5}$ | $8.91 \times 10^{-4}$ |

Table S4: Top 10 diseases in the "OMIM Diseases" category of Enrichr for 217 gene names selected by $u_{2i}$ for BioGrid

| Term | Overlap | P-value | Adjusted P-value |
|---|---|---|---|
| Parkinson disease | 3/22 | $1.67 \times 10^{-3}$ | $4.33 \times 10^{-2}$ |
| Breast cancer | 3/28 | $3.38 \times 10^{-3}$ | $4.39 \times 10^{-2}$ |
| Dementia | 2/12 | $7.20 \times 10^{-3}$ | $5.49 \times 10^{-2}$ |
| Ovarian cancer | 2/13 | $8.45 \times 10^{-3}$ | $5.49 \times 10^{-2}$ |
| Thyroid carcinoma | 2/17 | $1.43 \times 10^{-2}$ | $7.44 \times 10^{-2}$ |
| Autism | 2/19 | $1.77 \times 10^{-2}$ | $7.69 \times 10^{-2}$ |
| Prostate cancer | 2/30 | $4.18 \times 10^{-2}$ | $1.55 \times 10^{-1}$ |
| Colorectal cancer | 2/40 | $6.99 \times 10^{-2}$ | $2.27 \times 10^{-1}$ |
| Pancreatic cancer | 1/11 | $1.13 \times 10^{-1}$ | $2.30 \times 10^{-1}$ |
| Gastric cancer | 1/11 | $1.13 \times 10^{-1}$ | $2.30 \times 10^{-1}$ |

Table S5: Top 10 diseases in the "OMIM Diseases" category of Enrichr for 193 gene names selected by $u_{2i}$ for DIP

| Term | Overlap | P-value | Adjusted P-value |
|---|---|---|---|
| Colorectal cancer | 7/40 | $9.27 \times 10^{-8}$ | $2.97 \times 10^{-6}$ |
| Parkinson disease | 5/22 | $1.74 \times 10^{-6}$ | $2.78 \times 10^{-5}$ |
| Ectodermal dysplasia | 3/11 | $1.34 \times 10^{-4}$ | $1.43 \times 10^{-3}$ |
| Breast cancer | 3/28 | $2.36 \times 10^{-3}$ | $1.51 \times 10^{-2}$ |
| ·Immunodeficiency | 3/28 | $2.36 \times 10^{-3}$ | $1.51 \times 10^{-2}$ |
| Fanconi anemia | 2/13 | $6.60 \times 10^{-3}$ | $3.03 \times 10^{-2}$ |
| Leukemia | 4/78 | $6.64 \times 10^{-3}$ | $3.03 \times 10^{-2}$ |
| Myocardial infarction | 2/16 | $9.97 \times 10^{-3}$ | $3.99 \times 10^{-2}$ |
| Orofacial cleft | 2/19 | $1.39 \times 10^{-2}$ | $4.96 \times 10^{-2}$ |
| Lymphoma | 2/22 | $1.85 \times 10^{-2}$ | $5.92 \times 10^{-2}$ |

Table S6: Top 10 diseases in the "OMIM Diseases" category of Enrichr for 57 gene names selected by $u_{3i}$ for DIP.

| Term | Overlap | P-value | Adjusted P-value |
|---|---|---|---|
| Pancreatic cancer | 3/11 | $3.02 \times 10^{-6}$ | $3.13 \times 10^{-5}$ |
| Ovarian cancer | 3/13 | $5.22 \times 10^{-6}$ | $3.13 \times 10^{-5}$ |
| Colorectal cancer | 3/40 | $1.71 \times 10^{-4}$ | $6.85 \times 10^{-4}$ |
| Orofacial cleft | 2/19 | $1.19 \times 10^{-3}$ | $3.56 \times 10^{-3}$ |
| Breast cancer | 2/28 | $2.59 \times 10^{-3}$ | $5.93 \times 10^{-3}$ |
| Prostate cancer | 2/30 | $2.97 \times 10^{-3}$ | $5.93 \times 10^{-3}$ |
| Ectodermal dysplasia | 1/11 | $2.93 \times 10^{-2}$ | $4.61 \times 10^{-2}$ |
| Fanconi anemia | 1/13 | $3.45 \times 10^{-2}$ | $4.61 \times 10^{-2}$ |
| Melanoma | 1/13 | $3.45 \times 10^{-2}$ | $4.61 \times 10^{-2}$ |
| Immunodeficiency | 1/28 | $7.30 \times 10^{-2}$ | $8.75 \times 10^{-2}$ |

Table S7: Top 10 drugs in the "LINCS L1000 Chem Pert Consensus Sigs" category of Enrichr for 217 gene names selected by $u_{2i}$ for BioGrid.

| Term | Overlap | P-value | Adjusted P-value |
|---|---|---|---|
| Nutlin-3 Down | 15/240 | $6.03 \times 10^{-8}$ | $5.18 \times 10^{-4}$ |
| Amisulpride Up | 15/249 | $9.78 \times 10^{-8}$ | $5.18 \times 10^{-4}$ |
| Pyrvinium-Pamoate Down | 14/243 | $4.54 \times 10^{-7}$ | $1.35 \times 10^{-3}$ |
| Actinomycin-D Down | 14/248 | $5.80 \times 10^{-7}$ | $1.35 \times 10^{-3}$ |
| AR-C133057XX Up | 14/250 | $6.39 \times 10^{-7}$ | $1.35 \times 10^{-3}$ |
| CVF-SUMO-11 Up | 13/241 | $2.45 \times 10^{-6}$ | $2.97 \times 10^{-3}$ |
| KPT-330 Down | 13/243 | $2.69 \times 10^{-6}$ | $2.97 \times 10^{-3}$ |
| TAS-103 Down | 13/243 | $2.69 \times 10^{-6}$ | $2.97 \times 10^{-3}$ |
| XL-888 Up | 13/245 | $2.94 \times 10^{-6}$ | $2.97 \times 10^{-3}$ |
| APO-866 Up | 13/246 | $3.08 \times 10^{-6}$ | $2.97 \times 10^{-3}$ |

Table S8: Top 10 drugs in the "LINCS L1000 Chem Pert Consensus Sigs" category of Enrichr for 193 gene names selected by $u_{2i}$ for DIP.

| Term | Overlap | P-value | Adjusted P-value |
|---|---|---|---|
| KU-0060648 Up | 14/236 | $6.59 \times 10^{-8}$ | $3.41 \times 10^{-4}$ |
| Okadaic-Acid Up | 14/246 | $1.10 \times 10^{-7}$ | $3.41 \times 10^{-4}$ |
| Palmatine-Chloride Down | 14/247 | $1.16 \times 10^{-7}$ | $3.41 \times 10^{-4}$ |
| YL-55 Up | 14/249 | $1.28 \times 10^{-7}$ | $3.41 \times 10^{-4}$ |
| CHEMBL-399379 Up | 13/231 | $3.61 \times 10^{-7}$ | $6.87 \times 10^{-4}$ |
| LY-2606368 Up | 13/233 | $3.98 \times 10^{-7}$ | $6.87 \times 10^{-4}$ |
| RS-102221 Down | 13/242 | $6.13 \times 10^{-7}$ | $6.87 \times 10^{-4}$ |
| 4-HQN Down | 13/242 | $6.13 \times 10^{-7}$ | $6.87 \times 10^{-4}$ |
| Butalbital Down | 13/244 | $6.73 \times 10^{-7}$ | $6.87 \times 10^{-4}$ |
| TL-HRAS-61 Up | 13/247 | $7.72 \times 10^{-7}$ | $6.87 \times 10^{-4}$ |

Table S9: Top 10 drugs in the "DSigDB" category of Enrichr for 217 gene names selected by $u_{2i}$ for BioGRID.

| Term | Overlap | P-value | Adjusted P-value |
|---|---|---|---|
| troglitazone CTD 00002415 | 30/651 | $2.31 \times 10^{-11}$ | $6.72 \times 10^{-8}$ |
| Vorinostat CTD 00003560 | 24/425 | $4.48 \times 10^{-11}$ | $6.72 \times 10^{-8}$ |
| MG-132 CTD 00002789 | 15/173 | $7.04 \times 10^{-10}$ | $6.01 \times 10^{-7}$ |
| resveratrol CTD 00002483 | 46/1601 | $8.01 \times 10^{-10}$ | $6.01 \times 10^{-7}$ |
| N-Acetyl-L-cysteine CTD 00005305 | 19/315 | $1.77 \times 10^{-9}$ | $1.06 \times 10^{-6}$ |
| doxorubicin CTD 00005874 | 29/750 | $3.00 \times 10^{-9}$ | $1.50 \times 10^{-6}$ |
| Bortezomib CTD 00003736 | 30/878 | $2.61 \times 10^{-8}$ | $1.12 \times 10^{-5}$ |
| etoposide CTD 00005948 | 21/461 | $3.43 \times 10^{-8}$ | $1.15 \times 10^{-5}$ |
| piroxicam CTD 00006571 | 23/549 | $3.45 \times 10^{-8}$ | $1.15 \times 10^{-5}$ |
| Arsenenous acid CTD 00000922 | 37/1283 | $4.47 \times 10^{-8}$ | $1.34 \times 10^{-5}$ |

Table S10: Top 10 drugs in the "DSigDB" category of Enrichr for 193 gene names selected by $u_{2i}$ for DIP.

| Term | Overlap | P-value | Adjusted P-value |
|---|---|---|---|
| curcumin CTD 00000663 | 34/528 | $8.71 \times 10^{-19}$ | $2.63 \times 10^{-15}$ |
| doxorubicin CTD 00005874 | 38/750 | $2.02 \times 10^{-17}$ | $3.05 \times 10^{-14}$ |
| N-Acetyl-L-cysteine CTD 00005305 | 25/315 | $4.17 \times 10^{-16}$ | $4.19 \times 10^{-13}$ |
| Arsenenous acid CTD 00000922 | 47/1283 | $6.07 \times 10^{-16}$ | $4.58 \times 10^{-13}$ |
| MG-132 CTD 00002789 | 19/173 | $4.61 \times 10^{-15}$ | $2.78 \times 10^{-12}$ |
| wortmannin CTD 00000504 | 17/146 | $5.05 \times 10^{-14}$ | $2.54 \times 10^{-11}$ |
| LY 294002 CTD 00003061 | 20/227 | $6.01 \times 10^{-14}$ | $2.59 \times 10^{-11}$ |
| ZINC CTD 00007011 | 50/1642 | $8.50 \times 10^{-14}$ | $3.21 \times 10^{-11}$ |
| 5-Fluorouracil CTD 00005987 | 42/1202 | $1.48 \times 10^{-13}$ | $4.96 \times 10^{-11}$ |
| etoposide CTD 00005948 | 25/461 | $2.38 \times 10^{-12}$ | $7.18 \times 10^{-10}$ |

Table S11: Top 10 drugs in the "DSigDB" category of Enrichr for 57 gene names selected by $u_{3i}$ for DIP.

| Term | Overlap | P-value | Adjusted P-value |
|---|---|---|---|
| Vorinostat CTD 00003560 | 23/425 | $1.11 \times 10^{-24}$ | $2.69 \times 10^{-21}$ |
| FENRETINIDE CTD 00007166 | 16/228 | $6.88 \times 10^{-19}$ | $8.36 \times 10^{-16}$ |
| curcumin CTD 00000663 | 20/528 | $2.65 \times 10^{-18}$ | $2.15 \times 10^{-15}$ |
| resveratrol CTD 00002483 | 28/1601 | $3.80 \times 10^{-17}$ | $2.31 \times 10^{-14}$ |
| doxorubicin CTD 00005874 | 21/750 | $1.40 \times 10^{-16}$ | $6.79 \times 10^{-14}$ |
| aspirin CTD 00005447 | 19/561 | $1.77 \times 10^{-16}$ | $7.17 \times 10^{-14}$ |
| bay 11-7082 CTD 00003959 | 11/83 | $2.62 \times 10^{-16}$ | $9.09 \times 10^{-14}$ |
| Arsenenous acid CTD 00000922 | 25/1283 | $3.34 \times 10^{-16}$ | $1.02 \times 10^{-13}$ |
| tamoxifen CTD 00006827 | 20/802 | $8.12 \times 10^{-15}$ | $2.12 \times 10^{-12}$ |
| Leptomycin B CTD 00001805 | 9/50 | $8.70 \times 10^{-15}$ | $2.12 \times 10^{-12}$ |

Table S12: Top 10 drugs in the "DrugMatrix" category of Enrichr for 217 gene names selected by $u_{2i}$ for BioGRID.

| Term | Overlap | P-value | Adjusted P-value |
|---|---|---|---|
| Azauridine-1500 mg/kg in CMC-Rat-Bone marrow-3d-up | 16/311 | $3.13 \times 10^{-7}$ | $1.89 \times 10^{-3}$ |
| Lead (II) Acetate-200 mg/kg in Saline-Rat-Spleen-0.25d-up | 16/330 | $6.91 \times 10^{-7}$ | $1.89 \times 10^{-3}$ |
| Famciclovir-112 mg/kg in Saline-Rat-Bone marrow-1d-dn | 15/291 | $7.23 \times 10^{-7}$ | $1.89 \times 10^{-3}$ |
| 44'-Methylenedianiline-296 uM in DMSO-Rat-Primary rat hepatocytes-0.67d-up | 15/304 | $1.25 \times 10^{-6}$ | $2.45 \times 10^{-3}$ |
| Rabeprazole-1024 mg/kg in Water-Rat-Liver-1d-up | 14/279 | $2.34 \times 10^{-6}$ | $3.43 \times 10^{-3}$ |
| Carbon Tetrachloride-3178 mg/kg in Saline-Rat-Liver-1d-up | 15/326 | $2.96 \times 10^{-6}$ | $3.43 \times 10^{-3}$ |
| Diethylstilbestrol-280 mg/kg in Corn Oil-Rat-Liver-1d-up | 14/290 | $3.67 \times 10^{-6}$ | $3.43 \times 10^{-3}$ |
| Clobetasol Propionate-17 mg/kg in Corn Oil-Rat-Bone marrow-3d-dn | 14/294 | $4.30 \times 10^{-6}$ | $3.43 \times 10^{-3}$ |
| 6-Mercaptopurine-25 mg/kg in Corn Oil-Rat-Liver-0.25d-up | 14/299 | $5.22 \times 10^{-6}$ | $3.43 \times 10^{-3}$ |
| Cyclosporin A-70 mg/kg in Corn Oil-Rat-Spleen-0.25d-up | 15/347 | $6.31 \times 10^{-6}$ | $3.43 \times 10^{-3}$ |

Table S13: Top 10 drugs in the "DrugMatrix" category of Enrichr for 193 gene names selected by $u_{2i}$ for DIP .

| Term | Overlap | P-value | Adjusted P-value |
|---|---|---|---|
| Ceramide C2-44 uM in DMSO-Rat-Primary rat hepatocytes-0.67d-up | 16/311 | $5.40 \times 10^{-8}$ | $4.14 \times 10^{-4}$ |
| Ceramide C2-44 uM in DMSO-Rat-Primary rat hepatocytes-1d-up | 13/280 | $3.12 \times 10^{-6}$ | $1.19 \times 10^{-2}$ |
| Cerivastatin-7 mg/kg in Corn Oil-Rat-Skeletal muscle-5d-up | 12/274 | $1.35 \times 10^{-5}$ | $3.46 \times 10^{-2}$ |
| Methyl Salicylate-444 mg/kg in Corn Oil-Rat-Kidney-3d-up | 12/290 | $2.37 \times 10^{-5}$ | $4.37 \times 10^{-2}$ |
| Lithocholic Acid-74 uM in DMSO-Rat-Primary rat hepatocytes-0.67d-up | 12/300 | $3.31 \times 10^{-5}$ | $4.37 \times 10^{-2}$ |
| Stavudine-1120 uM in DMSO-Rat-Primary rat hepatocytes-0.67d-up | 12/301 | $3.42 \times 10^{-5}$ | $4.37 \times 10^{-2}$ |
| Carbon Tetrachloride-400 mg/kg in Corn Oil-Rat-Liver-0.25d-up | 11/280 | $8.30 \times 10^{-5}$ | $9.09 \times 10^{-2}$ |
| Microcystin-LR-0.07 uM in DMSO-Rat-Primary rat hepatocytes-0.67d-up | 11/309 | $1.97 \times 10^{-4}$ | $1.66 \times 10^{-1}$ |
| Pantoprazole-650 uM in DMSO-Rat-Primary rat hepatocytes-0.67d-up | 11/310 | $2.03 \times 10^{-4}$ | $1.66 \times 10^{-1}$ |
| Daunorubicin-1.5 uM in DMSO-Rat-Primary rat hepatocytes-1d-up | 11/324 | $2.96 \times 10^{-4}$ | $1.66 \times 10^{-1}$ |

Table S14: Top 10 drugs in the "Drug Perturbations from GEO down" category of Enrichr for 217 gene names selected by $u_{2i}$ for BioGRID.

| Term | Overlap | P-value | Adjusted P-value |
|---|---|---|---|
| cytarabine, EC50, 1 d 6253 human GSE6930 sample 3373 | 24/339 | $3.73 \times 10^{-13}$ | $3.14 \times 10^{-10}$ |
| ARC (NSC 188491) 302576 human GSE13477 sample 2503 | 22/329 | $1.16 \times 10^{-11}$ | $4.16 \times 10^{-9}$ |
| doxorubicin, EC50, 1 d 31703 human GSE6930 sample 3260 | 21/300 | $1.48 \times 10^{-11}$ | $4.16 \times 10^{-9}$ |
| lapatinib DB01259 human GSE38376 sample 2586 | 20/291 | $6.43 \times 10^{-11}$ | $1.35 \times 10^{-8}$ |
| lapatinib DB01259 human GSE38376 sample 2585 | 20/301 | $1.18 \times 10^{-10}$ | $1.98 \times 10^{-8}$ |
| EPZ004777 56962336 human GSE29828 sample 2650 | 22/376 | $1.54 \times 10^{-10}$ | $2.17 \times 10^{-8}$ |
| sangivamycin (NSC 65346) 14978 human GSE13477 sample 2504 | 20/330 | $5.95 \times 10^{-10}$ | $7.17 \times 10^{-8}$ |
| bexarotene DB00307 human GSE6914 sample 2680 | 23/453 | $9.32 \times 10^{-10}$ | $9.82 \times 10^{-8}$ |
| cytarabine, EC50, 5 d 6253 human GSE6930 sample 3375 | 23/468 | $1.74 \times 10^{-9}$ | $1.63 \times 10^{-7}$ |
| BPDE 41322 human GSE19510 sample 3377 | 17/274 | $8.60 \times 10^{-9}$ | $7.25 \times 10^{-7}$ |

Table S15: Top 10 drugs in the "Drug Perturbations from GEO down" category of Enrichr for 193 gene names selected by $u_{2i}$ for DIP.

| Term | Overlap | P-value | Adjusted P-value |
|---|---|---|---|
| dactinomycin DB00970 mouse GSE5324 sample 2701 | 22/522 | $6.08 \times 10^{-9}$ | $5.20 \times 10^{-6}$ |
| dactinomycin DB00970 mouse GSE5324 sample 2700 | 22/544 | $1.28 \times 10^{-8}$ | $5.48 \times 10^{-6}$ |
| cisplatin DB00515 human GSE6410 sample 2532 | 15/315 | $3.86 \times 10^{-7}$ | $1.10 \times 10^{-4}$ |
| SODIUM BUTYRATE 5222465 mouse GSE4410 sample 3582 | 14/289 | $7.78 \times 10^{-7}$ | $1.33 \times 10^{-4}$ |
| Etanercept DB00005 human GSE7524 sample 3295 | 16/379 | $7.80 \times 10^{-7}$ | $1.33 \times 10^{-4}$ |
| Curcumin CID 969516 human GSE10896 sample 3259 | 17/458 | $2.04 \times 10^{-6}$ | $2.91 \times 10^{-4}$ |
| actinomycin D 2019 human GSE6400 sample 3098 | 14/323 | $2.88 \times 10^{-6}$ | $3.52 \times 10^{-4}$ |
| SODIUM BUTYRATE 5222465 mouse GSE4410 sample 3584 | 13/343 | $2.72 \times 10^{-5}$ | $2.91 \times 10^{-3}$ |
| lapatinib DB01259 human GSE38376 sample 2585 | 12/301 | $3.42 \times 10^{-5}$ | $3.25 \times 10^{-3}$ |
| HYPOCHLOROUS ACID 24341 human GSE11630 sample 3202 | 11/289 | $1.10 \times 10^{-4}$ | $9.07 \times 10^{-3}$ |

Table S16: Top 10 drugs in the "Drug Perturbations from GEO down" category of Enrichr for 57 gene names selected by $u_{3i}$ for DIP.

| Term | Overlap | P-value | Adjusted P-value |
|---|---|---|---|
| trastuzumab DB00072 human GSE31432 sample 3123 | 7/280 | $9.87 \times 10^{-6}$ | $2.69 \times 10^{-3}$ |
| ZINC ACETATE 11192 human GSE2964 sample 3589 | 6/189 | $1.17 \times 10^{-5}$ | $2.69 \times 10^{-3}$ |
| lapatinib DB01259 human GSE38376 sample 2585 | 7/301 | $1.58 \times 10^{-5}$ | $2.69 \times 10^{-3}$ |
| doxycycline DB00254 human GSE2624 sample 3074 | 8/425 | $1.72 \times 10^{-5}$ | $2.69 \times 10^{-3}$ |
| genistein 5280961 mouse GSE2889 sample 3606 | 6/247 | $5.25 \times 10^{-5}$ | $6.57 \times 10^{-3}$ |
| doxycycline DB00254 human GSE2624 sample 3077 | 7/391 | $8.30 \times 10^{-5}$ | $8.37 \times 10^{-3}$ |
| 2,3,7,8-tetrachlorodibenzo-p-dioxin 10852289 mouse GSE2812 sample 3602 | 7/400 | $9.57 \times 10^{-5}$ | $8.37 \times 10^{-3}$ |
| Bisphenol A 6623 human GSE17624 sample 2654 | 6/281 | $1.07 \times 10^{-4}$ | $8.37 \times 10^{-3}$ |
| lapatinib DB01259 human GSE38376 sample 2586 | 6/291 | $1.29 \times 10^{-4}$ | $9.00 \times 10^{-3}$ |
| estradiol 5757 mouse GSE2889 sample 3609 | 7/433 | $1.56 \times 10^{-4}$ | $9.78 \times 10^{-3}$ |

Table S17: Top 10 drugs in the "Drug Perturbations from GEO up" category of Enrichr for 217 gene names selected by $u_{2i}$ for BioGRID.

| Term | Overlap | P-value | Adjusted P-value |
|---|---|---|---|
| Dmnq (2,3-Dimethoxy-1,4-naphthoquinone) 3136 human GSE6907 sample 3527 | 20/321 | $3.67 \times 10^{-10}$ | $1.68 \times 10^{-7}$ |
| Arachidonic acid DB04557 human GSE3737 sample 3171 | 22/395 | $3.93 \times 10^{-10}$ | $1.68 \times 10^{-7}$ |
| NICKEL 935 human GSE6907 sample 3531 | 18/288 | $2.75 \times 10^{-9}$ | $6.94 \times 10^{-7}$ |
| MT19c compound SID 134222379 human GSE23616 sample 3352 | 18/291 | $3.24 \times 10^{-9}$ | $6.94 \times 10^{-7}$ |
| quercetin 5280343 human GSE7259 sample 3415 | 19/336 | $5.10 \times 10^{-9}$ | $8.76 \times 10^{-7}$ |
| ARSENIC 5359596 human GSE6907 sample 3529 | 20/394 | $1.24 \times 10^{-8}$ | $1.77 \times 10^{-6}$ |
| Triiodothyronine-[13C6] hydrochloride (T3 thyronine) 53442265 human GSE29159 sample 3042 | 19/362 | $1.71 \times 10^{-8}$ | $1.94 \times 10^{-6}$ |
| MLN4924 73014187 human GSE30531 sample 2644 | 18/325 | $1.81 \times 10^{-8}$ | $1.94 \times 10^{-6}$ |
| nicotine 942 human GSE6264 sample 3064 | 19/370 | $2.42 \times 10^{-8}$ | $2.16 \times 10^{-6}$ |
| Triiodothyronine-[13C6] hydrochloride (T3 thyronine) 53442265 human GSE29159 sample 3045 | 18/332 | $2.51 \times 10^{-8}$ | $2.16 \times 10^{-6}$ |

Table S18: Top 10 drugs in the "Drug Perturbations from GEO up" category of Enrichr for 193 gene names selected by $u_{2i}$ for DIP.

| Term | Overlap | P-value | Adjusted P-value |
|---|---|---|---|
| bortezomib DB00188 human GSE30931 sample 2686 | 18/364 | $1.41 \times 10^{-8}$ | $1.23 \times 10^{-5}$ |
| Triiodothyronine-[13C6] hydrochloride (T3 thyronine) 53442265 human GSE29159 sample 3045 | 15/332 | $7.50 \times 10^{-7}$ | $2.53 \times 10^{-4}$ |
| phenol 996 human GSE6907 sample 3528 | 15/336 | $8.72 \times 10^{-7}$ | $2.53 \times 10^{-4}$ |
| resveratrol DB02709 human GSE36930 sample 3498 | 15/349 | $1.40 \times 10^{-6}$ | $2.96 \times 10^{-4}$ |
| Promegestone 36709 human GSE67561 sample 3694 | 13/265 | $1.70 \times 10^{-6}$ | $2.96 \times 10^{-4}$ |
| estradiol 5757 mouse GSE2195 sample 3619 | 13/271 | $2.17 \times 10^{-6}$ | $3.16 \times 10^{-4}$ |
| Bisphenol A 6623 mouse GSE4650 sample 3574 | 11/202 | $4.02 \times 10^{-6}$ | $5.01 \times 10^{-4}$ |
| pioglitazone DB01132 rat GSE20219 sample 2794 | 13/292 | $4.92 \times 10^{-6}$ | $5.37 \times 10^{-4}$ |
| Triiodothyronine-[13C6] hydrochloride (T3 thyronine) 53442265 human GSE29159 sample 3044 | 13/300 | $6.60 \times 10^{-6}$ | $6.39 \times 10^{-4}$ |
| NVP-TNKS656 (tankyrase inhibitor) 71227201 human GSE55624 sample 3247 | 13/307 | $8.45 \times 10^{-6}$ | $7.37 \times 10^{-4}$ |

Table S19: Top 10 drugs in the "Drug Perturbations from GEO up" category of Enrichr for 57 gene names selected by $u_{3i}$ for DIP.

| Term | Overlap | P-value | Adjusted P-value |
|---|---|---|---|
| cisplatin DB00515 mouse GSE6206 sample 3412 | 8/279 | $7.74 \times 10^{-7}$ | $3.96 \times 10^{-4}$ |
| Bisphenol A 6623 mouse GSE4650 sample 3574 | 7/202 | $1.14 \times 10^{-6}$ | $3.96 \times 10^{-4}$ |
| lipopolysaccharide (LPS) 11970143 mouse GSE23639 sample 3443 | 7/261 | $6.24 \times 10^{-6}$ | $1.44 \times 10^{-3}$ |
| Mehp 20393 rat GSE4514 sample 3578 | 7/293 | $1.32 \times 10^{-5}$ | $1.84 \times 10^{-3}$ |
| Hydrogen Peroxide 784 mouse GSE3078 sample 3599 | 6/199 | $1.56 \times 10^{-5}$ | $1.84 \times 10^{-3}$ |
| levetiracetam 5284583 rat GSE2880 sample 2669 | 6/201 | $1.65 \times 10^{-5}$ | $1.84 \times 10^{-3}$ |
| 2,3,7,8-tetrachlorodibenzo-p-dioxin 10852289 human GSE24193 sample 3189 | 7/309 | $1.87 \times 10^{-5}$ | $1.84 \times 10^{-3}$ |
| Triiodothyronine-[13C6] hydrochloride (T3 thyronine) 53442265 human GSE29159 sample 3043 | 7/320 | $2.34 \times 10^{-5}$ | $2.02 \times 10^{-3}$ |
| Bisphenol A 6623 mouse GSE4650 sample 3575 | 6/225 | $3.12 \times 10^{-5}$ | $2.40 \times 10^{-3}$ |
| Phosgene 6371 mouse GSE2565 sample 3611 | 6/243 | $4.80 \times 10^{-5}$ | $3.32 \times 10^{-3}$ |

Table S20: Top ranked 10 diseases in "Jensen DISEASES" category of Enrichr for top 200 hub proteins for BioGRID.

| Term | Overlap | P-value | Adjusted P-value |
|---|---|---|---|
| Stomach cancer | 6/14 | $5.87 \times 10^{-9}$ | $1.57 \times 10^{-6}$ |
| Adenoma | 5/8 | $1.03 \times 10^{-8}$ | $1.57 \times 10^{-6}$ |
| Cancer | 18/300 | $1.22 \times 10^{-8}$ | $1.57 \times 10^{-6}$ |
| DOID:9917 | 7/30 | $3.83 \times 10^{-8}$ | $3.70 \times 10^{-6}$ |
| Immune system cancer | 12/137 | $6.01 \times 10^{-8}$ | $4.64 \times 10^{-6}$ |
| Esophageal carcinoma | 4/7 | $5.70 \times 10^{-7}$ | $3.67 \times 10^{-5}$ |
| Toxic encephalopathy | 10/126 | $1.92 \times 10^{-6}$ | $1.06 \times 10^{-4}$ |
| Intestinal benign neoplasm | 4/11 | $5.19 \times 10^{-6}$ | $2.50 \times 10^{-4}$ |
| Dementia | 9/119 | $9.39 \times 10^{-6}$ | $3.55 \times 10^{-4}$ |
| Lymphoid leukemia | 10/152 | $1.03 \times 10^{-5}$ | $3.55 \times 10^{-4}$ |

Table S21: Top ranked 10 diseases in "OMIM Diseases" category of Enrichr for top 200 hub proteins for BioGRID.

| Term | Overlap | P-value | Adjusted P-value |
|---|---|---|---|
| Dementia | 4/12 | $7.71 \times 10^{-6}$ | $2.70 \times 10^{-4}$ |
| Ovarian cancer | 3/13 | $3.89 \times 10^{-4}$ | $6.81 \times 10^{-3}$ |
| Parkinson disease | 3/22 | $1.94 \times 10^{-3}$ | $2.27 \times 10^{-2}$ |
| Breast cancer | 3/28 | $3.93 \times 10^{-3}$ | $3.44 \times 10^{-2}$ |
| Colorectal cancer | 3/40 | $1.07 \times 10^{-2}$ | $7.50 \times 10^{-2}$ |
| Thyroid carcinoma | 2/17 | $1.59 \times 10^{-2}$ | $9.25 \times 10^{-2}$ |
| Autism | 2/19 | $1.96 \times 10^{-2}$ | $9.82 \times 10^{-2}$ |
| Prostate cancer | 2/30 | $4.60 \times 10^{-2}$ | $1.91 \times 10^{-1}$ |
| Muscular dystrophy | 2/33 | $5.46 \times 10^{-2}$ | $1.91 \times 10^{-1}$ |
| Cardiomyopathy, dilated | 2/33 | $5.46 \times 10^{-2}$ | $1.91 \times 10^{-1}$ |

Table S22: Top ranked 10 drugs in "LINCS L1000 Chem Pert Consensus Sigs" category of Enrichr for top 200 hub proteins for BioGRID.

| Term | Overlap | P-value | Adjusted P-value |
|---|---|---|---|
| Pralatrexate Down | 20/249 | $1.02 \times 10^{-11}$ | $1.09 \times 10^{-7}$ |
| Gemcitabine Down | 19/239 | $4.09 \times 10^{-11}$ | $2.06 \times 10^{-7}$ |
| Etoposide Down | 19/244 | $5.86 \times 10^{-11}$ | $2.06 \times 10^{-7}$ |
| Levothyroxine Up | 19/248 | $7.75 \times 10^{-11}$ | $2.06 \times 10^{-7}$ |
| Aphidicolin Down | 18/241 | $3.76 \times 10^{-10}$ | $7.48 \times 10^{-7}$ |
| TAS-103 Down | 18/243 | $4.30 \times 10^{-10}$ | $7.48 \times 10^{-7}$ |
| N-Bromoacetyltryptamine Up | 18/245 | $4.91 \times 10^{-10}$ | $7.48 \times 10^{-7}$ |
| GW-9662 Up | 18/248 | $5.98 \times 10^{-10}$ | $7.97 \times 10^{-7}$ |
| Clofarabine Down | 17/239 | $2.48 \times 10^{-9}$ | $2.94 \times 10^{-6}$ |
| BX-795 Down | 17/241 | $2.81 \times 10^{-9}$ | $3.00 \times 10^{-6}$ |

Table S23: Top ranked 10 drugs in "DSigDB" category of Enrichr for top 200 hub proteins for BioGRID.

| Term | Overlap | P-value | Adjusted P-value |
|---|---|---|---|
| Vorinostat CTD 00003560 | 35/425 | $4.12 \times 10^{-20}$ | $1.33 \times 10^{-16}$ |
| troglitazone CTD 00002415 | 41/651 | $3.68 \times 10^{-19}$ | $5.97 \times 10^{-16}$ |
| doxorubicin CTD 00005874 | 43/750 | $1.51 \times 10^{-18}$ | $1.63 \times 10^{-15}$ |
| resveratrol CTD 00002483 | 60/1601 | $8.26 \times 10^{-17}$ | $6.69 \times 10^{-14}$ |
| Arsenenous acid CTD 00000922 | 53/1283 | $1.43 \times 10^{-16}$ | $9.30 \times 10^{-14}$ |
| MG-132 CTD 00002789 | 21/173 | $8.11 \times 10^{-16}$ | $4.38 \times 10^{-13}$ |
| 2-Bromo-3-hydroxy-4-methoxybenzaldehyde CTD 00004641 | 18/128 | $7.28 \times 10^{-15}$ | $3.37 \times 10^{-12}$ |
| clindamycin HL60 DOWN | 56/1626 | $4.30 \times 10^{-14}$ | $1.74 \times 10^{-11}$ |
| Bortezomib CTD 00003736 | 40/878 | $6.22 \times 10^{-14}$ | $2.19 \times 10^{-11}$ |
| aspirin CTD 00005447 | 32/561 | $6.75 \times 10^{-14}$ | $2.19 \times 10^{-11}$ |

Table S24: Top ranked 10 drugs in "DrugMatrix" category of Enrichr for top 200 hub proteins for BioGRID.

| Term | Overlap | P-value | Adjusted P-value |
|---|---|---|---|
| Cyclosporin A, 70 mg/kg, Corn Oil, Rat, Spleen, 0.25d-up | 21/347 | $6.02 \times 10^{-10}$ | $4.73 \times 10^{-6}$ |
| Azathioprine, 20 mg/kg, Corn Oil, Rat, Spleen, 0.25d-up | 20/335 | $1.99 \times 10^{-9}$ | $5.25 \times 10^{-6}$ |
| NN-Dimethylformamide, 140 mg/kg, Saline, Rat, Spleen, 3d-dn | 20/345 | $3.29 \times 10^{-9}$ | $5.25 \times 10^{-6}$ |
| Podophyllotoxin, 4 mg/kg, Corn Oil, Rat, Bone marrow, 3d-up | 20/346 | $3.46 \times 10^{-9}$ | $5.25 \times 10^{-6}$ |
| Mitomycin C, 0.5 mg/kg, Saline, Rat, Bone marrow, 0.25d-dn | 19/311 | $3.51 \times 10^{-9}$ | $5.25 \times 10^{-6}$ |
| Diethylstilbestrol, 2.8 mg/kg, Corn Oil, Rat, Spleen, 0.25d-up | 20/349 | $4.01 \times 10^{-9}$ | $5.25 \times 10^{-6}$ |
| Cisplatin, 2 mg/kg, Saline, Rat, Bone marrow, 1d-dn | 18/300 | $1.22 \times 10^{-8}$ | $1.37 \times 10^{-5}$ |
| Leucovorin, 1500 mg/kg, CMC, Rat, Bone marrow, 1d-up | 17/289 | $4.19 \times 10^{-8}$ | $3.46 \times 10^{-5}$ |
| Doxapram, 20 mg/kg, Saline, Rat, Heart, 1d-up | 18/326 | $4.36 \times 10^{-8}$ | $3.46 \times 10^{-5}$ |
| Famciclovir, 112 mg/kg, Saline, Rat, Bone marrow, 1d-dn | 17/291 | $4.63 \times 10^{-8}$ | $3.46 \times 10^{-5}$ |

Table S25: Top ranked 10 drugs in "Drug Perturbations from GEO down" category of Enrichr for top 200 hub proteins for BioGRID.

| Term | Overlap | P-value | Adjusted P-value |
|---|---|---|---|
| apratoxin A 6326668 human GSE2742 sample 3067 | 30/355 | $1.17 \times 10^{-17}$ | $9.98 \times 10^{-15}$ |
| imatinib (glivec) 123596 human GSE12211 sample 2518 | 32/442 | $8.20 \times 10^{-17}$ | $3.48 \times 10^{-14}$ |
| doxycycline DB00254 human GSE2624 sample 3074 | 31/425 | $2.10 \times 10^{-16}$ | $5.96 \times 10^{-14}$ |
| cisplatin DB00515 human GSE6410 sample 2532 | 27/315 | $3.89 \times 10^{-16}$ | $8.26 \times 10^{-14}$ |
| apratoxin A 6326668 human GSE2742 sample 3071 | 29/389 | $1.19 \times 10^{-15}$ | $2.03 \times 10^{-13}$ |
| cytarabine, EC50, 1 d 6253 human GSE6930 sample 3373 | 27/339 | $2.42 \times 10^{-15}$ | $3.43 \times 10^{-13}$ |
| lapatinib DB01259 human GSE38376 sample 2585 | 25/301 | $1.06 \times 10^{-14}$ | $1.22 \times 10^{-12}$ |
| BPDE 41322 human GSE19510 sample 3377 | 24/274 | $1.14 \times 10^{-14}$ | $1.22 \times 10^{-12}$ |
| amoxicillin DB01060 rat GSE2354 sample 2678 | 29/431 | $1.71 \times 10^{-14}$ | $1.62 \times 10^{-12}$ |
| estradiol 5757 mouse GSE2889 sample 3609 | 29/433 | $1.93 \times 10^{-14}$ | $1.64 \times 10^{-12}$ |

Table S26: Top ranked 10 drugs in "Drug Perturbations from GEO up" category of Enrichr for top 200 hub proteins for BioGRID.

| Term | Overlap | P-value | Adjusted P-value |
|---|---|---|---|
| apratoxin A 6326668 human GSE2742 sample 3068 | 31/389 | $1.73 \times 10^{-17}$ | $1.49 \times 10^{-14}$ |
| MT19c compound SID 134222379 human GSE23616 sample 3352 | 27/291 | $5.26 \times 10^{-17}$ | $2.27 \times 10^{-14}$ |
| estradiol DB00783 human GSE1153 sample 2734 | 29/388 | $1.11 \times 10^{-15}$ | $3.20 \times 10^{-13}$ |
| estradiol 5757 human GSE4668 sample 3063 | 28/367 | $2.13 \times 10^{-15}$ | $4.59 \times 10^{-13}$ |
| quercetin 5280343 human GSE7259 sample 3416 | 25/327 | $7.07 \times 10^{-14}$ | $1.22 \times 10^{-11}$ |
| ZINC ACETATE 11192 human GSE2964 sample 3590 | 20/217 | $7.97 \times 10^{-13}$ | $1.15 \times 10^{-10}$ |
| quercetin 5280343 human GSE7259 sample 3415 | 24/336 | $1.00 \times 10^{-12}$ | $1.22 \times 10^{-10}$ |
| nicotine 942 human GSE6264 sample 3064 | 25/370 | $1.13 \times 10^{-12}$ | $1.22 \times 10^{-10}$ |
| NICKEL 935 human GSE6907 sample 3531 | 22/288 | $2.45 \times 10^{-12}$ | $2.35 \times 10^{-10}$ |
| Arachidonic acid DB04557 human GSE3737 sample 3171 | 25/395 | $4.77 \times 10^{-12}$ | $4.11 \times 10^{-10}$ |

Table S27: Top ranked 10 diseases in "Jensen DISEASES" category of Enrichr for top 200 hub proteins for DIP.

| Term | Overlap | P-value | Adjusted P-value |
|---|---|---|---|
| Cancer | 28/300 | $8.22 \times 10^{-20}$ | $4.05 \times 10^{-17}$ |
| Toxic encephalopathy | 13/126 | $2.43 \times 10^{-10}$ | $5.98 \times 10^{-8}$ |
| Hyperglycemia | 11/108 | $6.96 \times 10^{-9}$ | $1.14 \times 10^{-6}$ |
| Lung disease | 11/119 | $1.94 \times 10^{-8}$ | $2.39 \times 10^{-6}$ |
| Lymphoid leukemia | 12/152 | $2.58 \times 10^{-8}$ | $2.55 \times 10^{-6}$ |
| Intestinal benign neoplasm | 5/11 | $3.32 \times 10^{-8}$ | $2.73 \times 10^{-6}$ |
| Immune system cancer | 11/137 | $8.40 \times 10^{-8}$ | $5.62 \times 10^{-6}$ |
| Upper respiratory tract disease | 5/13 | $9.12 \times 10^{-8}$ | $5.62 \times 10^{-6}$ |
| Biliary tract cancer | 5/14 | $1.41 \times 10^{-7}$ | $6.94 \times 10^{-6}$ |
| Stomach cancer | 5/14 | $1.41 \times 10^{-7}$ | $6.94 \times 10^{-6}$ |

Table S28: Top ranked 10 diseases in "OMIM Diseases" category of Enrichr for top 200 hub proteins for DIP.

| Term | Overlap | P-value | Adjusted P-value |
|---|---|---|---|
| colorectal cancer | 7/40 | $9.27 \times 10^{-8}$ | $3.52 \times 10^{-6}$ |
| ovarian cancer | 4/13 | $5.39 \times 10^{-6}$ | $7.85 \times 10^{-5}$ |
| breast cancer | 5/28 | $6.20 \times 10^{-6}$ | $7.85 \times 10^{-5}$ |
| leukemia | 7/78 | $9.67 \times 10^{-6}$ | $9.19 \times 10^{-5}$ |
| lung cancer | 4/21 | $4.25 \times 10^{-5}$ | $3.23 \times 10^{-4}$ |
| dementia | 3/12 | $1.77 \times 10^{-4}$ | $1.12 \times 10^{-3}$ |
| melanoma | 3/13 | $2.29 \times 10^{-4}$ | $1.24 \times 10^{-3}$ |
| orofacial cleft | 3/19 | $7.42 \times 10^{-4}$ | $3.53 \times 10^{-3}$ |
| pancreatic cancer | 2/11 | $4.71 \times 10^{-3}$ | $1.79 \times 10^{-2}$ |
| ectodermal dysplasia | 2/11 | $4.71 \times 10^{-3}$ | $1.79 \times 10^{-2}$ |

Table S29: Top ranked 10 drugs in "LINCS L1000 Chem Pert Consensus Sigs" category of Enrichr for top 200 hub proteins for DIP.

| Term | Overlap | P-value | Adjusted P-value |
|---|---|---|---|
| Paracetamol Up | 13/244 | $6.73 \times 10^{-7}$ | $2.99 \times 10^{-3}$ |
| PRIMA1 Up | 13/247 | $7.72 \times 10^{-7}$ | $2.99 \times 10^{-3}$ |
| I-070759 Down | 13/249 | $8.46 \times 10^{-7}$ | $2.99 \times 10^{-3}$ |
| VU-0415556-1 Down | 12/240 | $3.53 \times 10^{-6}$ | $6.32 \times 10^{-3}$ |
| Teniposide Up | 12/245 | $4.36 \times 10^{-6}$ | $6.32 \times 10^{-3}$ |
| GALR2 M617 Up | 12/246 | $4.55 \times 10^{-6}$ | $6.32 \times 10^{-3}$ |
| BCI-hydrochloride Down | 12/246 | $4.55 \times 10^{-6}$ | $6.32 \times 10^{-3}$ |
| Dactinomycin Down | 12/248 | $4.94 \times 10^{-6}$ | $6.32 \times 10^{-3}$ |
| Methiazole Up | 12/250 | $5.37 \times 10^{-6}$ | $6.32 \times 10^{-3}$ |
| Anguidine Up | 11/237 | $1.82 \times 10^{-5}$ | $1.84 \times 10^{-2}$ |

Table S30: Top ranked 10 drugs in "DSigDB" category of Enrichr for top 200 hub proteins for DIP.

| Term | Overlap | P-value | Adjusted P-value |
|---|---|---|---|
| Vorinostat CTD 00003560 | 54/425 | $1.91 \times 10^{-45}$ | $6.86 \times 10^{-42}$ |
| curcumin CTD 00000663 | 54/528 | $2.35 \times 10^{-40}$ | $4.22 \times 10^{-37}$ |
| doxorubicin CTD 00005874 | 60/750 | $7.40 \times 10^{-39}$ | $8.84 \times 10^{-36}$ |
| MG-132 CTD 00002789 | 36/173 | $4.04 \times 10^{-38}$ | $3.62 \times 10^{-35}$ |
| aspirin CTD 00005447 | 53/561 | $8.75 \times 10^{-38}$ | $6.27 \times 10^{-35}$ |
| simvastatin CTD 00007319 | 42/304 | $1.16 \times 10^{-36}$ | $6.94 \times 10^{-34}$ |
| N-Acetyl-L-cysteine CTD 00005305 | 42/315 | $5.32 \times 10^{-36}$ | $2.72 \times 10^{-33}$ |
| Capsaicin CTD 00005570 | 36/206 | $3.29 \times 10^{-35}$ | $1.47 \times 10^{-32}$ |
| tamoxifen CTD 00006827 | 58/802 | $4.56 \times 10^{-35}$ | $1.82 \times 10^{-32}$ |
| Arsenenous acid CTD 00000922 | 70/1283 | $5.54 \times 10^{-35}$ | $1.99 \times 10^{-32}$ |

Table S31: Top ranked 10 drugs in "DrugMatrix" category of Enrichr for top 200 hub proteins for DIP.

| Term | Overlap | P-value | Adjusted P-value |
|---|---|---|---|
| Thioguanine-24 mg/kg in Corn Oil-Rat-Spleen-5d-up | 12/315 | $5.31 \times 10^{-5}$ | $1.68 \times 10^{-1}$ |
| Catechol-40 mg/kg in Saline-Rat-Bone marrow-3d-up | 11/284 | $9.42 \times 10^{-5}$ | $1.68 \times 10^{-1}$ |
| Catechol-195 mg/kg in Saline-Rat-Bone marrow-3d-up | 12/337 | $1.01 \times 10^{-4}$ | $1.68 \times 10^{-1}$ |
| NN-Dimethylformamide-140 mg/kg in Saline-Rat-Spleen-0.25d-up | 11/288 | $1.07 \times 10^{-4}$ | $1.68 \times 10^{-1}$ |
| Methyl Methanesulfonate-113 mg/kg in Water-Rat-Bone marrow-5d-dn | 11/295 | $1.32 \times 10^{-4}$ | $1.68 \times 10^{-1}$ |
| NN-Dimethylformamide-140 mg/kg in Saline-Rat-Spleen-1d-up | 11/300 | $1.52 \times 10^{-4}$ | $1.68 \times 10^{-1}$ |
| NN-Dimethylformamide-1400 mg/kg in Saline-Rat-Spleen-3d-up | 11/302 | $1.62 \times 10^{-4}$ | $1.68 \times 10^{-1}$ |
| Doxorubicin-3 mg/kg in Saline-Rat-Spleen-3d-up | 11/305 | $1.76 \times 10^{-4}$ | $1.68 \times 10^{-1}$ |
| Gentamicin-40 mg/kg in Saline-Rat-Kidney-5d-up | 11/319 | $2.59 \times 10^{-4}$ | $2.19 \times 10^{-1}$ |
| Carboplatin-6 mg/kg in Saline-Rat-Spleen-1d-up | 10/283 | $4.07 \times 10^{-4}$ | $2.85 \times 10^{-1}$ |

Table S32: Top ranked 10 drugs in "Drug Perturbations from GEO down" category of Enrichr for top 200 hub proteins for DIP.

| Term | Overlap | P-value | Adjusted P-value |
|---|---|---|---|
| 5-Fluorouracil 3385 mouse GSE1559 sample 3639 | 19/398 | $9.70 \times 10^{-9}$ | $8.40 \times 10^{-6}$ |
| doxycycline DB00254 human GSE2624 sample 3077 | 18/391 | $4.22 \times 10^{-8}$ | $1.57 \times 10^{-5}$ |
| 5-Fluorouracil 3385 mouse GSE1559 sample 3640 | 19/444 | $5.54 \times 10^{-8}$ | $1.57 \times 10^{-5}$ |
| Bisphenol A 6623 human GSE17624 sample 2654 | 15/281 | $8.85 \times 10^{-8}$ | $1.57 \times 10^{-5}$ |
| actinomycin D 2019 human GSE6400 sample 3098 | 16/323 | $9.09 \times 10^{-8}$ | $1.57 \times 10^{-5}$ |
| 5-Fluorouracil 3385 mouse GSE1559 sample 3635 | 15/302 | $2.25 \times 10^{-7}$ | $3.25 \times 10^{-5}$ |
| pioglitazone 4829 mouse GSE1458 sample 2587 | 14/282 | $5.81 \times 10^{-7}$ | $7.18 \times 10^{-5}$ |
| vitamin a DB00162 rat GSE284 sample 2839 | 16/381 | $8.36 \times 10^{-7}$ | $9.05 \times 10^{-5}$ |
| troglitazone 5591 mouse GSE1458 sample 2589 | 14/295 | $9.94 \times 10^{-7}$ | $9.56 \times 10^{-5}$ |
| mycophenolic acid DB01024 human GSE14630 sample 3302 | 13/261 | $1.43 \times 10^{-6}$ | $1.24 \times 10^{-4}$ |

Table S33: Top ranked 10 drugs in "Drug Perturbations from GEO up" category of Enrichr for top 200 hub proteins for DIP.

| Term | Overlap | P-value | Adjusted P-value |
|---|---|---|---|
| cyclophosphamide 2907 mouse GSE2254 sample 3628 | 19/344 | $8.86 \times 10^{-10}$ | $7.71 \times 10^{-7}$ |
| neocarzinostatin 5282473 human GSE1676 sample 3113 | 17/299 | $4.62 \times 10^{-9}$ | $2.01 \times 10^{-6}$ |
| Insulin 3989 human GSE22309 sample 3370 | 16/308 | $4.72 \times 10^{-8}$ | $1.15 \times 10^{-5}$ |
| Insulin 3989 human GSE22309 sample 3368 | 18/397 | $5.31 \times 10^{-8}$ | $1.15 \times 10^{-5}$ |
| doxycycline DB00254 human GSE2624 sample 3075 | 14/242 | $9.00 \times 10^{-8}$ | $1.57 \times 10^{-5}$ |
| phenol 996 human GSE6907 sample 3528 | 16/336 | $1.56 \times 10^{-7}$ | $2.21 \times 10^{-5}$ |
| cyclophosphamide 2907 mouse GSE2254 sample 3627 | 17/384 | $1.78 \times 10^{-7}$ | $2.21 \times 10^{-5}$ |
| Promyelocytic leukemia DB00755 human GSE5007 sample 2461 | 15/309 | $3.02 \times 10^{-7}$ | $3.28 \times 10^{-5}$ |
| estradiol 5757 mouse GSE2195 sample 3619 | 14/271 | $3.60 \times 10^{-7}$ | $3.48 \times 10^{-5}$ |
| methylprednisolone 6741 rat GSE490 sample 3659 | 14/284 | $6.32 \times 10^{-7}$ | $5.50 \times 10^{-5}$ |

Table S34: Top ranked 10 diseases in "Jensen DISEASES" category of Enrichr for top 50 hub proteins for DIP.

| Term | Overlap | P-value | Adjusted P-value |
|---|---|---|---|
| Cancer | 11/300 | $1.28 \times 10^{-9}$ | $2.85 \times 10^{-7}$ |
| Toxic encephalopathy | 8/126 | $3.86 \times 10^{-9}$ | $4.28 \times 10^{-7}$ |
| DOID:9917 | 5/30 | $2.76 \times 10^{-8}$ | $2.04 \times 10^{-6}$ |
| Thyroid cancer | 4/18 | $2.17 \times 10^{-7}$ | $1.20 \times 10^{-5}$ |
| Sporotrichosis | 3/6 | $5.10 \times 10^{-7}$ | $2.26 \times 10^{-5}$ |
| Esophageal carcinoma | 3/7 | $8.91 \times 10^{-7}$ | $3.30 \times 10^{-5}$ |
| Intestinal benign neoplasm | 3/11 | $4.16 \times 10^{-6}$ | $1.32 \times 10^{-4}$ |
| Li-Fraumeni syndrome | 3/12 | $5.54 \times 10^{-6}$ | $1.54 \times 10^{-4}$ |
| Upper respiratory tract disease | 3/13 | $7.19 \times 10^{-6}$ | $1.60 \times 10^{-4}$ |
| Pseudoachondroplasia | 3/13 | $7.19 \times 10^{-6}$ | $1.60 \times 10^{-4}$ |

Table S35: Top ranked 10 diseases in "OMIM Diseases" category of Enrichr for top 50 hub proteins for DIP.

| Term | Overlap | P-value | Adjusted P-value |
|---|---|---|---|
| colorectal cancer | 4/40 | $6.17 \times 10^{-6}$ | $7.19 \times 10^{-5}$ |
| ovarian cancer | 3/13 | $7.19 \times 10^{-6}$ | $7.19 \times 10^{-5}$ |
| breast cancer | 3/28 | $7.97 \times 10^{-5}$ | $5.31 \times 10^{-4}$ |
| migraine | 2/16 | $1.03 \times 10^{-3}$ | $5.17 \times 10^{-3}$ |
| orofacial cleft | 2/19 | $1.46 \times 10^{-3}$ | $5.86 \times 10^{-3}$ |
| lung cancer | 2/21 | $1.79 \times 10^{-3}$ | $5.97 \times 10^{-3}$ |
| ectodermal dysplasia | 1/11 | $3.25 \times 10^{-2}$ | $7.87 \times 10^{-2}$ |
| pancreatic cancer | 1/11 | $3.25 \times 10^{-2}$ | $7.87 \times 10^{-2}$ |
| dementia | 1/12 | $3.54 \times 10^{-2}$ | $7.87 \times 10^{-2}$ |
| malaria | 1/14 | $4.12 \times 10^{-2}$ | $8.24 \times 10^{-2}$ |

Table S36: Top ranked 10 drugs in "LINCS L1000 Chem Pert Consensus Sigs" category of Enrichr for top 50 hub proteins for DIP.

| Term | Overlap | P-value | Adjusted P-value |
|---|---|---|---|
| CGP-60474 Down | 7/249 | $9.38 \times 10^{-6}$ | $7.15 \times 10^{-2}$ |
| PFI-1 Up | 6/237 | $7.62 \times 10^{-5}$ | $8.28 \times 10^{-2}$ |
| Puromycin Up | 6/239 | $7.98 \times 10^{-5}$ | $8.28 \times 10^{-2}$ |
| Sparfosic-Acid Up | 6/240 | $8.17 \times 10^{-5}$ | $8.28 \times 10^{-2}$ |
| ZG-10 Down | 6/243 | $8.75 \times 10^{-5}$ | $8.28 \times 10^{-2}$ |
| XL-888 Up | 6/245 | $9.15 \times 10^{-5}$ | $8.28 \times 10^{-2}$ |
| MW-STK33-23 Down | 6/245 | $9.15 \times 10^{-5}$ | $8.28 \times 10^{-2}$ |
| Abamectin Up | 6/246 | $9.36 \times 10^{-5}$ | $8.28 \times 10^{-2}$ |
| Dactinomycin Down | 6/248 | $9.78 \times 10^{-5}$ | $8.28 \times 10^{-2}$ |
| I-BET-762 Up | 5/237 | $7.20 \times 10^{-4}$ | $9.13 \times 10^{-2}$ |

Table S37: Top ranked 10 drugs in "DSigDB" category of Enrichr for top 50 hub proteins for DIP.

| Term | Overlap | P-value | Adjusted P-value |
|---|---|---|---|
| Vorinostat CTD 00003560 | 22/425 | $6.25 \times 10^{-22}$ | $1.85 \times 10^{-18}$ |
| rapamycin CTD 00007350 | 18/247 | $1.39 \times 10^{-20}$ | $1.68 \times 10^{-17}$ |
| simvastatin CTD 00007319 | 19/304 | $1.89 \times 10^{-20}$ | $1.68 \times 10^{-17}$ |
| doxorubicin CTD 00005874 | 25/750 | $2.27 \times 10^{-20}$ | $1.68 \times 10^{-17}$ |
| wortmannin CTD 00000504 | 15/146 | $1.71 \times 10^{-19}$ | $1.01 \times 10^{-16}$ |
| aspirin CTD 00005447 | 22/561 | $2.50 \times 10^{-19}$ | $1.23 \times 10^{-16}$ |
| Arsenenous acid CTD 00000922 | 29/1283 | $3.12 \times 10^{-19}$ | $1.32 \times 10^{-16}$ |
| paclitaxel CTD 00007144 | 21/493 | $3.64 \times 10^{-19}$ | $1.34 \times 10^{-16}$ |
| curcumin CTD 00000663 | 20/528 | $2.98 \times 10^{-17}$ | $9.79 \times 10^{-15}$ |
| lovastatin CTD 00006225 | 12/97 | $9.68 \times 10^{-17}$ | $2.86 \times 10^{-14}$ |

Table S38: Top ranked 10 drugs in "DrugMatrix" category of Enrichr for top 50 hub proteins for DIP.

| Term | Overlap | P-value | Adjusted P-value |
|---|---|---|---|
| Catechol-195 mg/kg in Saline-Rat-Bone marrow-3d-up | 8/337 | $7.14 \times 10^{-6}$ | $3.76 \times 10^{-2}$ |
| Bisphenol A-610 mg/kg in Corn Oil-Rat-Spleen-3d-dn | 7/324 | $5.08 \times 10^{-5}$ | $1.34 \times 10^{-1}$ |
| NN-Dimethylformamide-1400 mg/kg in Saline-Rat-Bone marrow-3d-up | 7/379 | $1.35 \times 10^{-4}$ | $1.56 \times 10^{-1}$ |
| Catechol-40 mg/kg in Saline-Rat-Bone marrow-3d-up | 6/284 | $2.05 \times 10^{-4}$ | $1.56 \times 10^{-1}$ |
| Chlorambucil-4.5 mg/kg in Corn Oil-Rat-Bone marrow-1d-dn | 6/285 | $2.09 \times 10^{-4}$ | $1.56 \times 10^{-1}$ |
| Chlorambucil-4.5 mg/kg in Corn Oil-Rat-Bone marrow-3d-dn | 6/294 | $2.47 \times 10^{-4}$ | $1.56 \times 10^{-1}$ |
| Clobetasol Propionate-17 mg/kg in Corn Oil-Rat-Bone marrow-1d-dn | 6/299 | $2.70 \times 10^{-4}$ | $1.56 \times 10^{-1}$ |
| Betamethasone-79 mg/kg in Corn Oil-Rat-Bone marrow-3d-dn | 6/302 | $2.85 \times 10^{-4}$ | $1.56 \times 10^{-1}$ |
| Hydrocortisone-56 mg/kg in Corn Oil-Rat-Bone marrow-3d-dn | 6/307 | $3.11 \times 10^{-4}$ | $1.56 \times 10^{-1}$ |
| Melphalan-12 mg/kg in CMC-Rat-Bone marrow-5d-dn | 6/307 | $3.11 \times 10^{-4}$ | $1.56 \times 10^{-1}$ |

Table S39: Top ranked 10 drugs in "Drug Perturbations from GEO down" category of Enrichr for top 50 hub proteins for DIP.

| Term | Overlap | P-value | Adjusted P-value |
|---|---|---|---|
| Bisphenol A 6623 human GSE17624 sample 2654 | 10/281 | $1.04 \times 10^{-8}$ | $7.15 \times 10^{-6}$ |
| Etanercept DB00005 human GSE7524 sample 3295 | 10/379 | $1.72 \times 10^{-7}$ | $5.93 \times 10^{-5}$ |
| IFN-gamma1b DB00033 human GSE5542 sample 2476 | 8/266 | $1.24 \times 10^{-6}$ | $2.85 \times 10^{-4}$ |
| vitamin a DB00162 rat GSE284 sample 2839 | 9/381 | $1.89 \times 10^{-6}$ | $3.12 \times 10^{-4}$ |
| doxycycline DB00254 human GSE2624 sample 3077 | 9/391 | $2.33 \times 10^{-6}$ | $3.12 \times 10^{-4}$ |
| neocarzinostatin 5282473 human GSE1676 sample 3113 | 8/301 | $3.11 \times 10^{-6}$ | $3.12 \times 10^{-4}$ |
| phenytoin 1775 rat GSE2880 sample 3033 | 7/211 | $3.17 \times 10^{-6}$ | $3.12 \times 10^{-4}$ |
| methylprednisolone 6741 rat GSE490 sample 3671 | 9/425 | $4.60 \times 10^{-6}$ | $3.52 \times 10^{-4}$ |
| doxycycline DB00254 human GSE2624 sample 3074 | 9/425 | $4.60 \times 10^{-6}$ | $3.52 \times 10^{-4}$ |
| 5-Fluorouracil 3385 mouse GSE1559 sample 3640 | 9/444 | $6.55 \times 10^{-6}$ | $4.26 \times 10^{-4}$ |

Table S40: Top ranked 10 drugs in "Drug Perturbations from GEO up" category of Enrichr for top 200 hub proteins.

| Term | Overlap | P-value | Adjusted P-value |
|---|---|---|---|
| methylprednisolone 6741 rat GSE490 sample 3658 | 9/293 | $2.11 \times 10^{-7}$ | $8.66 \times 10^{-5}$ |
| neocarzinostatin 5282473 human GSE1676 sample 3113 | 9/299 | $2.51 \times 10^{-7}$ | $8.66 \times 10^{-5}$ |
| estradiol 5757 mouse GSE2195 sample 3619 | 8/271 | $1.43 \times 10^{-6}$ | $2.33 \times 10^{-4}$ |
| cisplatin DB00515 mouse GSE6206 sample 3412 | 8/279 | $1.77 \times 10^{-6}$ | $2.33 \times 10^{-4}$ |
| decitabine DB01262 human GSE29077 sample 2539 | 8/279 | $1.77 \times 10^{-6}$ | $2.33 \times 10^{-4}$ |
| methylprednisolone 6741 rat GSE490 sample 3659 | 8/284 | $2.02 \times 10^{-6}$ | $2.33 \times 10^{-4}$ |
| MT19c compound SID 134222379 human GSE23616 sample 3352 | 8/291 | $2.42 \times 10^{-6}$ | $2.39 \times 10^{-4}$ |
| apratoxin A 6326668 human GSE2742 sample 3071 | 7/211 | $3.17 \times 10^{-6}$ | $2.73 \times 10^{-4}$ |
| isoflurane 3763 rat GSE6433 sample 3654 | 8/316 | $4.45 \times 10^{-6}$ | $3.42 \times 10^{-4}$ |
| quercetin 5280343 human GSE7259 sample 3415 | 8/336 | $6.98 \times 10^{-6}$ | $4.33 \times 10^{-4}$ |

Table S41: Top 10 diseases in the "Jensen Diseases" category of Enrichr for 502 gene names selected by $u_{3i}$ for BioGRID.

| Term | Overlap | P-value | Adjusted P-value |
|---|---|---|---|
| Diamond-Blackfan anemia | 12/15 | $2.39 \times 10^{-17}$ | $1.35 \times 10^{-14}$ |
| Bowen-Conradi syndrome | 6/15 | $1.01 \times 10^{-6}$ | $2.86 \times 10^{-4}$ |
| Spondylolysis | 4/8 | $2.55 \times 10^{-5}$ | $4.81 \times 10^{-3}$ |
| Frontotemporal dementia | 7/39 | $4.67 \times 10^{-5}$ | $6.60 \times 10^{-3}$ |
| Paronychia | 3/6 | $2.99 \times 10^{-4}$ | $3.05 \times 10^{-2}$ |
| Stomach cancer | 4/14 | $3.24 \times 10^{-4}$ | $3.05 \times 10^{-2}$ |
| Dent disease | 4/17 | $7.25 \times 10^{-4}$ | $5.85 \times 10^{-2}$ |
| Vaccinia | 5/32 | $1.13 \times 10^{-3}$ | $6.01 \times 10^{-2}$ |
| Spinal muscular atrophy | 4/19 | $1.13 \times 10^{-3}$ | $6.01 \times 10^{-2}$ |
| Spinocerebellar ataxia type 2 | 4/19 | $1.13 \times 10^{-3}$ | $6.01 \times 10^{-2}$ |

Table S42: Top 10 diseases in the "OMIM Diseases" category of Enrichr for gene names selected by $u_{3i}$ for BioGRID.

| Term | Overlap | P-value | Adjusted P-value |
|---|---|---|---|
| thyroid carcinoma | 4/17 | $7.25 \times 10^{-4}$ | $1.52 \times 10^{-2}$ |
| anemia | 7/61 | $8.24 \times 10^{-4}$ | $1.52 \times 10^{-2}$ |
| dementia | 3/12 | $2.94 \times 10^{-3}$ | $3.62 \times 10^{-2}$ |
| lateral sclerosis | 3/19 | $1.14 \times 10^{-2}$ | $1.05 \times 10^{-1}$ |
| gastric cancer | 2/11 | $2.99 \times 10^{-2}$ | $2.21 \times 10^{-1}$ |
| ovarian cancer | 2/13 | $4.10 \times 10^{-2}$ | $2.53 \times 10^{-1}$ |
| diabetes mellitus, type 2 | 2/32 | $1.92 \times 10^{-1}$ | $7.17 \times 10^{-1}$ |
| alopecia | 1/11 | $2.44 \times 10^{-1}$ | $7.17 \times 10^{-1}$ |
| psoriasis | 1/12 | $2.63 \times 10^{-1}$ | $7.17 \times 10^{-1}$ |
| arrhythmogenic right ventricular dysplasia | 1/13 | $2.82 \times 10^{-1}$ | $7.17 \times 10^{-1}$ |

Table S43: Top 10 diseases in the "Jensen Diseases" category of Enrichr for 41 gene names selected by $u_{6i}$ for DIP.

| Term | Overlap | P-value | Adjusted P-value |
|---|---|---|---|
| Histoplasmosis | 3/17 | $4.57 \times 10^{-6}$ | $6.86 \times 10^{-4}$ |
| Hypohidrotic ectodermal dysplasia | 2/12 | $2.42 \times 10^{-4}$ | $1.81 \times 10^{-2}$ |
| Arthritis | 4/186 | $4.62 \times 10^{-4}$ | $2.31 \times 10^{-2}$ |
| Progressive supranuclear palsy | 2/24 | $9.95 \times 10^{-4}$ | $3.73 \times 10^{-2}$ |
| Lung disease | 3/119 | $1.61 \times 10^{-3}$ | $3.80 \times 10^{-2}$ |
| Wolf-Hirschhorn syndrome | 2/31 | $1.66 \times 10^{-3}$ | $3.80 \times 10^{-2}$ |
| Chagas disease | 2/32 | $1.77 \times 10^{-3}$ | $3.80 \times 10^{-2}$ |
| Exanthem | 2/36 | $2.24 \times 10^{-3}$ | $4.20 \times 10^{-2}$ |
| Cancer | 4/300 | $2.70 \times 10^{-3}$ | $4.50 \times 10^{-2}$ |
| Gastritis | 2/43 | $3.18 \times 10^{-3}$ | $4.54 \times 10^{-2}$ |

Table S44: Top 10 diseases in the "OMIM Diseases" category of Enrichr for gene names selected by $u_{6i}$ for DIP.

| Term | Overlap | P-value | Adjusted P-value |
|---|---|---|---|
| ectodermal dysplasia | 2/11 | $2.02 \times 10^{-4}$ | $2.22 \times 10^{-3}$ |
| immunodeficiency | 2/28 | $1.36 \times 10^{-3}$ | $7.46 \times 10^{-3}$ |
| pancreatic cancer | 1/11 | $2.12 \times 10^{-2}$ | $4.23 \times 10^{-2}$ |
| dementia | 1/12 | $2.32 \times 10^{-2}$ | $4.23 \times 10^{-2}$ |
| melanoma | 1/13 | $2.51 \times 10^{-2}$ | $4.23 \times 10^{-2}$ |
| adenoma | 1/14 | $2.70 \times 10^{-2}$ | $4.23 \times 10^{-2}$ |
| malaria | 1/14 | $2.70 \times 10^{-2}$ | $4.23 \times 10^{-2}$ |
| migraine | 1/16 | $3.08 \times 10^{-2}$ | $4.23 \times 10^{-2}$ |
| orofacial cleft | 1/19 | $3.64 \times 10^{-2}$ | $4.39 \times 10^{-2}$ |
| lung cancer | 1/21 | $4.02 \times 10^{-2}$ | $4.39 \times 10^{-2}$ |

Table S45: Top ranked 10 drugs in "LINCS L1000 Chem Pert Consensus Sigs" category of Enrichr for gene names selected by $u_{3i}$ for BioGRID.

| Term | Overlap | P-value | Adjusted P-value |
|---|---|---|---|
| PP-30 Down | 32/244 | $2.03 \times 10^{-14}$ | $2.20 \times 10^{-10}$ |
| CHIR-99021 Up | 29/244 | $4.27 \times 10^{-12}$ | $2.31 \times 10^{-8}$ |
| ABT-737 Up | 28/235 | $9.48 \times 10^{-12}$ | $2.85 \times 10^{-8}$ |
| Caffeic-Acid Up | 28/236 | $1.05 \times 10^{-11}$ | $2.85 \times 10^{-8}$ |
| Thapsigargin Up | 27/236 | $5.76 \times 10^{-11}$ | $1.25 \times 10^{-7}$ |
| Brefeldin-A Up | 26/236 | $3.02 \times 10^{-10}$ | $5.46 \times 10^{-7}$ |
| Ruxolitinib Up | 25/248 | $4.24 \times 10^{-9}$ | $6.56 \times 10^{-6}$ |
| BIBU-1361 Down | 24/246 | $1.65 \times 10^{-8}$ | $2.09 \times 10^{-5}$ |
| Pridinol Up | 24/247 | $1.78 \times 10^{-8}$ | $2.09 \times 10^{-5}$ |
| Arbutin Up | 24/248 | $1.93 \times 10^{-8}$ | $2.09 \times 10^{-5}$ |

Table S46: Top ranked 10 drugs in "DSigDB" category of Enrichr for gene names selected by $u_{3i}$ for BioGRID.

| Term | Overlap | P-value | Adjusted P-value |
|---|---|---|---|
| verteporfin HL60 DOWN | 45/283 | $2.84 \times 10^{-23}$ | $6.38 \times 10^{-20}$ |
| clindamycin HL60 DOWN | 105/1626 | $1.03 \times 10^{-19}$ | $1.15 \times 10^{-16}$ |
| captopril PC3 DOWN | 70/856 | $2.05 \times 10^{-18}$ | $1.54 \times 10^{-15}$ |
| lobeline HL60 DOWN | 93/1510 | $4.11 \times 10^{-16}$ | $2.31 \times 10^{-13}$ |
| neostigmine bromide PC3 DOWN | 55/650 | $3.02 \times 10^{-15}$ | $1.36 \times 10^{-12}$ |
| verteporfin MCF7 DOWN | 32/235 | $6.90 \times 10^{-15}$ | $2.58 \times 10^{-12}$ |
| glibenclamide HL60 DOWN | 83/1333 | $1.16 \times 10^{-14}$ | $3.73 \times 10^{-12}$ |
| puromycin PC3 DOWN | 57/722 | $1.85 \times 10^{-14}$ | $4.70 \times 10^{-12}$ |
| dexverapamil MCF7 DOWN | 37/326 | $1.88 \times 10^{-14}$ | $4.70 \times 10^{-12}$ |
| etifenin PC3 DOWN | 50/609 | $1.92 \times 10^{-13}$ | $4.31 \times 10^{-11}$ |

Table S47: Top ranked 10 drugs in "DrugMatrix" category of Enrichr for gene names selected by $u_{3i}$ for BioGRID.

| Term | Overlap | P-value | Adjusted P-value |
|---|---|---|---|
| NN-Dimethylformamide-140 mg/kg in Saline-Rat-Spleen-3d-dn | 34/345 | $1.23 \times 10^{-11}$ | $6.09 \times 10^{-8}$ |
| Miconazole-920 mg/kg in Corn Oil-Rat-Liver-5d-up | 31/298 | $2.55 \times 10^{-11}$ | $6.09 \times 10^{-8}$ |
| Allyl Alcohol-30 uM in DMSO-Rat-Primary rat hepatocytes-1d-up | 30/281 | $2.81 \times 10^{-11}$ | $6.09 \times 10^{-8}$ |
| Catechol-40 mg/kg in Saline-Rat-Bone marrow-0.25d-dn | 31/306 | $5.02 \times 10^{-11}$ | $6.09 \times 10^{-8}$ |
| Amoxapine-313 mg/kg in CMC-Rat-Liver-3d-up | 32/325 | $5.13 \times 10^{-11}$ | $6.09 \times 10^{-8}$ |
| Catechol-40 mg/kg in Saline-Rat-Bone marrow-1d-dn | 31/307 | $5.46 \times 10^{-11}$ | $6.09 \times 10^{-8}$ |
| Pravastatin-1200 mg/kg in Corn Oil-Rat-Liver-3d-up | 30/292 | $7.32 \times 10^{-11}$ | $6.09 \times 10^{-8}$ |
| Mitomycin C-0.5 mg/kg in Saline-Rat-Bone marrow-0.25d-dn | 31/311 | $7.59 \times 10^{-11}$ | $6.09 \times 10^{-8}$ |
| Vinblastine-0.3 mg/kg in Saline-Rat-Liver-5d-up | 31/311 | $7.59 \times 10^{-11}$ | $6.09 \times 10^{-8}$ |
| Pyrogallol-1000 mg/kg in Water-Rat-Liver-1d-up | 36/409 | $7.73 \times 10^{-11}$ | $6.09 \times 10^{-8}$ |

Table S48: Top ranked 10 drugs in "Drug Perturbations from GEO down" category of Enrichr for gene names selected by $u_{3i}$ for BioGRID.

| Term | Overlap | P-value | Adjusted P-value |
|---|---|---|---|
| apratoxin A 6326668 human GSE2742 sample 3068 | 61/211 | $1.85 \times 10^{-47}$ | $1.64 \times 10^{-44}$ |
| Promyelocytic leukemia DB00755 human GSE5007 sample 2461 | 67/291 | $4.86 \times 10^{-45}$ | $2.15 \times 10^{-42}$ |
| troglitazone DB00197 rat GSE21329 sample 2832 | 72/355 | $2.36 \times 10^{-44}$ | $6.98 \times 10^{-42}$ |
| pioglitazone DB01132 rat GSE21329 sample 2841 | 65/321 | $5.45 \times 10^{-40}$ | $1.14 \times 10^{-37}$ |
| estradiol 5757 human GSE4668 sample 3063 | 57/233 | $6.43 \times 10^{-40}$ | $1.14 \times 10^{-37}$ |
| Promegestone 36709 human GSE67561 sample 3694 | 64/335 | $9.24 \times 10^{-38}$ | $1.36 \times 10^{-35}$ |
| apratoxin A 6326668 human GSE2742 sample 3070 | 65/354 | $3.04 \times 10^{-37}$ | $3.85 \times 10^{-35}$ |
| adenosine triphosphate 5957 human GSE30903 sample 3219 | 63/341 | $2.87 \times 10^{-36}$ | $3.17 \times 10^{-34}$ |
| motexafin gadolinium (4 h) DB05428 human GSE2189 sample 3125 | 59/302 | $2.10 \times 10^{-35}$ | $2.07 \times 10^{-33}$ |
| tibolone 444008 human GSE12446 sample 3204 | 59/313 | $1.70 \times 10^{-34}$ | $1.51 \times 10^{-32}$ |
| atorvastatin DB01076 human GSE2450 sample 2484 | 53/250 | $9.52 \times 10^{-34}$ | $7.61 \times 10^{-32}$ |

Table S49: Top ranked 10 drugs in “Drug Perturbations from GEO up” category of Enrichr for gene names selected by $u_{3i}$ for BioGRID.

| Term | Overlap | P-value | Adjusted P-value |
|---|---|---|---|
| bexarotene DB00307 human GSE6914 sample 2680 | 59/147 | $1.28 \times 10^{-55}$ | $1.14 \times 10^{-52}$ |
| captopril DB01197 mouse GSE19286 sample 2689 | 54/134 | $4.78 \times 10^{-51}$ | $2.13 \times 10^{-48}$ |
| N-METHYLFORMAMIDE 31254 rat GSE5509 sample 3570 | 71/283 | $1.61 \times 10^{-50}$ | $4.80 \times 10^{-48}$ |
| MK-886 CID 3651377 human GSE3202 sample 3192 | 55/182 | $5.41 \times 10^{-44}$ | $1.21 \times 10^{-41}$ |
| lung cancer DB00928 human GSE29077 sample 2535 | 61/245 | $3.45 \times 10^{-43}$ | $6.16 \times 10^{-41}$ |
| Aplidin 56928089 human GSE5681 sample 2598 | 59/230 | $1.36 \times 10^{-42}$ | $2.02 \times 10^{-40}$ |
| diethylstilbestrol DB00255 rat GSE4028 sample 2716 | 58/226 | $6.87 \times 10^{-42}$ | $8.76 \times 10^{-40}$ |
| phenytoin DB00252 rat GSE2880 sample 3291 | 55/207 | $1.24 \times 10^{-40}$ | $1.39 \times 10^{-38}$ |
| imatinib (glivec) 123596 human GSE12211 sample 2518 | 49/158 | $7.16 \times 10^{-40}$ | $7.10 \times 10^{-38}$ |
| ubiquinol 9962735 mouse GSE15129 sample 3459 | 62/288 | $8.00 \times 10^{-40}$ | $7.15 \times 10^{-38}$ |

Table S50: Top ranked 10 drugs in “LINCS L1000 Chem Pert Consensus Sigs” category of Enrichr for gene names selected by $u_{6i}$ for DIP.

| Term | Overlap | P-value | Adjusted P-value |
|---|---|---|---|
| Anisomycin Up | 8/240 | $1.71 \times 10^{-8}$ | $8.30 \times 10^{-5}$ |
| Anguidine Up | 7/237 | $3.34 \times 10^{-7}$ | $6.24 \times 10^{-4}$ |
| Bruceantin Up | 7/242 | $3.85 \times 10^{-7}$ | $6.24 \times 10^{-4}$ |
| Verrucarin-A Up | 6/238 | $6.26 \times 10^{-6}$ | $5.71 \times 10^{-3}$ |
| Emetine Up | 6/242 | $6.89 \times 10^{-6}$ | $5.71 \times 10^{-3}$ |
| Bufalin Up | 6/243 | $7.05 \times 10^{-6}$ | $5.71 \times 10^{-3}$ |
| Cephaeline Up | 5/236 | $9.10 \times 10^{-5}$ | $5.17 \times 10^{-2}$ |
| Puromycin Up | 5/239 | $9.66 \times 10^{-5}$ | $5.17 \times 10^{-2}$ |
| Sarmentogenin Up | 5/246 | $1.11 \times 10^{-4}$ | $5.17 \times 10^{-2}$ |
| PF-06465469 Down | 5/246 | $1.11 \times 10^{-4}$ | $5.17 \times 10^{-2}$ |

Table S51: Top ranked 10 drugs in “DSigDB” category of Enrichr for gene names selected by $u_{6i}$ for DIP.

| Term | Overlap | P-value | Adjusted P-value |
|---|---|---|---|
| 2,6,9,9-tetramethylcycloundeca-2,6,10-trien-1-one CTD 00003763 | 9/27 | $6.88 \times 10^{-19}$ | $1.23 \times 10^{-15}$ |
| 1'-Acetoxychavicol acetate CTD 00002113 | 9/30 | $2.09 \times 10^{-18}$ | $1.87 \times 10^{-15}$ |
| doxorubicin CTD 00005874 | 16/750 | $2.20 \times 10^{-13}$ | $1.31 \times 10^{-10}$ |
| Vorinostat CTD 00003560 | 13/425 | $7.38 \times 10^{-13}$ | $3.30 \times 10^{-10}$ |
| ZINC SULFIDE CTD 00001487 | 7/51 | $6.60 \times 10^{-12}$ | $1.97 \times 10^{-9}$ |
| parthenolide CTD 00000087 | 7/51 | $6.60 \times 10^{-12}$ | $1.97 \times 10^{-9}$ |
| Anacardic acid C15:3 CTD 00003117 | 6/26 | $8.22 \times 10^{-12}$ | $2.10 \times 10^{-9}$ |
| Evodiamine CTD 00002158 | 6/27 | $1.06 \times 10^{-11}$ | $2.25 \times 10^{-9}$ |
| curcumin CTD 00000663 | 13/528 | $1.13 \times 10^{-11}$ | $2.25 \times 10^{-9}$ |
| carisoprodol BOSS | 9/156 | $1.47 \times 10^{-11}$ | $2.63 \times 10^{-9}$ |

Table S52: Top ranked 10 drugs in “DrugMatrix” category of Enrichr for gene names selected by $u_{6i}$ for DIP.

| Term | Overlap | P-value | Adjusted P-value |
|---|---|---|---|
| Nortriptyline-63 mg/kg in Water-Rat-Heart-5d-up | 3/287 | $1.83 \times 10^{-2}$ | $5.22 \times 10^{-1}$ |
| Doxorubicin-3 mg/kg in Saline-Rat-Spleen-3d-up | 3/305 | $2.14 \times 10^{-2}$ | $5.22 \times 10^{-1}$ |
| N-Nitrosodiethylamine-34 mg/kg in Saline-Rat-Spleen-3d-up | 3/332 | $2.67 \times 10^{-2}$ | $5.22 \times 10^{-1}$ |
| Bromisovalum-250 mg/kg in Corn Oil-Rat-Heart-3d-up | 3/346 | $2.96 \times 10^{-2}$ | $5.22 \times 10^{-1}$ |
| Cetylpyridinium Bromide-16.5 mg/kg in Water-Rat-Liver-0.25d-dn | 2/223 | $7.01 \times 10^{-2}$ | $5.22 \times 10^{-1}$ |
| Lamivudine-35 mg/kg in Water-Rat-Liver-0.25d-dn | 2/240 | $7.96 \times 10^{-2}$ | $5.22 \times 10^{-1}$ |
| Papaverine-69 mg/kg in CMC-Rat-Heart-3d-up | 2/243 | $8.13 \times 10^{-2}$ | $5.22 \times 10^{-1}$ |
| Emetine-1 mg/kg in Saline-Rat-Kidney-1d-up | 2/247 | $8.36 \times 10^{-2}$ | $5.22 \times 10^{-1}$ |
| Simvastatin-15 mg/kg in Corn Oil-Rat-Kidney-3d-up | 2/252 | $8.65 \times 10^{-2}$ | $5.22 \times 10^{-1}$ |
| NN-Dimethylformamide-140 mg/kg in Saline-Rat-Spleen-3d-up | 2/255 | $8.82 \times 10^{-2}$ | $5.22 \times 10^{-1}$ |

Table S53: Top ranked 10 drugs in "Drug Perturbations from GEO down" category of Enrichr for gene names selected by $u_{6i}$ for DIP.

| Term | Overlap | P-value | Adjusted P-value |
|---|---|---|---|
| Etanercept DB00005 human GSE7524 sample 3295 | 9/379 | $3.68 \times 10^{-8}$ | $2.12 \times 10^{-5}$ |
| alitretinoin DB00523 rat GSE3952 sample 2674 | 7/407 | $1.20 \times 10^{-5}$ | $3.48 \times 10^{-3}$ |
| Lapatinib DB01259 human GSE38376 sample 3252 | 6/313 | $2.96 \times 10^{-5}$ | $5.70 \times 10^{-3}$ |
| 4-Hydroxytamoxifen 449459 human GSE26298 sample 3239 | 5/199 | $4.06 \times 10^{-5}$ | $5.86 \times 10^{-3}$ |
| dactinomycin DB00970 mouse GSE5324 sample 2700 | 7/544 | $7.65 \times 10^{-5}$ | $8.85 \times 10^{-3}$ |
| estradiol 5757 human GSE4668 sample 3061 | 5/269 | $1.68 \times 10^{-4}$ | $1.62 \times 10^{-2}$ |
| isoflurane 3763 rat GSE6433 sample 3653 | 5/290 | $2.38 \times 10^{-4}$ | $1.97 \times 10^{-2}$ |
| lapatinib DB01259 human GSE38376 sample 2585 | 5/301 | $2.83 \times 10^{-4}$ | $2.02 \times 10^{-2}$ |
| foretinib 42642645 human GSE16179 sample 2582 | 4/168 | $3.14 \times 10^{-4}$ | $2.02 \times 10^{-2}$ |
| methylprednisolone 6741 rat GSE490 sample 3662 | 5/338 | $4.81 \times 10^{-4}$ | $2.36 \times 10^{-2}$ |

Table S54: Top ranked 10 drugs in "Drug Perturbations from GEO up" category of Enrichr for gene names selected by $u_{6i}$ for DIP.

| Term | Overlap | P-value | Adjusted P-value |
|---|---|---|---|
| doxycycline DB00254 human GSE2624 sample 3076 | 8/272 | $4.51 \times 10^{-8}$ | $2.96 \times 10^{-5}$ |
| soman 7305 rat GSE13428 sample 2638 | 9/441 | $1.34 \times 10^{-7}$ | $4.39 \times 10^{-5}$ |
| Lipopolysaccharide 11970143 human GSE5504 sample 3484 | 8/418 | $1.19 \times 10^{-6}$ | $2.60 \times 10^{-4}$ |
| levetiracetam 5284583 rat GSE2880 sample 2669 | 6/201 | $2.37 \times 10^{-6}$ | $3.27 \times 10^{-4}$ |
| sulforaphane 5350 human GSE28813 sample 3445 | 7/320 | $2.49 \times 10^{-6}$ | $3.27 \times 10^{-4}$ |
| doxycycline DB00254 human GSE2624 sample 3075 | 6/242 | $6.89 \times 10^{-6}$ | $7.53 \times 10^{-4}$ |
| lipopolysaccharide (LPS) 11970143 mouse GSE23639 sample 3443 | 6/261 | $1.06 \times 10^{-5}$ | $9.08 \times 10^{-4}$ |
| 4-Hydroxynonenal 5283344 human GSE2397 sample 3083 | 6/263 | $1.11 \times 10^{-5}$ | $9.08 \times 10^{-4}$ |
| methylprednisolone 6741 rat GSE490 sample 3661 | 6/277 | $1.49 \times 10^{-5}$ | $1.02 \times 10^{-3}$ |
| ubiquinol 9962735 human GSE19627 sample 3449 | 6/282 | $1.64 \times 10^{-5}$ | $1.02 \times 10^{-3}$ |
